# Genomic foundation models and complementary variant-scoring strategies in LGMD-associated genes: architecture, adaptation, and sequence context

**DOI:** 10.64898/2026.11.21.768324

**Authors:** Sarvesh Galgale, Isha Vats, Josna Wilson, Agnik Haldar

## Abstract

Genomic foundation models learn representations from DNA sequence, although their usefulness for variant classification depends on the architecture and downstream evaluation procedure. We compared six genomic foundation models using 6,960 ClinVar single-nucleotide variants from ten limb-girdle muscular dystrophy (LGMD)-associated genes, with 4,866 variants assigned to training and 2,094 to held-out testing. The analyses included frozen alternate-minus-reference embeddings at 101, 201 and 501 bp, supervised adaptation at 101 bp, and gene-held-out downstream classifier training. GPN-Star V100 and AlphaGenome Atlas AVI were evaluated separately using their native scores. At 101 bp, frozen Evo2 achieved the highest discrimination among the six foundation models (MCC 0.644; AUROC 0.879; AUPRC 0.818). Supervised adaptation increased MCC for three architectures and reduced it for the remaining three. Extending sequence context to 501 bp improved all three primary metrics in five of six frozen architectures, with Evo2 reaching an MCC of 0.864, AUROC of 0.964 and AUPRC of 0.952. Evo2 also led the frozen gene-held-out readouts (pooled MCC 0.641; AUROC 0.852; AUPRC 0.789). GPN-Star V100 and AVI achieved MCC values of 0.929 and 0.938, respectively, using distinct biological inputs and scoring procedures. The number of variants misclassified across all evaluated configurations decreased from five under the original 101-bp readouts to one after including longer contexts and native scores. An exploratory HGVS-category control identified substantial annotation-associated structure in the benchmark labels. The findings characterize classification performance against aggregate ClinVar labels within LGMD-associated genes. Disease-specific pathogenicity and clinical diagnostic utility remain untested.

## 1. Introduction

Limb-girdle muscular dystrophies (LGMDs) comprise a heterogeneous group of inherited neuromuscular disorders characterized by progressive weakness of the proximal muscles. Their genetic diversity presents challenges for variant interpretation, particularly when implicated genes are associated with more than one clinical phenotype. A recent ClinGen expert-panel assessment examined 31 genes implicated in LGMD and related disorders and identified differences in the strength of their gene-disease associations (Mohan et al., 2024). Although genomic sequencing can identify variants in these genes, establishing their clinical significance requires consideration of the evidence associated with each variant and its relationship to the relevant disease phenotype.

Genomic foundation models provide a means of extracting representations from DNA sequences through large-scale pretraining. Their architectures and learning objectives differ considerably. Nucleotide Transformer demonstrated the potential of pretrained genomic representations across regulatory and variant-related tasks (Dalla-Torre et al., 2025). DNABERT-2 uses a transformer architecture with byte-pair encoding (Zhou et al., 2024), while HyenaDNA applies implicit convolutions to process long sequence contexts (Nguyen et al., 2023). Caduceus incorporates bidirectional state-space modeling and reverse-complement equivariance (Schiff et al., 2024). Other approaches include modernGENA, which builds on BERT-style genomic representation learning (Aspidova et al., 2026), and Evo2, an autoregressive model trained at nucleotide resolution (Brixi et al., 2026).

Variant classification also depends on how pretrained representations are extracted and evaluated. Pooling procedures and downstream classifiers affect the information available for classification, while supervised adaptation updates pretrained representations using labelled examples. Increasing the surrounding sequence window can also alter these representations, although the effect may differ across architectures. Previous benchmarking has reported differences associated with model architecture, representation extraction and adaptation strategy (Feng et al., 2025). We examined these factors using a common variant dataset to compare their effects on classification performance.

Other computational approaches incorporate biological information beyond representations of local DNA sequence. Evolutionary constraint scoring uses patterns of conservation and substitution across species to estimate the tolerance of genomic positions to variation. GPN-Star derives constraint estimates from multispecies whole-genome alignments and phylogenetic information (Ye et al., 2026). AlphaGenome predicts molecular measurements associated with transcription, chromatin state, regulatory activity and splicing (Avsec et al., 2026). The AlphaGenome Atlas Variant Impact (AVI) score combines AlphaGenome-derived predictions with AlphaMissense, conservation measures and variant-annotation features in a separately trained composite predictor (Cheng et al., 2026). The biological inputs and development procedures underlying these scoring systems differ from those used for classifiers fitted to local sequence embeddings.

Models with similar aggregate discrimination can still misclassify different variants. We therefore examined errors at the variant level, recognizing that agreement between predictions does not establish a shared biological mechanism. Identifying correct predictions retrospectively also requires knowledge of the benchmark labels. Aggregate ClinVar clinical-significance classifications may reflect variant-consequence and annotation-associated information. Because several genes in the panel are associated with multiple phenotypes, these classifications do not necessarily establish pathogenicity specifically attributable to LGMD.

We evaluated six genomic foundation models using 6,960 ClinVar single-nucleotide variants from ten LGMD-associated genes. Frozen alternate-minus-reference embeddings were examined at 101, 201 and 501 bp, alongside supervised adaptation at 101 bp. We additionally assessed downstream classification when labelled variants from each evaluation gene were excluded from classifier training. GPN-Star V100 and AlphaGenome Atlas AVI were evaluated using their native variant-level scores. Variant-level error comparisons and an exploratory HGVS-derived annotation control were included to examine patterns underlying classification performance. Together, these analyses characterize the effects of representation extraction, adaptation, sequence context and scoring procedures within a fixed ClinVar benchmark, with interpretation restricted to the evaluated protocols and clinical-significance labels.

## 2. Methods

### 2.1 Benchmark construction and canonical variant set

We constructed a binary variant-classification benchmark from ClinVar (Landrum et al., 2018), focusing on ten genes associated with limb-girdle muscular dystrophy (LGMD): ANO5, CAPN3, DNAJB6, DYSF, FKRP, LMNA, SGCA, SGCB, SGCG and TCAP. Only single-nucleotide variants mapped to the GRCh38 reference assembly were considered. Class labels were assigned from aggregate ClinVar clinical-significance annotations. Pathogenic, Likely pathogenic and Pathogenic/Likely pathogenic records formed the positive class, while Benign, Likely benign and Benign/Likely benign records formed the negative class. Variants with other clinical-significance classifications were excluded. We did not impose a minimum ClinVar review-status threshold.

Two records with identical reference and alternate alleles were excluded, leaving 6,960 variants across 6,094 distinct genomic loci. The benchmark contained 1,692 positive-class and 5,268 negative-class variants. A fixed partition assigned 4,866 variants to training and 2,094 to held-out testing; the latter comprised 509 positive-class and 1,585 negative-class variants. ClinVar VariationID served as the unique variant identifier. Partitioning was performed at the chromosome-position locus level, so variants occupying the same genomic locus could not occur in both training and test sets. Distinct loci were not required to be separated by a minimum genomic distance. The held-out test set was excluded from classifier fitting, hyperparameter selection and decision-threshold optimization.

Gene membership determined eligibility for the benchmark, rather than a condition-specific ClinVar assertion. Because several of the selected genes are associated with phenotypes beyond LGMD, aggregate clinical-significance labels cannot be assumed to indicate pathogenicity specifically attributable to LGMD. The benchmark therefore addresses ClinVar variant classification within an LGMD-associated gene panel, not the identification of variants independently established to cause LGMD.

### 2.2 Sequence construction and local context windows

Reference and alternate DNA sequences were generated from the NCBI RefSeq GRCh38.p14 assembly (GCF_000001405.40). For each variant, we extracted centered windows of 101, 201 and 501 bp, corresponding to 50, 100 and 250 flanking nucleotides on either side of the variant position. The reference sequence was taken directly from the genome assembly. Its alternate counterpart was constructed by replacing the nucleotide at the central position with the recorded alternate allele. Thus, each reference-alternate pair differed at only one position. All sequences were extracted in reference-genome orientation, without reorientation according to transcript strand.

Matched sequence pairs were generated for all 6,960 variants at each window length. The 101-bp context formed the basis of the primary frozen-representation benchmark and the supervised-adaptation experiments. We used 201- and 501-bp windows in separate frozen-model analyses to examine whether additional flanking sequence altered classification performance. The canonical training and test assignments remained unchanged across context lengths.

### 2.3 Frozen sequence foundation-model benchmark

The frozen-representation benchmark included six pretrained genomic foundation models: DNABERT-2, Nucleotide Transformer V2 (NT-v2), HyenaDNA, Caduceus-Ph, modernGENA and Evo2. Each model processed the 101-bp reference and alternate sequences independently using its corresponding tokenizer and pretrained backbone. Model parameters remained fixed during representation extraction. Checkpoint identifiers, available revisions and model-specific representation-extraction procedures are detailed in Supplementary Methods S1.2.1.

Sequence-level embeddings were derived by pooling model activations, with the representation layer and pooling procedure determined by the architecture. DNABERT-2, NT-v2 and modernGENA used attention-mask-weighted mean pooling of final hidden states, excluding padding positions. For HyenaDNA, activations were averaged across nucleotide positions without additional special tokens. Caduceus-Ph representations were obtained by averaging the mean-pooled embeddings of each sequence and its reverse complement. Evo2 used mean-pooled activations from the intermediate layer blocks.28.mlp.l3 across the 101 nucleotide positions. The alternate and reference embeddings were then combined through subtraction to produce a single difference embedding for each variant:

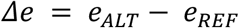

Here, *e_ALT_* and *e_REF_* denote the pooled alternate and reference sequence embeddings, respectively. A related reference-alternate embedding-difference strategy has been used in previous genomic foundation-model benchmarking (Feng et al., 2025).

For each foundation model, we trained a separate logistic-regression classifier using its variant-level difference embeddings. Feature standardization was performed with StandardScaler, followed by L2-regularized logistic regression without class weighting. The regularization parameter was selected from C ∈ {0.001, 0.01, 0.1, 1, 10, 100} using five-fold stratified group cross-validation within the training partition. Groups were defined by chromosome and genomic position to keep variants from identical loci within the same fold. Standardization parameters were estimated independently within each cross-validation training fold. The value of C yielding the highest mean validation Matthews correlation coefficient (MCC), evaluated at a probability threshold of 0.5, was selected.

After hyperparameter selection, the standardization and classification pipeline was fitted using the complete training partition and applied to the fixed held-out test set. Binary predictions were obtained at a probability cutoff of 0.5. Matthews correlation coefficient, area under the receiver operating characteristic curve (AUROC) and area under the precision-recall curve (AUPRC) were used to assess discrimination, with AUROC and AUPRC calculated from predicted class probabilities. The frozen 101-bp workflow is illustrated in Figure 1.

**Figure 1.**
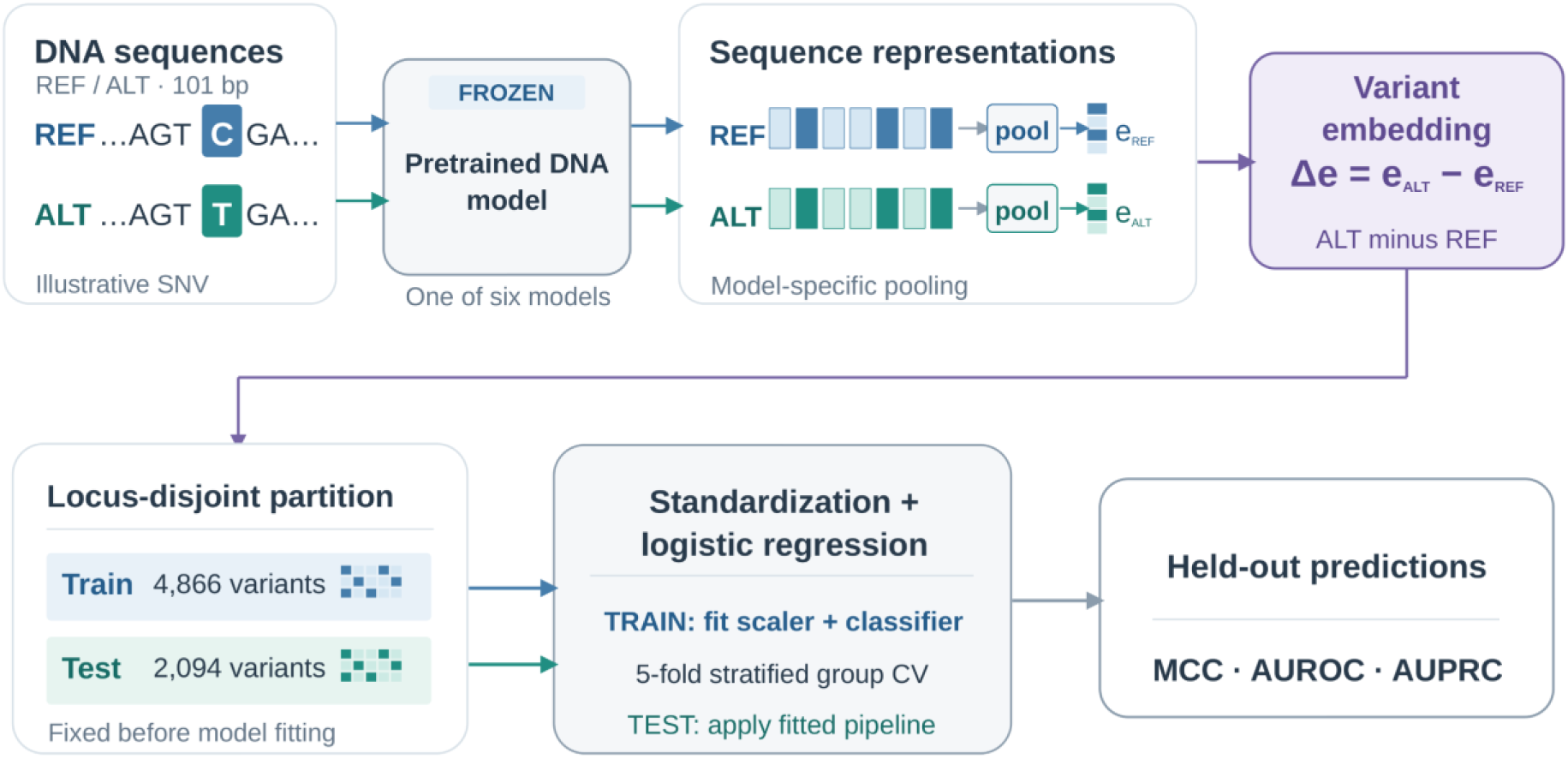
Frozen genomic foundation-model workflow for ClinVar variant classification. Reference and alternate allele-centered DNA sequences were independently processed by the same pretrained genomic foundation model, with model parameters held fixed. Model-specific pooling produced sequence-level representations, which were combined by subtracting the reference embedding from the alternate embedding. The resulting variant-level difference embeddings were used to train L2-regularized logistic-regression classifiers with feature standardization and five-fold stratified group cross-validation on the training partition (n = 4,866). Fitted classifiers were evaluated on the predefined locus-disjoint held-out test partition (n = 2,094) using MCC, AUROC and AUPRC. The diagram illustrates the 101-bp frozen-representation workflow; 201- and 501-bp frozen contexts and supervised adaptation were evaluated separately.

### 2.4 Supervised adaptation of sequence foundation models

Supervised adaptation used the same 101-bp reference and alternate sequences and canonical training and test partitions as the frozen-representation benchmark. For each model, both alleles were processed through a shared backbone using the architecture-specific representation procedures described in Section 2.3. The alternate-minus-reference embedding was passed to a classification head consisting of dropout (0.10) and a linear layer with two output units. Optimization minimized class-weighted cross-entropy loss, with class weights calculated from the optimization subset.

Five architectures, DNABERT-2, NT-v2, HyenaDNA, Caduceus-Ph and modernGENA, underwent full backbone fine-tuning. Evo2 was adapted using low-rank adaptation (LoRA), while its pretrained backbone parameters remained frozen (Hu et al., 2022). Trainable adapters were introduced into the blocks.28.mlp.l1, blocks.28.mlp.l2 and blocks.28.mlp.l3 projections, with rank 8, scaling factor 16 and adapter dropout 0.05. The classification head was optimized jointly with these adapters. In total, the Evo2 configuration contained 370,690 trainable parameters, including 362,496 adapter parameters and 8,194 classification-head parameters.

Hyperparameters and checkpoints were selected using an internal validation subset drawn from the canonical training partition. One fold of a ten-fold stratified group split was reserved for validation, with chromosome-position loci used as groups to prevent identical loci from appearing in both optimization and validation subsets. Backbone learning rates of 1 × 10⁻⁵, 3 × 10⁻⁵ and 1 × 10⁻⁴ were examined for the five fully fine-tuned models. For Evo2, adapter learning rates were 1 × 10⁻⁴, 3 × 10⁻⁴ and 1 × 10⁻³. The classification-head learning rate was fixed at 1 × 10⁻⁴ across configurations.

Training used AdamW with weight decay of 0.01, linear learning-rate decay after a 5% warmup, an effective batch size of 32 variants and gradient clipping at a maximum norm of 1.0. Optimization employed bfloat16 mixed precision and was limited to five epochs per learning-rate configuration. Checkpoint selection was based on validation MCC at a probability threshold of 0.5, with validation AUPRC used to resolve ties. Selected checkpoints were evaluated directly on the fixed test partition, without refitting on the complete training set. Test labels were not used for training, hyperparameter selection or checkpoint selection. The selected learning rates and epochs are reported in Supplementary Methods S1.2.

The frozen and adapted DNABERT-2 configurations used different implementations. Frozen embeddings were extracted through MultiMolecule, whereas supervised adaptation used the original DNABERT-2 implementation. Although the converted checkpoint is documented as reproducing the original model’s intermediate representations, numerical equivalence was not independently verified for the extraction and tokenization settings used here. The frozen-to-adapted performance difference for DNABERT-2 therefore cannot be attributed solely to supervised parameter adaptation.

### 2.5 Effect of sequence context length on frozen representations

To examine the influence of surrounding sequence, we evaluated all six foundation models using centered reference and alternate windows of 101, 201 and 501 bp. For each allele, the central 101-bp sequence remained identical across window lengths, while the longer inputs contained additional flanking nucleotides. This allowed us to examine how frozen representations responded to increased sequence context without changing the variant or its immediate sequence neighborhood.

Model parameters remained fixed throughout the context-length experiments. Reference and alternate embeddings were extracted using the architecture-specific procedures described in Section 2.3 and combined as alternate-minus-reference difference embeddings. For each model and sequence length, a separate L2-regularized logistic-regression classifier was trained on the canonical training partition, with feature standardization and regularization selection performed according to the frozen-readout protocol. All configurations were evaluated on the same fixed test partition.

The original 101-bp results provided the reference condition for comparisons with 201- and 501-bp inputs. MCC, AUROC and AUPRC were examined separately for each architecture. Because classifier fitting and regularization selection were repeated for each context length, these comparisons reflect the performance of context-specific frozen readouts rather than an isolated change in input length with an otherwise identical fitted classifier. No supervised backbone adaptation was performed in these experiments.

### 2.6 GPN-Star evolutionary-constraint scoring

GPN-Star was evaluated as an evolutionary-constraint predictor using native variant-level scores (Ye et al., 2026). Unlike the sequence foundation models evaluated through local DNA embeddings, GPN-Star uses multispecies whole-genome alignments and phylogenetic relationships to estimate sequence constraint. The primary analysis used GPN-Star V100, based on an alignment of 100 vertebrate genomes. GPN-Star M447 and P243, derived from alignments of 447 mammalian and 243 primate genomes, respectively, were included as sensitivity analyses.

For each of the 6,960 variants, we used the sign-reversed mutation-rate-calibrated log-likelihood ratio (-LLR) as the continuous score, with higher values indicating stronger predicted evolutionary constraint. Scores were obtained without further training or adaptation to the benchmark labels.

AUROC and AUPRC were calculated directly from the continuous native scores. For threshold-dependent metrics, including MCC, the operating threshold was chosen by maximizing MCC on the canonical training partition. Ties were resolved first by balanced accuracy and then by selecting the higher threshold. The selected threshold was applied without modification to the held-out test partition. GPN-Star V100 served as the primary evolutionary-constraint comparator, while M447 and P243 were assessed separately to examine sensitivity to the underlying evolutionary alignment. These scores represent predicted sequence constraint and were not interpreted as calibrated probabilities of clinical pathogenicity.

### 2.7 AlphaGenome Atlas variant-impact scoring

AlphaGenome Atlas Variant Impact (AVI) scores were evaluated as an independent approach to variant-impact prediction (Cheng et al., 2026). AVI is a composite predictor that combines ten groups of AlphaGenome-derived molecular-effect predictions with AlphaMissense scores, evolutionary conservation measures, protein-consequence annotations and variant-type indicators. Its training used population allele-frequency categories as proxies for variant impact rather than clinical pathogenicity labels. AVI was therefore assessed separately from the sequence foundation models, whose downstream classifiers were trained on the ClinVar benchmark.

AVI scores were obtained for all 6,960 GRCh38 variants. The raw composite scores, their genome-wide quantiles and 18 SHAP-derived feature-attribution fields were retained. Quantiles were transformed to PHRED-scaled scores as follows:

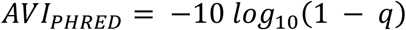

where q represents the genome-wide AVI score quantile. Higher PHRED values indicate greater predicted variant impact. No additional training or recalibration of the native scoring model was performed.

AUROC and AUPRC were calculated from the continuous AVI PHRED scores. For threshold-dependent metrics, the decision threshold was selected by maximizing MCC on the canonical training partition. Ties were resolved using balanced accuracy, followed by selection of the higher threshold. This threshold was then applied unchanged to the fixed test partition. The SHAP-derived attributions were retained as supplementary information but were not used for classifier selection, threshold optimization or the reported discrimination analyses. They were not interpreted as causal evidence of individual molecular contributions. AVI scores were treated as relative measures of predicted variant impact, not calibrated probabilities of clinical pathogenicity.

### 2.8 Performance evaluation and statistical analysis

Classification performance was assessed on the fixed test partition using Matthews correlation coefficient (MCC), area under the receiver operating characteristic curve (AUROC) and area under the precision-recall curve (AUPRC) as the primary metrics (Chicco et al., 2021; Saito & Rehmsmeier, 2015). Precision, sensitivity, specificity, F1 score, balanced accuracy and confusion-matrix counts were also calculated. AUPRC was computed as average precision using the non-interpolated precision-recall formulation. Sequence-model predictions used a probability threshold of 0.5, whereas native-score thresholds were determined from the training partition as described in Sections 2.6 and 2.7.

Uncertainty in the primary frozen and adapted 101-bp evaluations was estimated using 10,000 genomic-locus bootstrap replicates. The fixed test partition contained 1,834 distinct loci.

Sampling was performed with replacement at the chromosome-position level, retaining all variants associated with each sampled locus. This accounted for multiple alternate alleles occurring at the same position. Resampling was not stratified by clinical-significance class because individual loci could contain variants with different labels. Ninety-five percent confidence intervals were calculated using the percentile method. Frozen-to-adapted differences were estimated from paired bootstrap samples to preserve the correspondence between predictions for the same variants.

Three targeted comparisons were examined after reviewing the primary 101-bp results: frozen Evo2 versus adapted NT-v2, adapted Evo2 versus adapted NT-v2, and adapted NT-v2 versus adapted HyenaDNA. Differences in MCC, AUROC and AUPRC were evaluated using two-sided paired random-swap permutation tests, with 10,000 permutations at the genomic-locus level. Holm correction was applied across the nine tests. These comparisons were exploratory rather than prespecified. Their estimates and adjusted p-values are reported in Supplementary Table S2, Panel E. Context-length comparisons were interpreted descriptively, while uncertainty in the separate cross-gene evaluation was assessed using the procedure described in Section 2.10.

A separate paired analysis compared AlphaGenome Atlas AVI and GPN-Star V100 on the same fixed test partition. For each primary metric, 10,000 paired genomic-locus bootstrap replicates were used to estimate 95% percentile confidence intervals for the difference between their scores. Two-sided random-swap permutation tests used 10,000 permutations, with Holm correction across the three metrics. This comparison was treated as a separate statistical family from the nine sequence-model tests (Supplementary Table S4, Panel B).

Probability calibration was examined for the six frozen and six adapted 101-bp sequence-model configurations. Brier score, logarithmic loss and expected calibration error (ECE) were calculated from unadjusted test probabilities, using ten equal-width probability bins for ECE (Guo et al., 2017). Calibration intercepts and slopes were estimated through diagnostic regressions against the test labels and reported only when the corresponding fit was considered successful (Supplementary Table S2, Panels B and F). These regressions were used for evaluation, not for modifying probabilities or selecting models. No test-set recalibration was performed, and the resulting probabilities were not treated as clinically validated pathogenicity estimates.

To examine the influence of variant-annotation structure, we conducted an exploratory control based on six categories assigned through rule-based patterns in ClinVar HGVS descriptions: stop-gained, synonymous, amino-acid substitution, untranslated region, intronic and unclassified. The positive-class frequency of each category was estimated from the training partition and used as a continuous score for variants in the fixed test set. Binary classifications used a threshold of 0.5. We repeated this procedure while excluding the evaluated gene from the training data. Because these categories were inferred from HGVS naming patterns rather than independently validated consequence annotations, the analysis was interpreted as an annotation-based diagnostic rather than a comparison involving equivalent model inputs (Supplementary Table S14).

We also examined discrimination within these HGVS-derived categories using the original frozen and adapted 101-bp predictions and the two primary native scores. Scores and classification thresholds were retained without subgroup-specific optimization. AUROC and AUPRC were calculated where both classes were represented, with comparative interpretation restricted to categories containing at least 20 variants from each class. Exploratory 95% paired genomic-locus bootstrap intervals were estimated using 2,000 replicates within each category (Supplementary Table S15).

### 2.9 Variant-level error overlap across model configurations

Classification errors were compared across 26 configurations evaluated on the same fixed test partition. These included twelve 101-bp sequence-model configurations (six frozen and six adapted), six frozen 201-bp configurations, six frozen 501-bp configurations, GPN-Star V100 and AlphaGenome AVI. The alternative GPN-Star evolutionary alignments were excluded. Predictions were aligned by ClinVar VariationID and evaluated against the corresponding benchmark labels using each configuration’s established classification threshold.

We first identified variants that were repeatedly misclassified by the twelve original 101-bp sequence configurations. A recurrent-error subset comprised 787 variants misclassified by at least four configurations, while five variants were misclassified by all twelve. For these subsets, we examined whether at least one frozen 201-bp model, one frozen 501-bp model or either native scoring approach produced a correct classification.

We additionally calculated the intersection of misclassified variants across three nested sets: the original twelve 101-bp configurations, all 24 sequence-model configurations, and the complete set of 26 configurations. These comparisons characterized how classification errors overlapped as additional approaches were considered.

Because correctness was determined retrospectively using the known test labels, the analysis did not represent a trained ensemble or a prospective prediction strategy. A correct prediction from at least one configuration does not establish that the appropriate configuration could have been selected without knowledge of the true label.

### 2.10 Cross-gene evaluation of frozen representations and native impact scores

We examined cross-gene classification using a leave-one-gene-out downstream training design across the ten LGMD-associated genes. For each evaluation gene, labelled variants from that gene were excluded from classifier development, while the remaining nine genes contributed variants from the canonical training partition. Evaluation was restricted to the excluded gene’s variants in the fixed test set. Together, the ten evaluations covered all 2,094 test variants, with each variant assessed once.

For each of the six genomic foundation models, frozen 101-bp alternate-minus-reference embeddings were used to train an L2-regularized logistic-regression classifier with feature standardization. The regularization parameter was selected from C ∈ {0.001, 0.01, 0.1, 1, 10, 100} using five-fold stratified group cross-validation within the nine-gene training subset. Chromosome-position loci defined the cross-validation groups, and selection was based on mean validation MCC at a probability threshold of 0.5. Ties favored the smaller C value. The selected classifier was then fitted using the complete nine-gene training subset and applied to the excluded gene’s test variants. Foundation-model parameters remained frozen. This procedure yielded 60 gene-specific classifier fits, with their selected regularization parameters reported in Supplementary Table S13.

GPN-Star V100 and AlphaGenome AVI were assessed separately using their existing continuous variant scores, without retraining or recalibration. For each excluded gene, an operating threshold was selected by maximizing MCC on labelled training variants from the other nine genes, with ties resolved in favor of the higher threshold. These gene-specific thresholds were applied to the corresponding test variants. AUROC and AUPRC were calculated from the unchanged continuous scores. The selected native-score thresholds are provided in Supplementary Table S12.

Predictions from the ten gene-specific evaluations were pooled to calculate overall performance. Uncertainty was estimated using 1,000 paired bootstrap replicates, with genomic loci resampled with replacement within each evaluation gene. The same sampled loci were used for paired comparisons, and 95% confidence intervals were obtained using the percentile method. We also assessed sensitivity to individual genes by recalculating pooled performance after excluding each evaluation gene in turn. Gene-level results and exploratory paired comparisons are reported in Supplementary Tables S10 and S11.

These analyses assess downstream classification when labelled training variants from the evaluation gene are unavailable. They do not establish that the evaluated genes were absent from foundation-model pretraining or the development of external scoring systems. The bootstrap intervals are conditional on the observed genes, loci and fitted classifiers; they do not account for classifier refitting or variation across independently sampled genes. Because the comparisons were exploratory and were not adjusted for multiple testing, the intervals were interpreted descriptively rather than as confirmatory evidence of differences between approaches.

## 3. Results

### 3.1 Performance of frozen genomic foundation models

The six frozen genomic foundation models showed considerable variation in classification performance using 101-bp alternate-minus-reference embeddings (Table 1). Evo2 achieved the highest discrimination on the fixed test partition of 2,094 ClinVar variants, with an MCC of 0.644, AUROC of 0.879 and AUPRC of 0.818. The remaining five architectures performed substantially below Evo2. Their MCC values ranged from 0.151 for DNABERT-2 to 0.249 for modernGENA, with AUROC values between 0.598 and 0.663 and AUPRC values between 0.320 and 0.446.

**Table 1.** Primary frozen and adapted 101-bp foundation-model readouts compared with native variant scores.

| Model | Regime | MCC | AUROC | AUPRC | Precision | Recall | Specificity | F1 |
|---|---|---|---|---|---|---|---|---|
| AlphaGenome Atlas AVI | Native score | 0.938 | 0.991 | 0.987 | 0.979 | 0.927 | 0.994 | 0.953 |
| GPN-Star V100 | Native score | 0.929 | 0.990 | 0.983 | 0.958 | 0.935 | 0.987 | 0.946 |
| Evo2 | Frozen + LR | 0.644 | 0.879 | 0.818 | 0.884 | 0.570 | 0.976 | 0.693 |
| Evo2 | Adapted (LoRA) | 0.432 | 0.866 | 0.784 | 0.441 | 0.849 | 0.654 | 0.580 |
| NT-v2 | Adapted (full) | 0.336 | 0.794 | 0.686 | 0.403 | 0.743 | 0.647 | 0.523 |
| HyenaDNA | Adapted (full) | 0.306 | 0.724 | 0.503 | 0.382 | 0.741 | 0.616 | 0.504 |
| modernGENA | Frozen + LR | 0.249 | 0.663 | 0.446 | 0.573 | 0.224 | 0.946 | 0.322 |
| NT-v2 | Frozen + LR | 0.234 | 0.656 | 0.430 | 0.516 | 0.253 | 0.924 | 0.340 |
| Caduceus-Ph | Frozen + LR | 0.223 | 0.656 | 0.423 | 0.545 | 0.204 | 0.945 | 0.297 |
| DNABERT-2 | Adapted (full) | 0.215 | 0.696 | 0.479 | 0.326 | 0.745 | 0.505 | 0.453 |
| HyenaDNA | Frozen + LR | 0.209 | 0.655 | 0.392 | 0.548 | 0.179 | 0.953 | 0.270 |
| Caduceus-Ph | Adapted (full) | 0.160 | 0.630 | 0.360 | 0.306 | 0.682 | 0.504 | 0.423 |
| DNABERT-2 | Frozen + LR | 0.151 | 0.598 | 0.320 | 0.388 | 0.265 | 0.866 | 0.315 |
| modernGENA | Adapted (full) | 0.089 | 0.593 | 0.374 | 0.288 | 0.499 | 0.603 | 0.365 |

Fourteen configurations are included: six frozen and six adapted 101-bp genomic foundation-model readouts, GPN-Star V100 and AlphaGenome Atlas AVI. All metrics were calculated on the fixed test partition of 2,094 variants (509 positive, 1,585 negative). Frozen models used L2-regularized logistic regression on alternate-minus-reference embeddings; adaptation comprised five full fine-tuning regimes and Evo2 LoRA. Sequence-model classifications used a probability threshold of 0.5, while native-score thresholds were selected on the training partition. AUPRC denotes average precision. The native scores use distinct biological inputs and scoring protocols, so between-family comparisons are descriptive.

The differences extended to classification sensitivity and specificity. At the common probability threshold of 0.5, Evo2 correctly identified 290 of the 509 positive-class variants, giving a sensitivity of 0.570. It also correctly classified 1,547 of the 1,585 negative-class variants, corresponding to a specificity of 0.976. Sensitivity among the other five frozen models ranged from 0.179 to 0.265, while specificity ranged from 0.866 to 0.953. Evo2 thus achieved higher discrimination and sensitivity than the other frozen 101-bp readouts while maintaining high specificity. Figure 2 presents these results alongside the adaptation, longer-context and native-score evaluations, covering 26 configurations in total.

**Figure 2.**
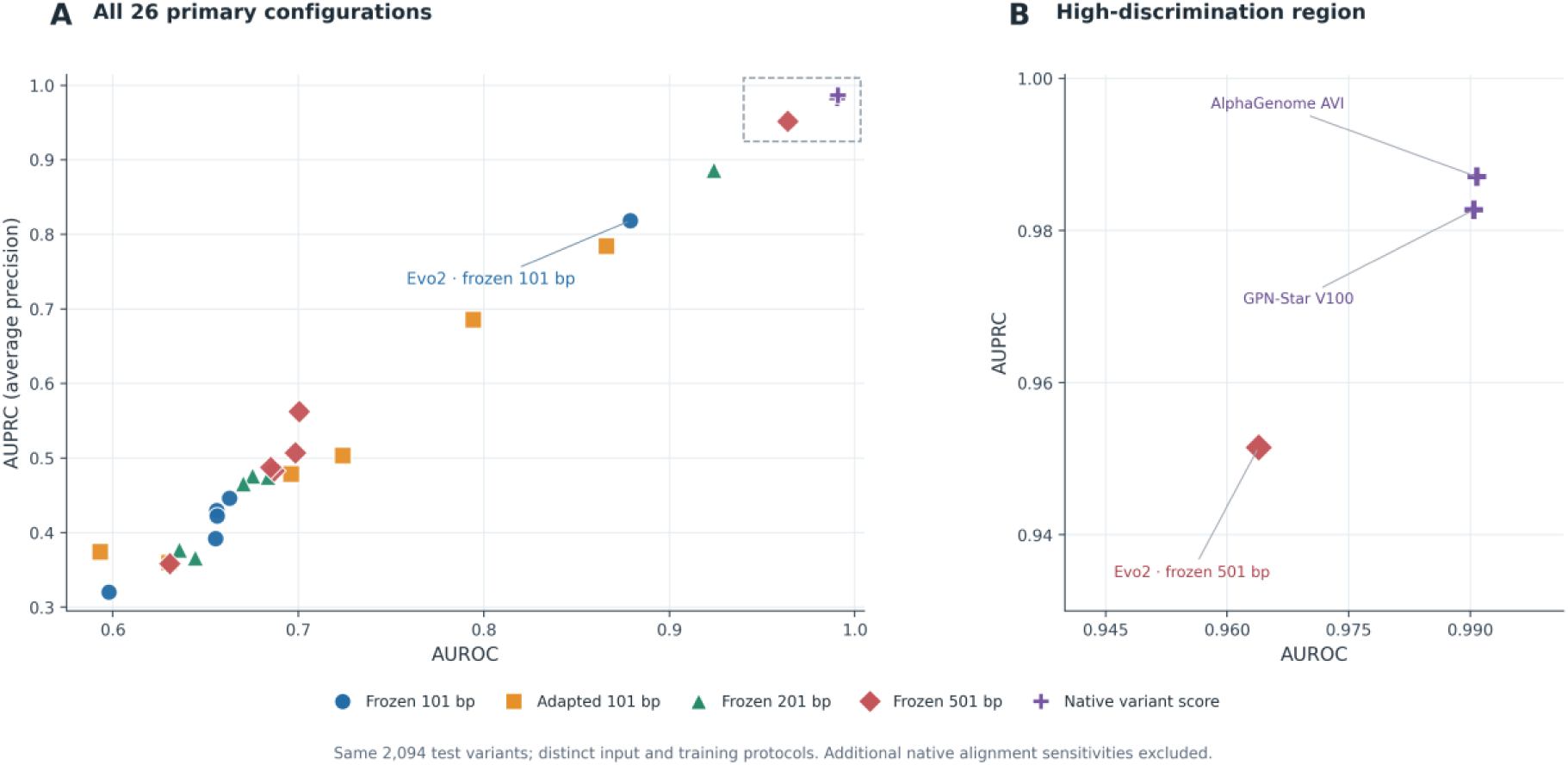
Classification discrimination across 26 genomic foundation-model and native-score configurations. AUROC and AUPRC (average precision) on the fixed test partition of 2,094 ClinVar variants. Panel A shows all 26 primary configurations; panel B enlarges the high-discrimination region containing frozen Evo2 at 501 bp, GPN-Star V100 and AlphaGenome Atlas AVI. The 24 sequence-model configurations comprise six frozen and six adapted 101-bp readouts, together with six frozen readouts each at 201 and 501 bp. Native scores use separately developed evolutionary-constraint or composite variant-impact predictors. The alternative GPN-Star M447 and P243 configurations are excluded and reported in Supplementary Table S4.

### 3.2 Effects of supervised adaptation on classification performance

The effects of supervised adaptation varied across the six architectures (Figure 3; Table 2). Full fine-tuning improved MCC for DNABERT-2 from 0.151 to 0.215, NT-v2 from 0.234 to 0.336, and HyenaDNA from 0.209 to 0.306. All three models also improved in AUROC and AUPRC, with paired genomic-locus bootstrap confidence intervals excluding zero for each metric change (Supplementary Table S2). Among the five fully fine-tuned models, NT-v2 achieved the highest MCC (0.336), AUROC (0.794) and AUPRC (0.686).

**Figure 3.**
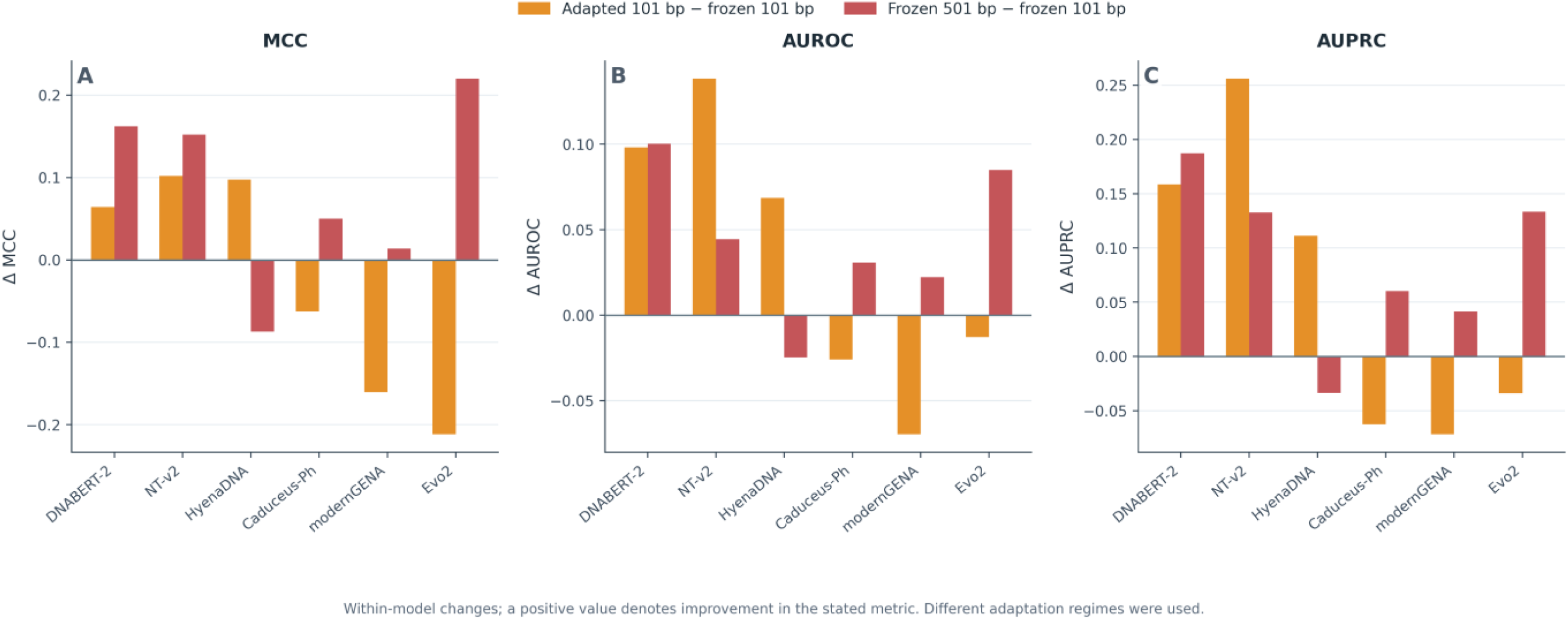
Architecture-dependent changes following supervised adaptation and increased frozen sequence context. Within-model differences in (A) MCC, (B) AUROC and (C) AUPRC on the fixed 2,094-variant test partition. Orange bars represent adapted 101-bp minus frozen 101-bp performance; red bars represent frozen 501-bp minus frozen 101-bp performance. Positive values indicate higher metric values under the comparison condition. Adaptation comprised full backbone fine-tuning for five models and LoRA for Evo2. Context-length comparisons used separately fitted logistic-regression readouts with frozen backbone parameters. The bars show signed differences, not inferential significance; exploratory paired uncertainty estimates for adaptation are reported in Supplementary Table S2.

**Table 2.**
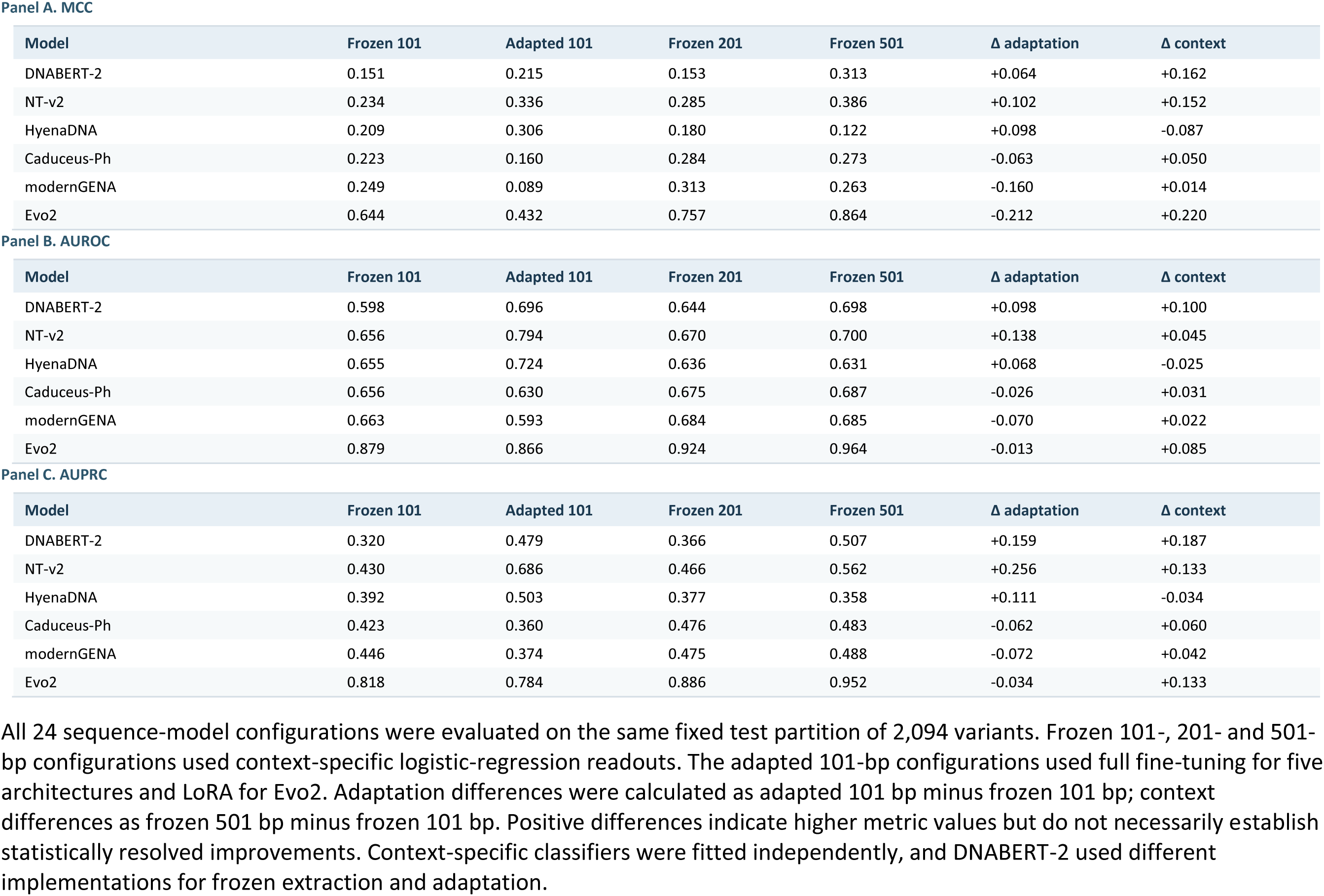
Sequence foundation-model performance under supervised adaptation and frozen sequence-context changes.

| **Panel A. MCC** |  |  |  |  |  |  |
|---|---|---|---|---|---|---|
| **Model** | **Frozen 101** | **Adapted 101** | **Frozen 201** | **Frozen 501** | **Δ adaptation** | **Δ context** |
| DNABERT-2 | 0.151 | 0.215 | 0.153 | 0.313 | +0.064 | +0.162 |
| NT-v2 | 0.234 | 0.336 | 0.285 | 0.386 | +0.102 | +0.152 |
| HyenaDNA | 0.209 | 0.306 | 0.180 | 0.122 | +0.098 | -0.087 |
| Caduceus-Ph | 0.223 | 0.160 | 0.284 | 0.273 | -0.063 | +0.050 |
| modernGENA | 0.249 | 0.089 | 0.313 | 0.263 | -0.160 | +0.014 |
| Evo2 | 0.644 | 0.432 | 0.757 | 0.864 | -0.212 | +0.220 |
| **Panel B. AUROC** |  |  |  |  |  |  |
| **Model** | **Frozen 101** | **Adapted 101** | **Frozen 201** | **Frozen 501** | **Δ adaptation** | **Δ context** |
| DNABERT-2 | 0.598 | 0.696 | 0.644 | 0.698 | +0.098 | +0.100 |
| NT-v2 | 0.656 | 0.794 | 0.670 | 0.700 | +0.138 | +0.045 |
| HyenaDNA | 0.655 | 0.724 | 0.636 | 0.631 | +0.068 | -0.025 |
| Caduceus-Ph | 0.656 | 0.630 | 0.675 | 0.687 | -0.026 | +0.031 |
| modernGENA | 0.663 | 0.593 | 0.684 | 0.685 | -0.070 | +0.022 |
| Evo2 | 0.879 | 0.866 | 0.924 | 0.964 | -0.013 | +0.085 |
| **Panel C. AUPRC** |  |  |  |  |  |  |
| **Model** | **Frozen 101** | **Adapted 101** | **Frozen 201** | **Frozen 501** | **Δ adaptation** | **Δ context** |
| DNABERT-2 | 0.320 | 0.479 | 0.366 | 0.507 | +0.159 | +0.187 |
| NT-v2 | 0.430 | 0.686 | 0.466 | 0.562 | +0.256 | +0.133 |
| HyenaDNA | 0.392 | 0.503 | 0.377 | 0.358 | +0.111 | -0.034 |
| Caduceus-Ph | 0.423 | 0.360 | 0.476 | 0.483 | -0.062 | +0.060 |
| modernGENA | 0.446 | 0.374 | 0.475 | 0.488 | -0.072 | +0.042 |
| Evo2 | 0.818 | 0.784 | 0.886 | 0.952 | -0.034 | +0.133 |

All 24 sequence-model configurations were evaluated on the same fixed test partition of 2,094 variants. Frozen 101-, 201- and 501-bp configurations used context-specific logistic-regression readouts. The adapted 101-bp configurations used full fine-tuning for five architectures and LoRA for Evo2. Adaptation differences were calculated as adapted 101 bp minus frozen 101 bp; context differences as frozen 501 bp minus frozen 101 bp. Positive differences indicate higher metric values but do not necessarily establish statistically resolved improvements. Context-specific classifiers were fitted independently, and DNABERT-2 used different implementations for frozen extraction and adaptation.

The remaining architectures showed less favorable outcomes. For modernGENA, MCC decreased from 0.249 to 0.089, AUROC from 0.663 to 0.593, and AUPRC from 0.446 to 0.374. The bootstrap intervals for all three changes excluded zero. Caduceus-Ph showed a reduction in AUPRC from 0.423 to 0.360, while its MCC and AUROC changes remained uncertain. Evo2 was adapted using LoRA, with MCC declining from 0.644 to 0.432 (ΔMCC = −0.212; 95% CI, −0.255 to −0.168). Its AUPRC also decreased from 0.818 to 0.784. AUROC changed from 0.879 to 0.866, although the confidence interval for this smaller difference included zero. Despite these reductions, Evo2 retained the highest MCC, AUROC and AUPRC among the six adapted configurations.

Three post hoc comparisons were examined using selected 101-bp configurations (Supplementary Table S2, Panel E). Frozen and adapted Evo2 each outperformed adapted NT-v2 across MCC, AUROC and AUPRC, with Holm-adjusted p = 0.00090 for all six comparisons. Adapted NT-v2 also showed higher AUROC and AUPRC than adapted HyenaDNA (Holm-adjusted p = 0.00090 for both). Their MCC difference was less conclusive (ΔMCC = 0.030; 95% CI, −0.016 to 0.077; Holm-adjusted p = 0.20018). These comparisons were selected after examining the primary results and remain exploratory.

Differences in the adaptation procedures also affect the interpretation of these results. Evo2 underwent LoRA adaptation, whereas the other five architectures received full backbone fine-tuning. For DNABERT-2, frozen representations were obtained using MultiMolecule, while supervised adaptation used the original implementation. Run-specific numerical equivalence between the two implementations was not established, introducing additional uncertainty when interpreting the observed improvement after adaptation.

Changes in classification behavior extended beyond the primary discrimination metrics. At the common decision threshold, all six adapted configurations predicted more test variants as positive. Brier scores and expected calibration errors increased for every architecture (Supplementary Table S2; Supplementary Figure S1). For Evo2, the Brier score increased from 0.091 to 0.199, while expected calibration error rose from 0.027 to 0.266. Thus, the improvements in discrimination observed for three architectures occurred alongside poorer raw-probability calibration. Gene-level MCC results are presented in Supplementary Figure S2, although the small positive-class subsets for several genes limit the precision of their individual estimates.

### 3.3 Effect of sequence context length on frozen-model performance

Increasing the sequence window from 101 to 501 bp improved MCC, AUROC and AUPRC in five of the six frozen foundation models (Figure 3; Table 2). Evo2 showed the largest absolute MCC increase, from 0.644 at 101 bp to 0.864 at 501 bp. Its AUROC increased from 0.879 to 0.964, while AUPRC rose from 0.818 to 0.952. The 501-bp Evo2 readout achieved the highest discrimination among all sequence-model configurations evaluated in this study. DNABERT-2, NT-v2, Caduceus-Ph and modernGENA also improved across the three primary metrics at 501 bp relative to their respective 101-bp results. HyenaDNA followed the opposite pattern, with MCC decreasing from 0.209 to 0.122, AUROC from 0.655 to 0.631, and AUPRC from 0.392 to 0.358.

The intermediate 201-bp window provided additional insight into these differences. Caduceus-Ph achieved its highest MCC at 201 bp (0.284), followed by 0.273 at 501 bp. A similar pattern was observed for modernGENA, whose MCC reached 0.313 at 201 bp before declining to 0.263 at 501 bp. Thus, the relative benefit of additional sequence varied with both architecture and context length. All three context conditions used frozen model parameters, with a separate logistic-regression classifier and regularization selection for each model-context combination. The reported changes therefore reflect the performance of independently fitted frozen readouts at different sequence lengths. Results for all 18 model-context combinations are provided in Supplementary Table S3.

### 3.4 Native evolutionary-constraint and composite variant-impact scores show high discrimination

GPN-Star V100 and AlphaGenome Atlas AVI achieved higher discrimination than the frozen sequence-model readouts on the fixed test partition (Figure 2; Table 1). GPN-Star V100 uses multispecies sequence alignments to estimate evolutionary constraint, while AVI incorporates predicted molecular effects, conservation measures and variant-annotation information. Both approaches were evaluated using their existing continuous scores, with classification thresholds selected from the training partition.

AVI achieved an MCC of 0.938, AUROC of 0.991 and AUPRC of 0.987. The corresponding values for GPN-Star V100 were 0.929, 0.990 and 0.983. Among the sequence-model configurations, frozen Evo2 at 501 bp achieved the highest discrimination, with an MCC of 0.864, AUROC of 0.964 and AUPRC of 0.952. Both native scoring systems exceeded these values across all three primary metrics, although their biological inputs and scoring procedures differed from those used for the sequence-model readouts.

The exploratory paired comparison between AVI and GPN-Star V100 showed small numerical differences (Supplementary Table S4, Panel B). The AVI-minus-GPN-Star V100 difference was 0.009 for MCC (95% CI, −0.012 to 0.030), 0.0004 for AUROC (95% CI, −0.0103 to 0.0091) and 0.0044 for AUPRC (95% CI, −0.0060 to 0.0139). All three confidence intervals included zero, and the Holm-adjusted permutation p-values were 1.000 for each metric. The paired analysis therefore left the difference between the two scoring systems unresolved. Statistical equivalence was not evaluated.

We also examined the association between variant annotations and benchmark labels using six categories derived from ClinVar HGVS descriptions. A control based on category-specific positive-class frequencies estimated from the training partition achieved an MCC of 0.796, AUROC of 0.939 and AUPRC of 0.828. Repeating the analysis with training variants from each evaluation gene excluded gave an MCC of 0.796, AUROC of 0.927 and AUPRC of 0.842. These categories were assigned retrospectively using patterns in HGVS descriptions, without independent validation of molecular consequences. Their performance revealed substantial annotation-associated structure in the benchmark labels (Supplementary Table S14).

Within-category analysis provided further evidence of variation in classification performance (Supplementary Table S15). Among 267 amino-acid substitution variants, AUROC was 0.923 for AVI, 0.889 for GPN-Star V100 and 0.700 for frozen 101-bp Evo2. The corresponding values among 778 intronic variants were 0.9997, 0.986 and 0.856. Notably, 104 of the 116 positive-class intronic variants had HGVS offsets of ±1 or ±2, compared with none of the 662 negative-class variants. This pronounced difference in class composition around splice-adjacent positions may contribute to the high discrimination observed within the intronic category.

The evolutionary-timescale sensitivity analysis showed high discrimination across all three GPN-Star alignments. M447 achieved a slightly higher MCC than V100 (0.935 versus 0.929), accompanied by lower AUROC (0.989 versus 0.990) and AUPRC (0.979 versus 0.983). The comparisons were descriptive and did not establish a consistently superior evolutionary reference. Complete results are reported in Supplementary Tables S1 and S4.

### 3.5 Shared classification errors across expanded model configurations

Five of the 2,094 test variants were misclassified by all twelve original 101-bp sequence-model configurations, which included six frozen and six adapted readouts (Figure 4; Table 3). Adding the twelve frozen configurations evaluated at 201 and 501 bp reduced the shared-error intersection to two variants. When GPN-Star V100 and AlphaGenome AVI were included, a single variant remained misclassified across all 26 configurations.

These counts describe intersections of errors across progressively larger sets of configurations. By construction, adding configurations can only maintain or reduce the number of variants misclassified by every model. The observed progression from five to two to one reflects differences in classification outcomes across the evaluated approaches. Identifying which configurations produced correct predictions required knowledge of the benchmark labels, so the analysis provides no estimate of prospective ensemble performance. Section 3.7 examines the remaining universally misclassified variant. Further details on classification-error burden and native-score outcomes are reported in Supplementary Tables S5-S9.

**Figure 4.**
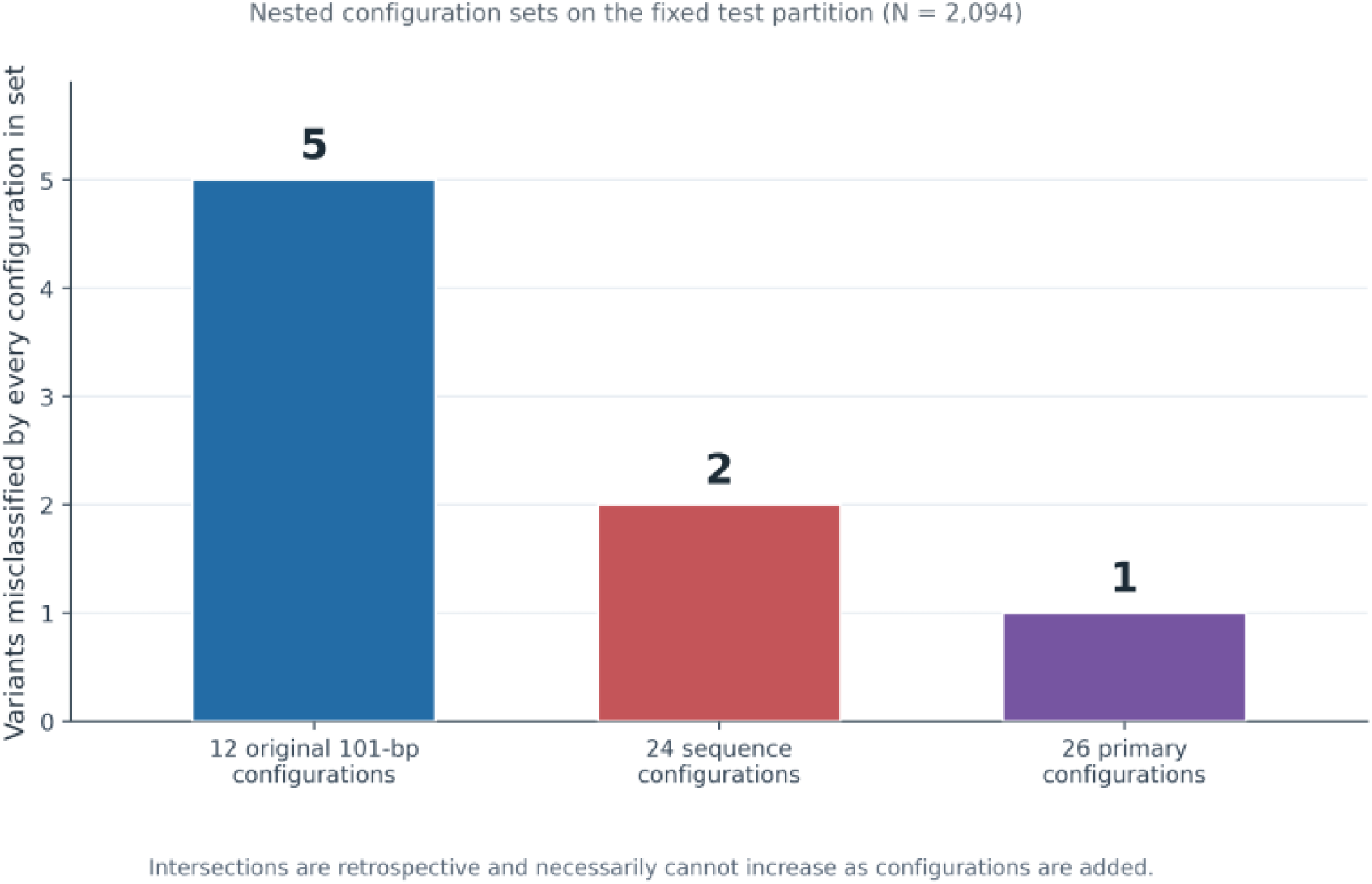
Shared classification errors across progressively expanded model configurations. Number of test variants (n = 2,094) misclassified by every configuration in three nested sets: the twelve original 101-bp sequence readouts (six frozen, six adapted); 24 sequence readouts after including frozen 201- and 501-bp configurations; and all 26 configurations after adding GPN-Star V100 and AlphaGenome Atlas AVI. The shared-error intersection decreased from five to two to one. Counts are retrospective and do not represent ensemble performance.

**Table 3.** Retrospective classification-error overlap and outcomes among difficult-variant subsets.

| Original subset | N | Any 201 bp | Any 501 bp | Any native | Any additional | All 24 wrong | All 26 wrong |
|---|---|---|---|---|---|---|---|
| All 12 wrong | 5 | 1 | 2 | 4 | 4 | 2 | 1 |
| ≥4 of 12 wrong | 787 | 709 | 731 | 772 | 782 | 2 | 1 |

Table 3 summarizes variants misclassified by all twelve original 101-bp sequence configurations (n = 5) and variants misclassified by at least four of those configurations (n = 787). The 201-bp, 501-bp and native-score columns indicate correct classification by at least one configuration within each additional strategy. These groups overlap, so their counts are not additive. Correct classification by any additional configuration represents a retrospective union determined using known benchmark labels, not the performance of a trained ensemble. Universal-error columns identify variants misclassified across all 24 sequence or all 26 primary configurations. Of the five recurrently difficult variants without a correct prediction from additional configurations, only one was incorrect across all 26.

### 3.6 Classification outcomes among recurrently difficult variants

Of the 2,094 test variants, 787 were misclassified by at least four of the twelve original 101-bp sequence-model configurations (Table 3). We examined how these variants were classified by the fourteen additional configurations. At least one frozen 201-bp model correctly classified 709 variants (90.1%), while the corresponding count for the 501-bp models was 731 (92.9%). GPN-Star V100 and AlphaGenome AVI together provided at least one correct classification for 772 variants (98.1%). Considering all fourteen additional configurations, 782 of the 787 variants (99.4%) were correctly classified by at least one approach. The groups of correctly classified variants overlapped across configurations.

Five variants remained incorrectly classified by every additional configuration. Four were located in SGCA and had received a correct classification from at least one of the original 101-bp readouts. The fifth, an ANO5 variant, was misclassified across all 26 configurations (Supplementary Tables S6 and S9).

We also examined native-score correctness across five strata defined by the number of errors among the 24 sequence-model configurations (Figure 5). At least one native score correctly classified 1,348 of 1,352 variants with 0–5 sequence-model errors, 351 of 355 with 6–11 errors, 252 of 254 with 12–17 errors, and 124 of 131 with 18–23 errors. Two variants were misclassified by all 24 sequence configurations, of which one received a correct classification from a native score. The small size of this final stratum limits any broader interpretation of its outcome.

**Figure 5.**
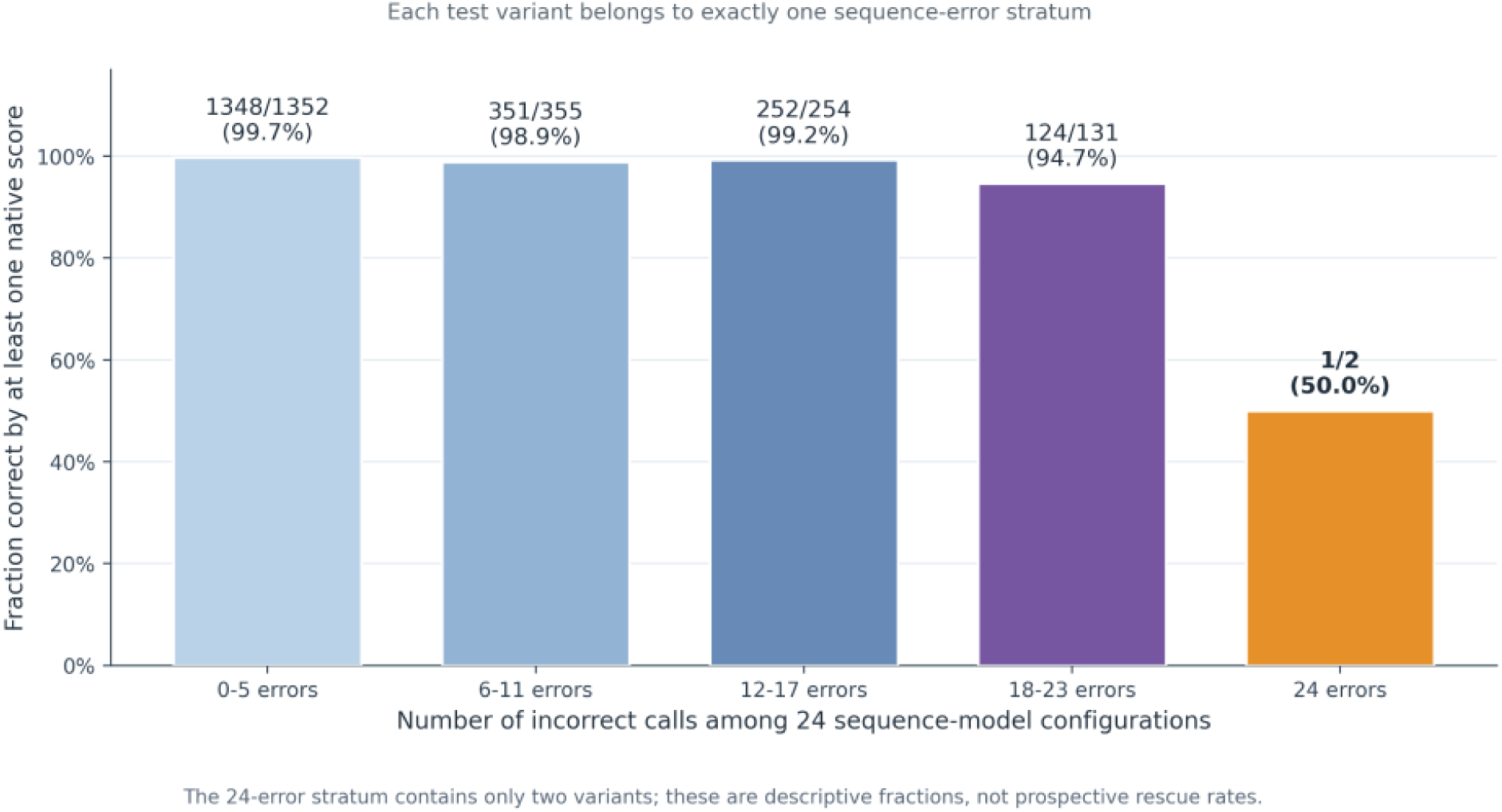
Native-score classification across strata of sequence-model error burden. The fixed test partition (n = 2,094) was divided into five mutually exclusive strata according to the number of incorrect predictions among 24 sequence-model configurations. Bars show the proportion of variants correctly classified by at least one of two native scoring approaches, GPN-Star V100 or AlphaGenome Atlas AVI. Numerators, denominators and percentages are displayed for each stratum. Classification thresholds were not optimized separately within strata. The stratum containing variants misclassified by all 24 sequence configurations comprised only two variants, one of which was correctly classified by a native score. These retrospective fractions should not be interpreted as prospective rescue rates.

These analyses describe the correct predictions available across the evaluated configurations for variants with different error burdens. Determining whether a prediction was correct required the known benchmark label, and the overlapping counts cannot be interpreted as independent contributions from each strategy. The observed coverage consequently provides no estimate of prospective variant rescue or the performance of a combined classifier.

### 3.7 A single ANO5 variant remains misclassified across all 26 configurations

ANO5 VariationID 4538462 (GRCh38 chr11:22239682 T>G) was the only variant misclassified by all 26 evaluated configurations (Table 3; Supplementary Table S9). The variant carried a positive clinical-significance label in the benchmark, yet every configuration assigned it to the negative class. These included the twelve original 101-bp sequence readouts, twelve additional frozen readouts at 201 and 501 bp, GPN-Star V100 and AlphaGenome Atlas AVI.

The disagreement with the benchmark label persisted following the evaluated supervised adaptation and across the longer frozen sequence contexts. Both native scoring systems also assigned the negative class despite drawing on different sources of biological information. The reason for this consistent misclassification remains unresolved. The available predictions provide insufficient evidence to question the clinical annotation or identify a molecular explanation for the shared error.

### 3.8 Gene-held-out performance of frozen sequence representations

We evaluated the frozen 101-bp representations under a gene-held-out downstream training procedure, where labelled variants from each evaluation gene were excluded from classifier development. Predictions from the ten evaluations were pooled across the fixed test partition of 2,094 variants, comprising 509 positive-class and 1,585 negative-class variants (Table 4).

**Table 4.**
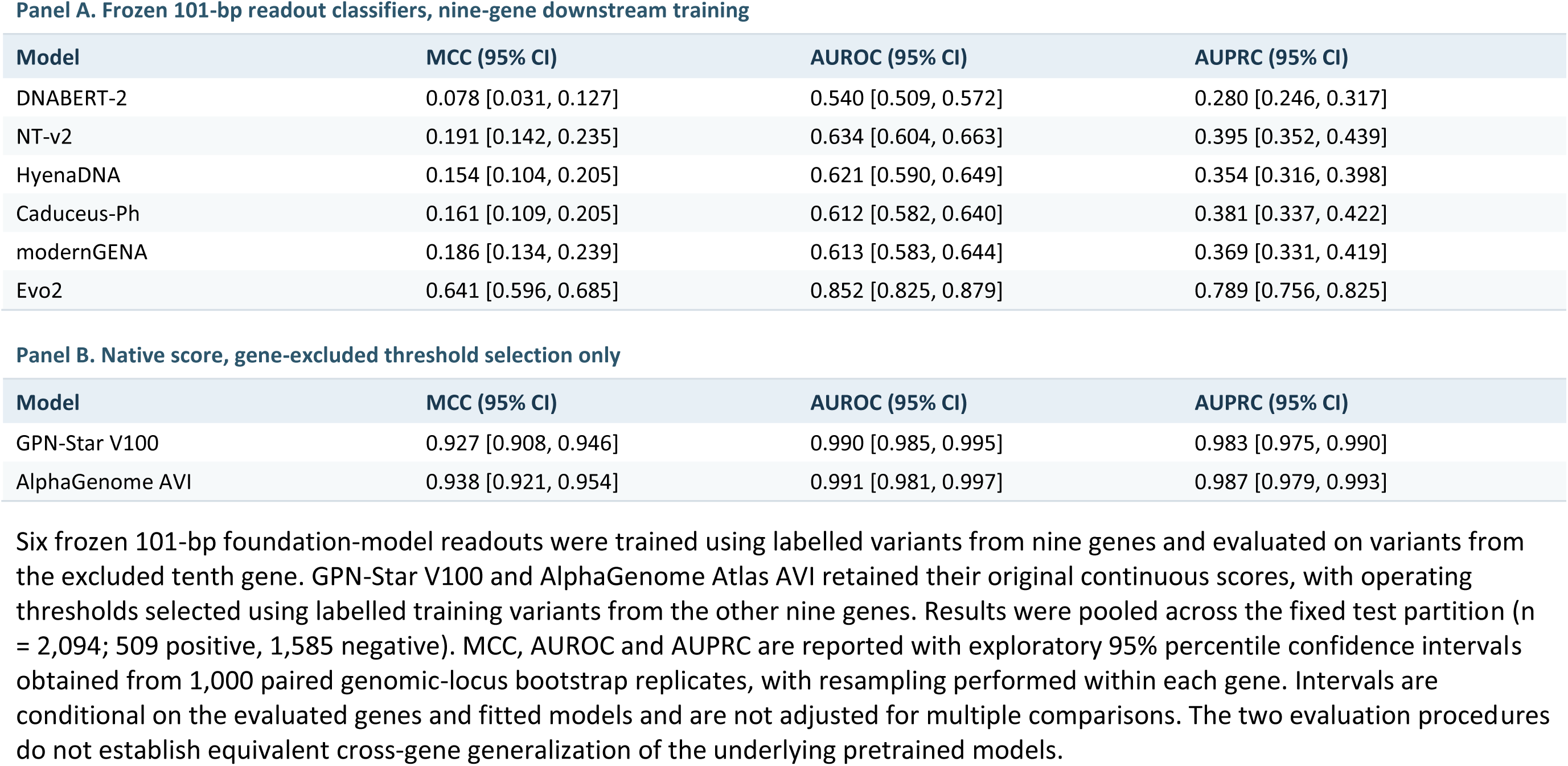
Pooled gene-held-out frozen-readout performance and gene-excluded native-score threshold evaluation.

| **Panel A. Frozen 101-bp readout classifiers, nine-gene downstream training** |  |  |  |
|---|---|---|---|
| **Model** | **MCC (95% CI)** | **AUROC (95% CI)** | **AUPRC (95% CI)** |
| DNABERT-2 | 0.078 [0.031, 0.127] | 0.540 [0.509, 0.572] | 0.280 [0.246, 0.317] |
| NT-v2 | 0.191 [0.142, 0.235] | 0.634 [0.604, 0.663] | 0.395 [0.352, 0.439] |
| HyenaDNA | 0.154 [0.104, 0.205] | 0.621 [0.590, 0.649] | 0.354 [0.316, 0.398] |
| Caduceus-Ph | 0.161 [0.109, 0.205] | 0.612 [0.582, 0.640] | 0.381 [0.337, 0.422] |
| modernGENA | 0.186 [0.134, 0.239] | 0.613 [0.583, 0.644] | 0.369 [0.331, 0.419] |
| Evo2 | 0.641 [0.596, 0.685] | 0.852 [0.825, 0.879] | 0.789 [0.756, 0.825] |
| **Panel B. Native score, gene-excluded threshold selection only** |  |  |  |
| **Model** | **MCC (95% CI)** | **AUROC (95% CI)** | **AUPRC (95% CI)** |
| GPN-Star V100 | 0.927 [0.908, 0.946] | 0.990 [0.985, 0.995] | 0.983 [0.975, 0.990] |
| AlphaGenome AVI | 0.938 [0.921, 0.954] | 0.991 [0.981, 0.997] | 0.987 [0.979, 0.993] |

Six frozen 101-bp foundation-model readouts were trained using labelled variants from nine genes and evaluated on variants from the excluded tenth gene. GPN-Star V100 and AlphaGenome Atlas AVI retained their original continuous scores, with operating thresholds selected using labelled training variants from the other nine genes. Results were pooled across the fixed test partition (n = 2,094; 509 positive, 1,585 negative). MCC, AUROC and AUPRC are reported with exploratory 95% percentile confidence intervals obtained from 1,000 paired genomic-locus bootstrap replicates, with resampling performed within each gene. Intervals are conditional on the evaluated genes and fitted models and are not adjusted for multiple comparisons. The two evaluation procedures do not establish equivalent cross-gene generalization of the underlying pretrained models.

Evo2 achieved the highest pooled discrimination among the six foundation models, with an MCC of 0.641 (95% bootstrap CI, 0.596–0.685), AUROC of 0.852 (0.825–0.879) and AUPRC of 0.789 (0.756–0.825). MCC values for the other five architectures ranged from 0.078 to 0.191. NT-v2 ranked second by MCC (0.191), with an AUROC of 0.634 and AUPRC of 0.395. Compared with NT-v2, Evo2 showed differences of 0.450 in MCC (95% CI, 0.388–0.516), 0.218 in AUROC (0.180–0.257) and 0.394 in AUPRC (0.344–0.443). All fifteen exploratory paired bootstrap intervals comparing Evo2 with the other five architectures excluded zero (Supplementary Table S11).

Performance varied across the ten evaluation genes. Evo2’s gene-level MCC ranged from 0.389 for SGCB to 0.702 for DNAJB6. Its AUROC reached 1.000 for DNAJB6, although this estimate involved only two positive-class test variants. We also recalculated pooled performance after excluding each gene in turn. Evo2’s MCC ranged from 0.613 to 0.653 across these analyses, indicating that the pooled result was relatively stable to the removal of individual evaluation genes. Supplementary Table S10 provides the complete gene-level results for all six frozen readouts.

Evo2 retained its performance advantage when downstream classifiers were trained without labelled examples from the evaluation gene. Possible exposure to these genes during foundation-model pretraining remains unresolved. The exploratory bootstrap intervals describe uncertainty conditional on the observed gene panel, genomic loci and fitted classifiers. They do not account for classifier refitting or variation associated with independently sampled genes.

### 3.9 Native variant scores under gene-excluded threshold selection

We evaluated GPN-Star V100 and AlphaGenome Atlas AVI using gene-excluded threshold selection (Table 4). Their continuous variant scores were retained, and a separate classification threshold was selected for each evaluation gene using labelled training variants from the remaining nine genes. The selected thresholds and corresponding gene-level results are provided in Supplementary Table S12.

GPN-Star V100 achieved a pooled MCC of 0.927 (95% bootstrap CI, 0.908–0.946), AUROC of 0.990 (0.985–0.995) and AUPRC of 0.983 (0.975–0.990). For AVI, the corresponding values were 0.938 (0.921–0.954), 0.991 (0.981–0.997) and 0.987 (0.979–0.993). AUROC and AUPRC were unchanged from the primary fixed-test evaluation because both the continuous scores and the evaluated variants remained identical. The gene-specific thresholds produced a small change in GPN-Star V100’s pooled MCC, from 0.929 in the primary evaluation to 0.927.

At the individual-gene level, GPN-Star V100 achieved MCC values ranging from 0.834 for SGCB to 1.000 for DNAJB6, FKRP and SGCG. AVI ranged from 0.827 for SGCA to 1.000 for DNAJB6, SGCB, SGCG and TCAP (Supplementary Table S10). The perfect MCC estimates require caution in the smaller test subsets. DNAJB6, SGCG and TCAP contained only 2, 11 and 6 positive-class variants, respectively.

Both native scoring approaches achieved higher pooled discrimination than Evo2, the strongest frozen gene-held-out readout. The paired MCC differences, calculated as Evo2 minus the native scorer, were −0.286 (95% CI, −0.332 to −0.244) for GPN-Star V100 and −0.297 (−0.342 to −0.253) for AVI (Supplementary Table S11).

The comparison involves two distinct evaluation procedures. For the frozen sequence representations, downstream classifiers were fitted using labelled variants from nine genes. GPN-Star V100 and AVI retained their previously developed scoring functions, with only the classification thresholds selected using gene-excluded training data. Their higher observed discrimination therefore has different implications for cross-gene performance. The experiment does not establish how either native predictor would perform on genes absent from its upstream development data.

## 4. Discussion

The comparison of six genomic foundation models revealed considerable variation in classification performance across architectures and evaluation procedures. Supervised adaptation improved discrimination for some models and reduced it for others, while increasing sequence context generally benefited the frozen readouts. The magnitude and direction of these changes differed across architectures. GPN-Star V100 and AlphaGenome Atlas AVI achieved higher discrimination using native scoring systems that incorporated additional sources of biological information.

The interpretation of these findings depends on the representation-extraction procedures, downstream training strategies and biological information available to each method. All evaluations used aggregate ClinVar clinical-significance labels for variants within ten LGMD-associated genes. The results describe classification performance under the protocols examined in this study. Their relevance to LGMD-specific pathogenicity and clinical diagnostic applications remains to be established.

### 4.1 Supervised adaptation does not consistently improve genomic representations

Supervised adaptation had markedly different effects across the six architectures (Figure 3; Table 2). Full fine-tuning increased MCC, AUROC and AUPRC for DNABERT-2, NT-v2 and HyenaDNA. ModernGENA showed declines in all three metrics, while Caduceus-Ph had a lower AUPRC after adaptation. For Caduceus-Ph, the changes in MCC and AUROC remained uncertain. These results suggest that the benefit of supervised training depends on the particular model and the conditions under which it is adapted.

Evo2 showed a substantial decline in MCC following LoRA adaptation, from 0.644 with frozen representations to 0.432 after adaptation. The experiment used one parameter-efficient adaptation design for Evo2, whereas the other five architectures underwent full backbone fine-tuning. The observed changes consequently reflect different training procedures across models. DNABERT-2 introduces a further consideration because frozen embeddings were extracted using MultiMolecule, while adaptation used the original implementation. Numerical equivalence between these implementations was not independently established under our extraction settings, leaving some uncertainty about the source of its performance improvement.

The changes in predicted probabilities were also notable. All six adapted configurations assigned the positive class to more test variants at the common decision threshold. Class-weighted optimization may have contributed to this shift, although its effect was not examined independently. Brier scores and expected calibration errors increased for every architecture after adaptation (Supplementary Table S2; Supplementary Figure S1). Even among the three models with improved discrimination, the adapted configurations produced less favorable raw-probability calibration.

Variation in adaptation outcomes has also been reported in broader genomic foundation-model benchmarking (Feng et al., 2025). In our evaluation, the changes in discrimination, positive-class predictions and calibration provide complementary evidence of how supervised adaptation affected each architecture. Examining these outcomes together is particularly important when comparing models trained under different optimization procedures.

### 4.2 Additional sequence context has different effects across architectures

Extending the input window from 101 to 501 bp improved MCC, AUROC and AUPRC for five of the six frozen architectures (Figure 3; Table 2). Evo2 showed the largest improvement in MCC, increasing from 0.644 at 101 bp to 0.864 at 501 bp, with progressively higher values across all three context lengths. HyenaDNA’s MCC declined as the window increased, whereas Caduceus-Ph and modernGENA achieved their highest MCC at 201 bp.

Additional flanking sequence may provide information about local nucleotide composition, splice-related regions or regulatory elements. We did not examine the contribution of individual positions or identify the sequence features responsible for the observed differences in discrimination. Although the pretrained backbones remained frozen, each model-context combination used a separately trained logistic-regression classifier, with regularization selected independently. The performance differences therefore also reflect the fitted downstream readouts.

Evo2’s discrimination increased with longer frozen input windows, while the evaluated 101-bp LoRA adaptation reduced performance. These experiments differed in their representation procedures and training interventions, limiting conclusions about the independent effects of either factor. Within the protocols examined here, extending sequence context was more beneficial for Evo2 than the selected supervised adaptation procedure.

### 4.3 Native variant-effect scores draw on distinct but overlapping biological information

GPN-Star V100 and AlphaGenome Atlas AVI achieved higher discrimination than the frozen sequence-model readouts, including Evo2 at 501 bp (Figure 2; Table 1). Their scoring procedures draw on biological information beyond the local DNA representations used by the downstream classifiers. GPN-Star estimates evolutionary constraint from multispecies sequence alignments and phylogenetic relationships, with mutation-rate calibration accounting for differences in background mutation processes (Ye et al., 2026). AVI combines AlphaGenome-derived molecular-effect predictions with AlphaMissense scores, evolutionary conservation and variant-consequence annotations (Avsec et al., 2026; Cheng et al., 2026). The AVI composite predictor was developed using allele-frequency-derived proxies for variant impact. Its training objective and input features therefore differ from those of the classifiers fitted to our ClinVar benchmark.

Some of the information used by these scoring systems may also contribute to clinical variant interpretation. Evolutionary conservation and predicted consequences are relevant to the evidence supporting clinical-significance assignments, creating a possible association between the native scores and the benchmark labels. The exploratory HGVS-category control identified pronounced differences in label composition across annotation categories (Supplementary Table S14). This provides further reason to consider annotation-associated information when interpreting the high discrimination of the native scores. The contribution of individual features remains unresolved.

AVI achieved slightly higher MCC, AUROC and AUPRC than GPN-Star V100, although the paired comparisons did not identify a statistically resolved difference in any of the three metrics (Supplementary Table S4, Panel B). Both scoring systems incorporate evolutionary conservation, and AVI combines it with several additional sources of evidence. We cannot determine from the present comparisons whether its molecular-effect predictions contributed independently to classification performance.

The feature attributions available for AVI could support a more detailed investigation of individual predictions. Their biological validity was outside the scope of this study, and the reported scores have not been calibrated as probabilities of disease causation. A controlled analysis of the separate conservation, consequence and molecular-effect components would be needed to assess their respective contributions to variant classification.

### 4.4 Classification-error patterns identify differences between prediction strategies

Five variants were misclassified by all twelve original 101-bp sequence configurations. When the twelve additional frozen 201- and 501-bp readouts were included, the shared-error intersection decreased to two variants. Adding GPN-Star V100 and AlphaGenome AVI reduced it to one (Figure 4; Table 3).

We examined these differences further among 787 variants misclassified by at least four of the original twelve configurations. Of these, 782 were correctly classified by at least one of the fourteen additional configurations (Figure 5; Table 3). Both longer sequence contexts and native scores provided correct predictions within this subset, although the strategy groups overlapped. Some variants were classified correctly by several configurations.

The shared-error intersection can only decrease or remain unchanged as additional configurations are included. Identifying which configuration correctly classified a variant also required access to its benchmark label. The resulting coverage represents an upper bound based on retrospective selection from the evaluated predictions. It does not estimate how a combined classifier would perform without knowledge of the correct label. Because the strategy-level counts overlap, they cannot be interpreted as independent contributions.

One ANO5 variant remained misclassified across all 26 configurations. The available predictions do not explain the underlying molecular cause, and differences in classification errors alone cannot establish that the models recognize distinct biological mechanisms. A subsequent study could examine whether sequence representations and native impact scores can be combined using a prespecified, label-independent procedure, with performance assessed through separate validation and independent testing.

### 4.5 Gene-held-out evaluation distinguishes downstream transfer from native-score threshold selection

We examined how frozen sequence representations performed when the downstream classifier was trained without labelled examples from the evaluation gene. For each of the ten genes, logistic regression was fitted using variants from the remaining nine genes and evaluated on the excluded gene’s variants in the fixed test partition. Evo2 achieved the highest pooled discrimination among the six foundation-model readouts (Table 4).

This design imposed a stronger restriction on labelled training data than the original locus-disjoint evaluation. Excluding a gene also changed the size and composition of the training subset, which complicates comparisons between the two experiments. Possible exposure to sequences from the evaluation genes during foundation-model pretraining remains unresolved.

For GPN-Star V100 and AlphaGenome AVI, the existing continuous variant scores were retained. A separate classification threshold was selected for each evaluation gene using labelled training variants from the other nine genes. Because the scores and test variants were unchanged, pooled AUROC and AUPRC matched the original fixed-test results. Threshold selection affected the classification metrics that depended on a decision cutoff. The sequence-model experiments involved fitting new downstream classifiers, while the native-score evaluations adjusted thresholds for previously developed predictors. Their results must be considered within these respective procedures.

The exploratory bootstrap analysis resampled genomic loci within each evaluation gene while retaining the fitted predictors. Its confidence intervals reflect variation among the observed loci in the ten-gene panel. They exclude uncertainty associated with classifier refitting, foundation-model pretraining, native-score development and the selection of additional genes. Independently assembled variant cohorts, together with better documentation of upstream training information, would provide a more stringent assessment of cross-gene performance.

### 4.6 Limitations and scope of interpretation

We selected aggregate ClinVar classifications for variants within ten LGMD-associated genes. Several of these genes are implicated in other clinical phenotypes, and the supporting evidence and review status of ClinVar records vary. Consequently, the reported performance reflects agreement with the selected ClinVar labels. The extent to which these results apply specifically to LGMD pathogenicity remains uncertain.

The exploratory HGVS-derived control showed a strong association between broad annotation categories and benchmark labels (Supplementary Table S14). Further analysis identified pronounced class differences among intronic variants with splice-adjacent positions (Supplementary Table S15). Such patterns may contribute to the discrimination achieved by the evaluated methods. However, the categories were inferred from HGVS naming conventions, and their molecular consequences were not independently verified. We therefore cannot determine how much of each model’s performance arises from particular biological features or from annotation-associated label structure.

The original partition excluded identical chromosome-position loci across the two sets, but imposed no minimum distance between distinct loci. In a retrospective proximity analysis, 1,820 of 2,094 test variants (86.9%) were located within 10 bp of at least one training locus. Nearby variants can share substantial portions of their reference-sequence windows, especially at longer context lengths. This proximity alone does not demonstrate label leakage. Nevertheless, the present evaluation offers limited evidence of performance on previously unseen genomic regions. Testing with region-disjoint partitions would help establish how well the observed discrimination extends beyond closely related sequence neighborhoods.

The six foundation models have distinct pretraining objectives, datasets, tokenization schemes and representation-extraction procedures. Each context-length condition used a separately fitted downstream classifier, while supervised adaptation involved full fine-tuning for five architectures and LoRA for Evo2. GPN-Star and AVI additionally relied on biological information supplied through their own scoring frameworks. These design differences complicate attempts to attribute performance changes to a particular architectural property, sequence length or adaptation procedure.

Our analysis was confined to single-nucleotide variants from the selected gene panel. Other variant classes, including insertions, deletions and structural variants, were not evaluated. Several gene-level subsets contained few positive-class variants, making their individual performance estimates less precise. We also lacked independently adjudicated clinical outcomes and functional measurements that could clarify the molecular basis of particular predictions. The study therefore leaves clinical diagnostic utility, calibrated disease-causation probabilities and performance in independent patient cohorts unresolved. Future evaluations should incorporate condition-specific clinical evidence, stronger genomic separation and clearer documentation of upstream model information.

## 5. Conclusion

We compared six genomic foundation models for ClinVar variant classification across ten LGMD-associated genes, examining frozen sequence representations, supervised adaptation and three input-context lengths. Adaptation improved discrimination for some architectures and reduced it for others. Longer sequence windows generally improved the frozen readouts, with Evo2 achieving the highest discrimination among the evaluated sequence-model configurations. Evo2 also performed best among the frozen representations in the gene-held-out downstream classifier evaluation.

GPN-Star V100 and AlphaGenome Atlas AVI achieved higher discrimination using native scores that incorporated additional biological information. The HGVS-derived control revealed strong associations between annotation categories and the benchmark labels. Variant-level analysis also identified differences in classification errors across configurations, although the retrospective coverage calculations cannot establish the performance of a prospective combined predictor. The contribution of individual molecular features to these predictions remains unresolved.

The results provide a reproducible account of how representation extraction, adaptation, sequence context and scoring procedures relate to classification performance in this ClinVar benchmark. Their interpretation remains constrained by the available clinical-significance labels and evaluation protocols. Assessment in region-disjoint datasets and independently assembled, condition-aware cohorts is needed to establish whether these patterns extend to less closely related variants and clinical settings.

## Data availability

The canonical ClinVar-derived variant benchmark, fixed training and test partitions, model predictions, variant-level scores and supporting numerical results are publicly available through the accompanying research repository and its archived release (https://doi.org/10.5281/zenodo.24028595).

Original ClinVar downloads, reference genome sequences and third-party model resources are not redistributed. Their use remains subject to the terms of the respective providers.

## Code availability

The analysis and validation scripts, together with the data required to reproduce the reported numerical summaries from archived predictions, are available at https://github.com/sarvezh1/lgmd-genomic-foundation-models.

The repository supports numerical reproduction and verification of the released results. It does not include all original training and inference workflows, pretrained model weights or adapted checkpoints, and therefore does not provide complete end-to-end reproduction of model training or inference.

## Author contributions

Sarvesh Galgale: Methodology, Software, Formal analysis, Validation, Investigation, Writing (original draft, review, and editing).

Isha Vats: Methodology, Software, Formal analysis, Validation, Investigation, Writing (original draft, review, and editing).

Josna Wilson: Methodology, Investigation, clinical interpretation, Writing (original draft, review, and editing).

Agnik Haldar: Conceptualization, Methodology, Supervision, Writing (review and editing).

