## Supplementary Information for "Genomic foundation models and complementary variant-scoring strategies in LGMD-associated genes: architecture, adaptation, and sequence context"

Supplementary Methods S1 | Supplementary Figures S1-S2 | Supplementary Tables S1-S15

#### Contents

| Item | Description | Page No. |
|---|---|---|
| **Supplementary Methods** | Extended computational methodology and provenance | **2** |
| **Supplementary Figures** | Supplementary Figures S1-S2 | **5** |
| **Supplementary Tables** | Supplementary Tables S1-S15 | **7** |

Terminology. All clinical-significance labels refer to the curated aggregate ClinVar variant classes within ten LGMD-associated genes, not independently established LGMD-specific pathogenicity. Unless otherwise stated, analyses use the fixed test partition of 2,094 SNVs.

### Supplementary Methods S1

### Extended computational methodology and provenance

#### S1.1 Benchmark and variant identity

The canonical benchmark comprised 6,960 ClinVar-derived single-nucleotide variants from ten LGMD-associated genes: ANO5, CAPN3, DNAJB6, DYSF, FKRP, LMNA, SGCA, SGCB, SGCG and TCAP. All variants were represented using GRCh38 genomic coordinates. The fixed partition contained 4,866 training variants and 2,094 test variants, with 1,585 negative-class and 509 positive-class variants in the test set. ClinVar VariationID was used to identify individual variants. Training and test records were separated at the chromosome-position locus level, ensuring that variants at identical genomic positions remained within the same partition.

The canonical variant-level dataset is available at data/canonical_variants.tsv.

#### S1.2 Sequence representations and supervised adaptation

Six genomic foundation models were evaluated: DNABERT-2, Nucleotide Transformer v2, HyenaDNA, Caduceus-Ph, modernGENA and Evo2. Frozen representations were assessed using 101-bp sequence windows in the primary benchmark, followed by separate context-length experiments at 201 and 501 bp. Supervised adaptation was restricted to 101-bp inputs and followed the architecture-specific procedures described in Section 2.4 of the main manuscript. Five architectures underwent full backbone fine-tuning, while Evo2 was adapted using LoRA.

The table below reports the selected learning rates and training epochs recorded in the archived adaptation-selection results.

| Model | Adaptation | Selected_LR | Best_Epoch |
|---|---|---|---|
| DNABERT-2 | Full FT | 3e-05 | 4 |
| NT-v2 | Full FT | 3e-05 | 3 |
| HyenaDNA | Full FT | 3e-05 | 3 |
| Caduceus-Ph | Full FT | 0.0001 | 5 |
| modernGENA | Full FT | 3e-05 | 3 |
| Evo2 | LoRA | 0.0003 | 5 |

##### S1.2.1 Pretrained checkpoint and execution provenance

The checkpoint identifiers and available revisions were reconciled with archived frozen-embedding extraction records and selected supervised-adaptation manifests. All six frozen architectures were evaluated at 101, 201 and 501 bp. The extraction manifests for the longer contexts identify the same checkpoint families recorded for the 101-bp evaluations. Supervised adaptation used 101-bp inputs exclusively. Checkpoint revisions are reported where they were preserved in the original execution records.

| Architecture | Frozen extraction checkpoint | Adapted-run checkpoint |
|---|---|---|
| DNABERT-2 | multimolecule/dnabert2<br>Revision not archived | zhihan1996/DNABERT-2-117M<br>rev 9c69a758f2f582f59a88a38948743ae5f733b23d |
| NT-v2 | InstaDeepAI/nucleotide-transformer-v2-500m-multi-species<br>rev 06615c1660c892fc199840c18123f8385b3542a8 | Same checkpoint and revision as frozen extraction |
| HyenaDNA | LongSafari/hyenadna-medium-160k-seqlen-hf<br>Frozen 101-bp revision not archived | LongSafari/hyenadna-medium-160k-seqlen-hf<br>rev 7ebf71773d22c0ede2cc55cb2be15ee8c289e1ce |
| Caduceus-Ph | kuleshov-group/caduceus-ph_seqlen-131k_d_model-256_n_layer-16<br>rev aa843f6d59724cddce28b6ebd075bfce2b6396ff | Same checkpoint and revision as frozen extraction |

| Architecture | Frozen extraction checkpoint | Adapted-run checkpoint |
|---|---|---|
| modernGENA | AIRI-Institute/moderngena-base<br>rev b26f1cb | Same checkpoint and revision as frozen extraction |
| Evo2 | evo2_7b_base<br>Layer blocks.28.mlp.l3 | evo2_7b_base<br>LoRA on block 28; layer blocks.28.mlp.l3 |

Frozen DNABERT-2 embeddings were extracted using the MultiMolecule implementation, while supervised adaptation used the original DNABERT-2 model repository. MultiMolecule describes its checkpoint as reproducing the intermediate representations of the original model. Run-specific numerical equivalence under the extraction and tokenization settings used in this study was not independently verified. Some frozen extraction records also lacked exact source revisions. The retained checkpoint identifiers, tokenizer implementations and execution environments therefore document the individual runs for which they were recorded.

Frozen representation extraction used architecture-specific tokenization and pooling procedures. DNABERT-2, NT-v2 and modernGENA used attention-mask mean pooling; HyenaDNA used nucleotide-position mean pooling; Caduceus-Ph used the mean of forward and reverse-complement representations; and Evo2 used mean-pooled activations from layer blocks.28.mlp.l3. These procedures are preserved in the archived context-length extraction scripts. Most recorded frozen extractions used Python 3.13 and PyTorch 2.11 on an NVIDIA L4 GPU. Evo2 extraction and adaptation used PyTorch 2.7.1 with CUDA 12.8, and its adaptation environment used Python 3.12.3. Some adapted runs also recorded CUDA 13.0. The software environments varied across the original experiments.

#### S1.3 Gene-held-out downstream evaluation

The cross-gene experiment used a leave-one-gene-out downstream classifier-training design. For each of the ten evaluation genes, supervised training was restricted to canonical training variants from the remaining nine genes. Predictions were then obtained for the excluded gene's variants within the original fixed-test partition. Across the ten evaluation folds, every one of the 2,094 fixed-test variants was evaluated exactly once.

For each of the six sequence models, frozen 101-bp alternate-minus-reference embeddings were used with a StandardScaler and L2-regularized logistic-regression classifier. Regularization was selected through five-fold StratifiedGroupKFold cross-validation, with chromosome-position loci defining the groups. Candidate values of C were 0.001, 0.01, 0.1, 1, 10 and 100. Selection maximized mean cross-validation MCC, with ties resolved in favor of the smallest C.

The selected classifier was fitted using the complete nine-gene training subset, and binary predictions were obtained at a probability threshold of 0.5. The fitting and evaluation methodology is also documented in the public repository (METHODS.md). Supplementary Table S13 provides the 60 selected gene/model regularization values.

#### S1.4 Native biological-impact scoring

GPN-Star V100 and AlphaGenome Atlas AVI were evaluated using their native variant-level scores. For GPN-Star V100, the continuous score was the sign-reversed mutation-rate-calibrated log-likelihood ratio (−LLR), with higher values indicating stronger predicted evolutionary constraint. AVI was evaluated using its PHRED-transformed score, where higher values indicate greater predicted variant impact. Scores from both systems were retained without further model training or recalibration.

For each gene-held-out evaluation, the operating threshold was selected by maximizing MCC on labelled training variants from the remaining nine genes. Ties were resolved by selecting the higher threshold. The excluded gene's test variants and labels were not involved in threshold selection. This procedure restricted supervised threshold selection to the other nine genes, while the upstream training data of the native scoring systems remained unchanged. Supplementary Table S12 reports the 20 selected gene-specific operating thresholds.

#### S1.5 Statistical analysis

MCC, AUROC and AUPRC were the primary discrimination metrics. Uncertainty in the exploratory cross-gene evaluation was estimated using 1,000 paired bootstrap replicates, with chromosome-position loci resampled with replacement within each of the ten evaluation genes. Paired model comparisons used identical resampled loci, and 95% confidence intervals were calculated using the percentile method. The intervals describe uncertainty conditional on the evaluated genes, their observed genomic loci and the fitted model readouts. They exclude variability associated with classifier refitting, sampling additional genes and assembling independent patient cohorts. Gene-level results and exploratory paired comparisons are reported in Supplementary Tables S10 and S11.

Three comparisons among the original 101-bp frozen and adapted configurations were selected after reviewing the primary results: frozen Evo2 versus adapted NT-v2, adapted Evo2 versus adapted NT-v2, and adapted NT-v2 versus adapted HyenaDNA. Differences in MCC, AUROC and AUPRC were evaluated using two-sided paired random-swap permutation tests with 10,000 permutations (seed 42). Predictions were swapped jointly for variants sharing a chromosome-position locus. Holm adjustment was applied across the nine resulting tests. Paired 95% percentile confidence intervals were calculated using the synchronized 10,000-replicate genomic-locus bootstrap. These post hoc comparisons were exploratory and were not specified before examining the primary results (Supplementary Table S2, Panel E).

A separate analysis compared AlphaGenome Atlas AVI with GPN-Star V100 using the same 2,094 fixed-test variants at 1,834 distinct genomic loci. Differences were calculated as AVI minus GPN-Star V100 for MCC, AUROC and AUPRC. Uncertainty was estimated from 10,000 paired genomic-locus bootstrap replicates (seed 42), with 95% confidence intervals obtained using the percentile method. Two-sided paired locus-level random-swap permutation tests were performed using 10,000 permutations (seed 42). Holm adjustment was restricted to the three native-score metrics, forming a separate correction family from the nine exploratory sequence-model comparisons. The resulting estimates characterize performance differences on the fixed test partition. The analyses were not designed to establish statistical equivalence or the independent contribution of individual biological information sources (Supplementary Table S4, Panel B).

Calibration was assessed using the original, unadjusted test probabilities from all twelve frozen and adapted 101-bp configurations. Brier score, logarithmic loss and expected calibration error (ECE) were calculated, with ECE estimated from ten equal-width probability bins. Calibration intercepts and slopes were obtained through diagnostic regressions against the test labels. These regressions were used exclusively to evaluate probability behavior and did not modify predictions or influence model selection. Intercepts and slopes are reported only for configurations with a successful diagnostic fit according to the archived quality-control flag. The remaining calibration metrics were calculated directly from predicted probabilities and remain reportable for configurations with unsuccessful regression fits (Supplementary Table S2, Panels B and F).

#### S1.6 Reproducibility and interpretation

The frozen-representation, supervised-adaptation, sequence-context, native-score and gene-held-out experiments followed different evaluation procedures. They varied in model training, representation extraction, downstream classifier fitting and available biological information. Performance differences across these experiments must therefore be interpreted within their respective protocols. The study used a curated ClinVar benchmark, where classification outcomes may be influenced by variant ascertainment, annotation-associated label structure and information encountered during upstream model development. The reported discrimination does not establish clinical diagnostic utility.

Supplementary analyses were assembled from archived scores, predictions and computational records. The gene-held-out results used previously saved model representations and evaluation outputs, without further updates to foundation-model parameters during supplementary assembly. The accompanying repository provides the retained numerical evidence and validation procedures supporting the reported analyses.

#### S1.7 Exploratory HGVS-category control

Six categories were assigned retrospectively using regular-expression patterns in the ClinVar HGVS-like Name field: stop-gained, synonymous, amino-acid substitution, untranslated region (UTR), intronic and unclassified, in that order of precedence. UTR-prefixed intronic descriptions were assigned to the UTR category. These categories were derived from naming patterns without independent molecular-consequence validation (Supplementary Table S14).

For the fixed-test control, category-specific pathogenic-class frequencies were calculated from the 4,866 training variants and assigned as continuous scores to the corresponding test variants. Binary predictions used a threshold of 0.5. The gene-excluded control repeated this procedure using training variants from the other nine genes for each evaluation gene. Predictions were pooled across the same 2,094 test variants, with AUROC and average precision calculated from continuous prevalence scores. Both analyses were exploratory and used annotation-derived information distinct from the inputs of the evaluated prediction models.

#### S1.8 Retrospective error-set coverage

The error-overlap analysis included 26 configurations: twelve frozen and adapted 101-bp sequence readouts, six frozen 201-bp readouts, six frozen 501-bp readouts, GPN-Star V100 and AlphaGenome Atlas AVI. GPN-Star M447 and P243 were excluded. Variants misclassified by at least four of the twelve original 101-bp configurations formed the recurrent-error subset (n = 787), including five variants misclassified by all twelve.

Correct classification by any of the fourteen additional configurations was assessed retrospectively using the known benchmark labels. The analysis describes the observed coverage across available predictions, without estimating the performance of a prospectively trained ensemble or label-independent selection procedure.

The universal-error intersection was evaluated across the nested sets of 12, 24 and 26 configurations. Its size can only remain unchanged or decrease as configurations are added. Variants without a correct prediction among the fourteen additional configurations may still have been correctly classified by an original 101-bp readout. Variant-level records and summary counts are provided in Supplementary Tables S5–S9.

#### Supplementary Figures

##### Supplementary Figure S1. Probability reliability of 101-bp sequence readouts

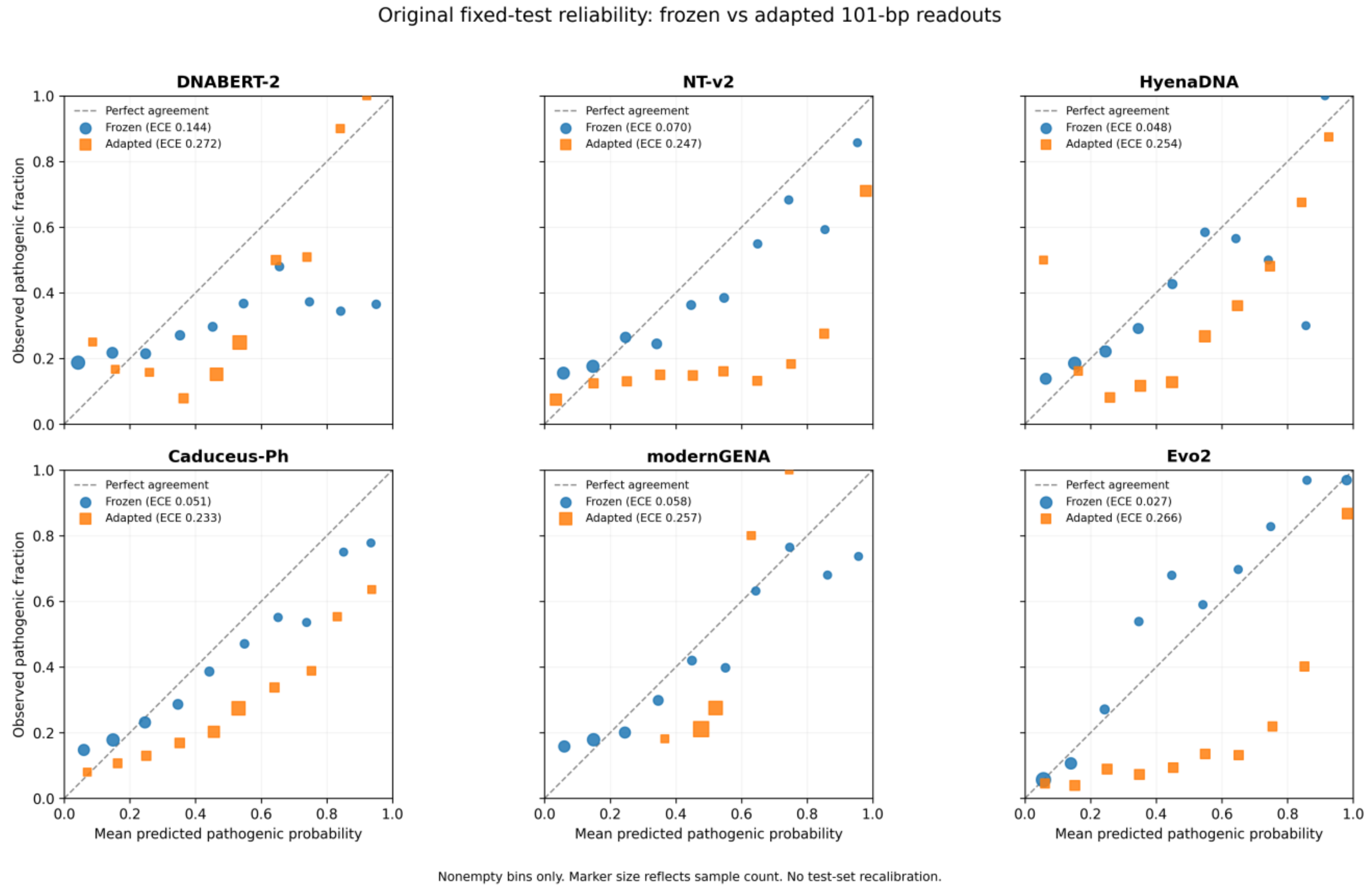


**Figure S1.** Raw-probability reliability of six frozen and six supervised-adapted 101-bp sequence readouts on the fixed test partition (n = 2,094; 509 positive-class variants). Each panel shows observed positive-class frequency against mean predicted probability in nonempty equal-width bins. The diagonal represents perfect agreement, and marker area indicates the number of variants per bin. ECE denotes ten-bin expected calibration error. No test-set recalibration was performed or confidence intervals calculated. The probabilities have not been validated as clinical pathogenicity estimates.

#### Supplementary Figure S2. Gene-level performance of frozen and adapted models

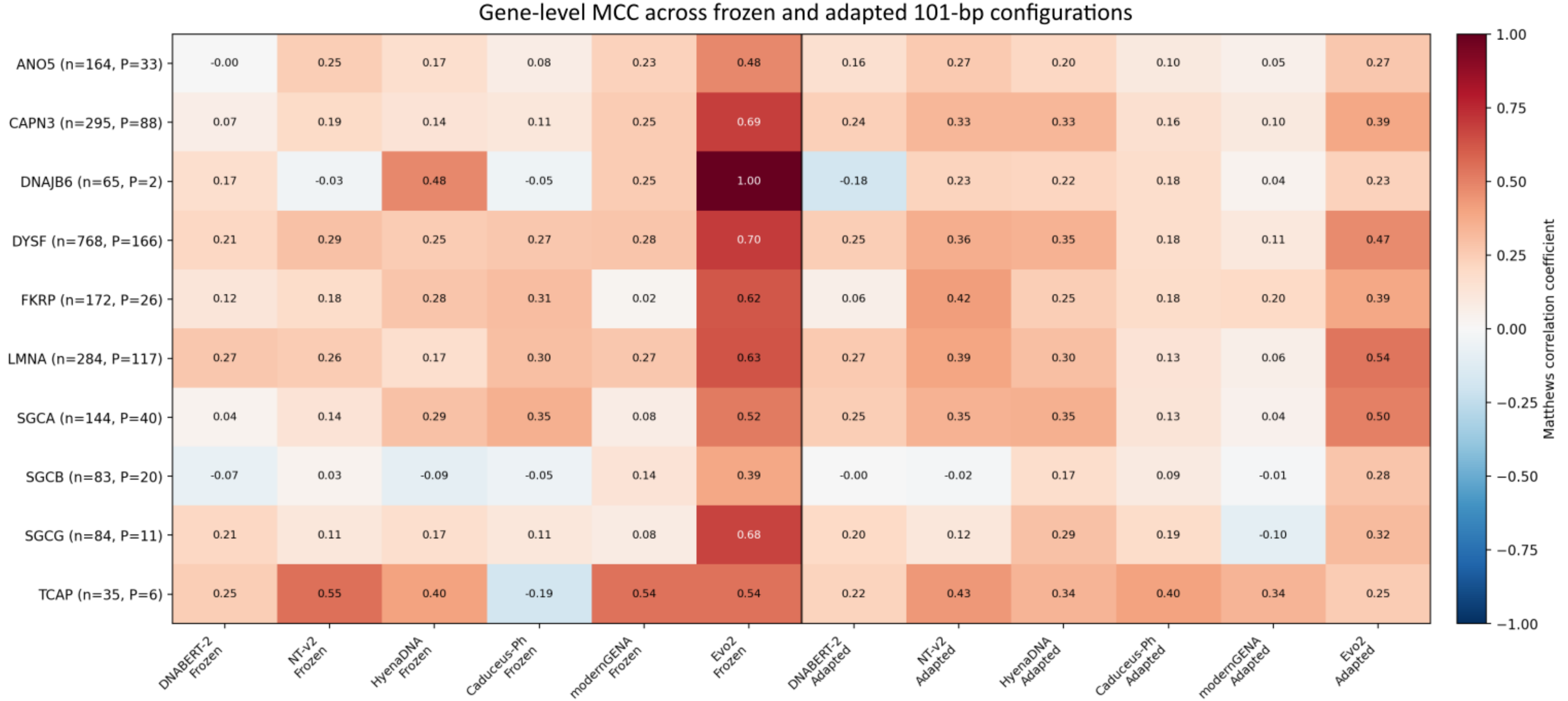


| | DNABERT-2 Frozen | NT-v2 Frozen | HyenaDNA Frozen | Caduceus-Ph Frozen | modernGENA Frozen | Evo2 Frozen | DNABERT-2 Adapted | NT-v2 Adapted | HyenaDNA Adapted | Caduceus-Ph Adapted | modernGENA Adapted | Evo2 Adapted |
|---|---|---|---|---|---|---|---|---|---|---|---|---|
| ANO5 (n=164, P=33) | -0.00 | 0.25 | 0.17 | 0.08 | 0.23 | 0.48 | 0.16 | 0.27 | 0.20 | 0.10 | 0.05 | 0.27 |
| CAPN3 (n=295, P=88) | 0.07 | 0.19 | 0.14 | 0.11 | 0.25 | 0.69 | 0.24 | 0.33 | 0.33 | 0.16 | 0.10 | 0.39 |
| DNAJB6 (n=65, P=2) | 0.17 | -0.03 | 0.48 | -0.05 | 0.25 | 1.00 | -0.18 | 0.23 | 0.22 | 0.18 | 0.04 | 0.23 |
| DYSF (n=768, P=166) | 0.21 | 0.29 | 0.25 | 0.27 | 0.28 | 0.70 | 0.25 | 0.36 | 0.35 | 0.18 | 0.11 | 0.47 |
| FKRP (n=172, P=26) | 0.12 | 0.18 | 0.28 | 0.31 | 0.02 | 0.62 | 0.06 | 0.42 | 0.25 | 0.18 | 0.20 | 0.39 |
| LMNA (n=284, P=117) | 0.27 | 0.26 | 0.17 | 0.30 | 0.27 | 0.63 | 0.27 | 0.39 | 0.30 | 0.13 | 0.06 | 0.54 |
| SGCA (n=144, P=40) | 0.04 | 0.14 | 0.29 | 0.35 | 0.08 | 0.52 | 0.25 | 0.35 | 0.35 | 0.13 | 0.04 | 0.50 |
| SGCB (n=83, P=20) | -0.07 | 0.03 | -0.09 | -0.05 | 0.14 | 0.39 | -0.00 | -0.02 | 0.17 | 0.09 | -0.01 | 0.28 |
| SGCG (n=84, P=11) | 0.21 | 0.11 | 0.17 | 0.11 | 0.08 | 0.68 | 0.20 | 0.12 | 0.29 | 0.19 | -0.10 | 0.32 |
| TCAP (n=35, P=6) | 0.25 | 0.55 | 0.40 | -0.19 | 0.54 | 0.54 | 0.22 | 0.43 | 0.34 | 0.40 | 0.34 | 0.25 |

**Figure S2.** Gene-level Matthews correlation coefficients (MCC) for six frozen and six supervised-adapted 101-bp sequence configurations on the original fixed test partition. Row labels indicate the total number of test variants (n) and positive-class variants (P) for each gene. DNAJB6 (P = 2), SGCG (P = 11) and TCAP (P = 6) have small positive-class subsets, limiting the interpretation of their individual MCC estimates. The separate gene-held-out downstream evaluations are reported in Supplementary Tables S10–S13.

### Supplementary Tables

Primary configurations, exploratory analyses and source-data qualifications are specified in individual table titles and notes. Statistics are rounded for presentation; archived machine-readable source tables preserve full reported precision.

#### Supplementary Table S1. Complete configuration inventory and primary fixed-test performance

Panel A summarizes the 24 sequence foundation-model configurations evaluated on the fixed test partition (n = 2,094; 509 positive, 1,585 negative). Frozen models used separately fitted logistic-regression readouts at 101, 201 and 501 bp. Supervised adaptation was evaluated at 101 bp only.

| Model | Regime | Context (bp) | MCC | AUROC | AUPRC | Precision | Recall | Specificity |
|---|---|---|---|---|---|---|---|---|
| DNABERT-2 | Frozen + LR | 101 | 0.151 | 0.598 | 0.320 | 0.388 | 0.265 | 0.866 |
| DNABERT-2 | Adapted | 101 | 0.215 | 0.696 | 0.479 | 0.326 | 0.745 | 0.505 |
| DNABERT-2 | Frozen + LR | 201 | 0.153 | 0.644 | 0.366 | 0.440 | 0.181 | 0.926 |
| DNABERT-2 | Frozen + LR | 501 | 0.313 | 0.698 | 0.507 | 0.780 | 0.189 | 0.983 |
| NT-v2 | Frozen + LR | 101 | 0.234 | 0.656 | 0.430 | 0.516 | 0.253 | 0.924 |
| NT-v2 | Adapted | 101 | 0.336 | 0.794 | 0.686 | 0.403 | 0.743 | 0.647 |
| NT-v2 | Frozen + LR | 201 | 0.285 | 0.670 | 0.466 | 0.577 | 0.281 | 0.934 |
| NT-v2 | Frozen + LR | 501 | 0.386 | 0.700 | 0.562 | 0.718 | 0.320 | 0.960 |
| HyenaDNA | Frozen + LR | 101 | 0.209 | 0.655 | 0.392 | 0.548 | 0.179 | 0.953 |
| HyenaDNA | Adapted | 101 | 0.306 | 0.724 | 0.503 | 0.382 | 0.741 | 0.616 |
| HyenaDNA | Frozen + LR | 201 | 0.180 | 0.636 | 0.377 | 0.503 | 0.167 | 0.947 |
| HyenaDNA | Frozen + LR | 501 | 0.122 | 0.631 | 0.358 | 0.439 | 0.120 | 0.951 |
| Caduceus-Ph | Frozen + LR | 101 | 0.223 | 0.656 | 0.423 | 0.545 | 0.204 | 0.945 |
| Caduceus-Ph | Adapted | 101 | 0.160 | 0.630 | 0.360 | 0.306 | 0.682 | 0.504 |
| Caduceus-Ph | Frozen + LR | 201 | 0.284 | 0.675 | 0.476 | 0.709 | 0.187 | 0.975 |
| Caduceus-Ph | Frozen + LR | 501 | 0.273 | 0.687 | 0.483 | 0.689 | 0.183 | 0.974 |
| modernGENA | Frozen + LR | 101 | 0.249 | 0.663 | 0.446 | 0.573 | 0.224 | 0.946 |
| modernGENA | Adapted | 101 | 0.089 | 0.593 | 0.374 | 0.288 | 0.499 | 0.603 |
| modernGENA | Frozen + LR | 201 | 0.313 | 0.684 | 0.475 | 0.622 | 0.285 | 0.944 |
| modernGENA | Frozen + LR | 501 | 0.263 | 0.685 | 0.488 | 0.725 | 0.155 | 0.981 |
| Evo2 | Frozen + LR | 101 | 0.644 | 0.879 | 0.818 | 0.884 | 0.570 | 0.976 |
| Evo2 | Adapted | 101 | 0.432 | 0.866 | 0.784 | 0.441 | 0.849 | 0.654 |
| Evo2 | Frozen + LR | 201 | 0.757 | 0.924 | 0.886 | 0.927 | 0.701 | 0.982 |
| Evo2 | Frozen + LR | 501 | 0.864 | 0.964 | 0.952 | 0.974 | 0.819 | 0.993 |

Abbreviations: LR, L2-regularized logistic regression; MCC, Matthews correlation coefficient; AUROC, area under the receiver operating characteristic curve; AUPRC, area under the precision–recall curve. Frozen sequence readouts and adapted models used a 0.5 probability cutoff. Specificity for 201/501-bp rows was calculated from the source confusion matrices (TN/(TN + FP)).

#### Supplementary Table S1 (continued)

Panel B. The four native variant-impact scoring configurations. V100 and AVI are primary comparators; M447 and P243 are GPN-Star sensitivity analyses, not additional primary strategies.

| Native configuration | Role | Basis | MCC | AUROC | AUPRC | Specificity | Threshold |
|---|---|---|---|---|---|---|---|
| GPN-Star V100 | Primary | 100-vertebrate | 0.929 | 0.990 | 0.983 | 0.987 | 5.767 |
| GPN-Star M447 | Sensitivity | 447-mammal | 0.935 | 0.989 | 0.979 | 0.978 | 4.889 |
| GPN-Star P243 | Sensitivity | 243-primate | 0.900 | 0.973 | 0.956 | 0.983 | 4.897 |
| AlphaGenome Atlas AVI | Primary | Composite AVI impact score | 0.938 | 0.991 | 0.987 | 0.994 | 23.8766 |

GPN-Star scores were calculated using the sign-reversed mutation-rate-calibrated log-likelihood ratio (−llr_calibrated). AlphaGenome Atlas AVI used a PHRED-transformed composite impact score incorporating AlphaGenome-derived molecular-effect predictions, AlphaMissense, conservation and variant-consequence features. Decision thresholds were selected on the canonical training partition (n = 4,866) and applied unchanged to the fixed test partition (n = 2,094). The two scoring systems use different score scales and biological inputs.

The primary error-overlap analysis included the 24 sequence configurations in Panel A, GPN-Star V100 and AlphaGenome Atlas AVI. GPN-Star M447 and P243 were retained for evolutionary-timescale sensitivity analysis and excluded from the 26-configuration error comparison.

#### Supplementary Table S2. Effects of supervised adaptation at 101 bp

#### Panel A. Frozen-versus-adapted changes in discrimination.

Each entry reports the adapted-minus-frozen difference in MCC, AUROC or AUPRC, followed by its 95% percentile confidence interval from 10,000 paired genomic-locus bootstrap replicates.

| Model | Adaptation | ΔMCC [95% CI] | ΔAUROC [95% CI] | ΔAUPRC [95% CI] |
|---|---|---|---|---|
| DNABERT-2 | Full FT | +0.064 [+0.005, +0.125] | +0.098 [+0.059, +0.137] | +0.159 [+0.108, +0.203] |
| NT-v2 | Full FT | +0.102 [+0.041, +0.163] | +0.138 [+0.105, +0.172] | +0.256 [+0.208, +0.303] |
| HyenaDNA | Full FT | +0.098 [+0.041, +0.155] | +0.068 [+0.041, +0.095] | +0.111 [+0.069, +0.150] |
| Caduceus-Ph | Full FT | −0.063 [−0.127, +0.001] | −0.026 [−0.064, +0.012] | −0.062 [−0.111, −0.012] |
| modernGENA | Full FT | −0.160 [−0.222, −0.097] | −0.070 [−0.105, −0.033] | −0.072 [−0.116, −0.027] |
| Evo2 | LoRA | −0.212 [−0.255, −0.168] | −0.013 [−0.029, +0.003] | −0.034 [−0.052, −0.016] |

Confidence intervals excluding zero support a directional change within the fixed benchmark, while intervals spanning zero leave the direction uncertain. These exploratory estimates are conditional on the evaluated test set. Five architectures underwent full backbone fine-tuning, and Evo2 was adapted using LoRA.

#### Panel B. Raw probability behavior and calibration diagnostics

| Model | Brier, frozen | Brier, adapted | ECE, frozen | ECE, adapted | Mean P(+), frozen | Mean P(+), adapted |
|---|---|---|---|---|---|---|
| DNABERT-2 | 0.214 | 0.239 | 0.144 | 0.272 | 0.251 | 0.512 |
| NT-v2 | 0.175 | 0.227 | 0.070 | 0.247 | 0.238 | 0.472 |
| HyenaDNA | 0.175 | 0.226 | 0.048 | 0.254 | 0.239 | 0.495 |
| Caduceus-Ph | 0.174 | 0.235 | 0.051 | 0.233 | 0.240 | 0.476 |
| modernGENA | 0.170 | 0.244 | 0.058 | 0.257 | 0.239 | 0.496 |
| Evo2 | 0.091 | 0.199 | 0.027 | 0.266 | 0.234 | 0.509 |

Brier scores and expected calibration errors (ECE) were calculated from the original test probabilities without post hoc recalibration. Lower values indicate better performance under the respective calibration metrics. The predicted probabilities have not been validated for clinical pathogenicity assessment.

#### Supplementary Table S2 (continued)

#### Panel C. Counts of classification transitions from frozen to adapted at the common 0.5 threshold.

| Model | All rescued | All harmed | Positive rescued | Positive harmed | Negative rescued | Negative harmed |
|---|---|---|---|---|---|---|
| DNABERT-2 | 354 | 682 | 266 | 22 | 88 | 660 |
| NT-v2 | 324 | 513 | 263 | 14 | 61 | 499 |
| HyenaDNA | 304 | 552 | 290 | 4 | 14 | 548 |
| Caduceus-Ph | 295 | 751 | 263 | 20 | 32 | 731 |
| modernGENA | 216 | 620 | 177 | 37 | 39 | 583 |
| Evo2 | 145 | 513 | 144 | 2 | 1 | 511 |

Rescued = incorrect under frozen evaluation and correct after adaptation; harmed = correct under frozen evaluation and incorrect after adaptation. “Positive” and “negative” refer to benchmark ClinVar clinical-significance classes.

#### Panel D. Direction supported across seven prespecified stable genes

| Model | MCC (+/−/?) | AUROC (+/−/?) | AUPRC (+/−/?) |
|---|---|---|---|
| DNABERT-2 | +1 / −0 / ?6 | +3 / −0 / ?4 | +4 / −0 / ?3 |
| NT-v2 | +1 / −0 / ?6 | +5 / −0 / ?2 | +5 / −0 / ?2 |
| HyenaDNA | +2 / −0 / ?5 | +3 / −0 / ?4 | +3 / −0 / ?4 |
| Caduceus-Ph | +0 / −1 / ?6 | +0 / −0 / ?7 | +0 / −1 / ?6 |
| modernGENA | +0 / −2 / ?5 | +0 / −2 / ?5 | +0 / −0 / ?7 |
| Evo2 | +0 / −4 / ?3 | +1 / −1 / ?5 | +0 / −3 / ?4 |

Eligible genes: ANO5, CAPN3, DYSF, FKRP, LMNA, SGCA and SGCB. DNAJB6 (2 positive test variants), SGCG (11) and TCAP (6) were excluded from this subgroup inferential summary using the predefined rule of at least 20 test variants in each class. Counts denote bootstrap intervals supporting increase (+), decrease (−), or uncertain direction (?), with 5,000 within-gene locus-cluster resamples. All ten genes remain represented in the overall fixed-test evaluation.

#### Panel E. Targeted post hoc model comparisons

Paired differences are expressed as the first configuration minus the second. Confidence intervals are from 10,000 paired locus-bootstrap replicates; the two-sided random-swap permutation tests used 10,000 permutations at the genomic-locus level. Holm correction covered all nine metric tests. Comparisons were chosen after examination of primary results and are exploratory.

| Comparison (first – second) | Metric | Difference | 95% bootstrap CI | Raw p | Holm p |
|---|---|---|---|---|---|
| Frozen Evo2 – adapted NT-v2 | MCC | +0.3077 | [+0.2515, +0.3620] | 0.00010 | 0.00090 |
| Frozen Evo2 – adapted NT-v2 | AUROC | +0.0848 | [+0.0518, +0.1165] | 0.00010 | 0.00090 |
| Frozen Evo2 – adapted NT-v2 | AUPRC | +0.1328 | [+0.0904, +0.1751] | 0.00010 | 0.00090 |
| Adapted Evo2 – adapted NT-v2 | MCC | +0.0960 | [+0.0462, +0.1436] | 0.00030 | 0.00090 |
| Adapted Evo2 – adapted NT-v2 | AUROC | +0.0720 | [+0.0392, +0.1032] | 0.00010 | 0.00090 |
| Adapted Evo2 – adapted NT-v2 | AUPRC | +0.0988 | [+0.0535, +0.1423] | 0.00010 | 0.00090 |
| Adapted NT-v2 – adapted HyenaDNA | MCC | +0.0301 | [-0.0164, +0.0771] | 0.20018 | 0.20018 |
| Adapted NT-v2 – adapted HyenaDNA | AUROC | +0.0702 | [+0.0423, +0.0982] | 0.00010 | 0.00090 |
| Adapted NT-v2 – adapted HyenaDNA | AUPRC | +0.1822 | [+0.1382, +0.2256] | 0.00010 | 0.00090 |

Adjusted p-values describe the selected exploratory family only. Eight tests had Holm-adjusted p = 0.00090. Adapted NT-v2 and HyenaDNA did not have a resolved MCC difference (Holm-adjusted p = 0.20018), despite differences in AUROC and AUPRC. None of these comparisons establishes clinical utility or equivalent adaptation regimes.

#### Panel F. Additional raw-probability calibration diagnostics

Panel F extends the raw-probability results in Panel B with logarithmic loss, calibration intercepts and calibration slopes. The regression estimates were used for diagnostic evaluation and did not modify test predictions. Intercepts and slopes are reported only when the archived calibration fit-success flag indicated a successful fit.

| Model | Regime | Log loss | Cal. intercept | Cal. slope | Fit QC |
|---|---|---|---|---|---|
| DNABERT-2 | frozen | 0.7166 | -0.9018 | 0.1556 | Passed |
| DNABERT-2 | adapted | 0.6698 | -1.3449 | 1.7630 | Passed |
| NT-v2 | frozen | 0.5496 | -0.5045 | 0.4888 | Passed |
| NT-v2 | adapted | 0.7457 | -1.5094 | 0.3849 | Passed |
| HyenaDNA | frozen | 0.5419 | -0.4406 | 0.5587 | Passed |
| HyenaDNA | adapted | 0.6461 | -1.2868 | 1.0682 | Passed |
| Caduceus-Ph | frozen | 0.5432 | -0.4711 | 0.5312 | Passed |
| Caduceus-Ph | adapted | 0.6626 | Not reported† | Not reported† | Flagged unsuccessful |
| modernGENA | frozen | 0.5353 | Not reported† | Not reported† | Flagged unsuccessful |
| modernGENA | adapted | 0.6814 | Not reported† | Not reported† | Flagged unsuccessful |
| Evo2 | frozen | 0.3225 | +0.1465 | 1.0351 | Passed |
| Evo2 | adapted | 0.6071 | -1.8706 | 0.7233 | Passed |

† The original source table contains numeric diagnostic intercept/slope outputs for the three rows flagged unsuccessful, but these estimates are withheld from presentation because their fit success was not established. Flagged rows: adapted Caduceus-Ph and frozen/adapted modernGENA. The archived files do not identify the cause of unsuccessful fitting. All log-loss values, Brier scores and ECE values remain reportable. No test-label-derived calibration regression was applied to transform predictions.

#### Supplementary Table S3. Frozen sequence-context evaluation at 101, 201 and 501 bp

All six architectures were evaluated at three centered reference/alternate sequence lengths using the same canonical training (N = 4,866) and test partitions (N = 2,094). A separate logistic-regression readout and regularization selection was used for each model/context condition.

| Model | Context (bp) | Chosen C | MCC | AUROC | AUPRC | Recall | Specificity | CV mean MCC |
|---|---|---|---|---|---|---|---|---|
| DNABERT-2 | 101 | 1 | 0.151 | 0.598 | 0.320 | 0.265 | 0.866 | Not archived |
| DNABERT-2 | 201 | 0.01 | 0.153 | 0.644 | 0.366 | 0.181 | 0.926 | 0.159 |
| DNABERT-2 | 501 | 0.001 | 0.313 | 0.698 | 0.507 | 0.189 | 0.983 | 0.306 |
| NT-v2 | 101 | 0.01 | 0.234 | 0.656 | 0.430 | 0.253 | 0.924 | Not archived |
| NT-v2 | 201 | 0.01 | 0.285 | 0.670 | 0.466 | 0.281 | 0.934 | 0.321 |
| NT-v2 | 501 | 0.01 | 0.386 | 0.700 | 0.562 | 0.320 | 0.960 | 0.374 |
| HyenaDNA | 101 | 100 | 0.209 | 0.655 | 0.392 | 0.179 | 0.953 | Not archived |
| HyenaDNA | 201 | 10 | 0.180 | 0.636 | 0.377 | 0.167 | 0.947 | 0.169 |
| HyenaDNA | 501 | 100 | 0.122 | 0.631 | 0.358 | 0.120 | 0.951 | 0.123 |
| Caduceus-Ph | 101 | 100 | 0.223 | 0.656 | 0.423 | 0.204 | 0.945 | Not archived |
| Caduceus-Ph | 201 | 0.1 | 0.284 | 0.675 | 0.476 | 0.187 | 0.975 | 0.226 |
| Caduceus-Ph | 501 | 0.1 | 0.273 | 0.687 | 0.483 | 0.183 | 0.974 | 0.234 |
| modernGENA | 101 | 0.01 | 0.249 | 0.663 | 0.446 | 0.224 | 0.946 | Not archived |
| modernGENA | 201 | 0.01 | 0.313 | 0.684 | 0.475 | 0.285 | 0.944 | 0.319 |
| modernGENA | 501 | 0.001 | 0.263 | 0.685 | 0.488 | 0.155 | 0.981 | 0.255 |
| Evo2 | 101 | 0.1 | 0.644 | 0.879 | 0.818 | 0.570 | 0.976 | Not archived |
| Evo2 | 201 | 0.1 | 0.757 | 0.924 | 0.886 | 0.701 | 0.982 | 0.780 |
| Evo2 | 501 | 0.1 | 0.864 | 0.964 | 0.952 | 0.819 | 0.993 | 0.816 |

The 101-bp entries were taken from the primary frozen benchmark and retained without additional model fitting. Cross-validation means and standard deviations for these six configurations were unavailable in the archived context-analysis summary and were left unreported. Their confusion-matrix counts were recovered from Table S1 and preserved in the accompanying machine-readable TSV.

#### Supplementary Table S4

#### Panel A. Native-score configurations and evolutionary-timescale sensitivity

Panel A reports performance for four native-score configurations on the fixed test partition of 2,094 variants. GPN-Star V100 served as the primary evolutionary-constraint comparator, with M447 and P243 used to examine alternative evolutionary alignments. AlphaGenome Atlas AVI was evaluated as a separate composite variant-impact predictor.

| Score | Role | Reference basis | Threshold | MCC | AUROC | AUPRC | TN | FP | FN | TP |
|---|---|---|---|---|---|---|---|---|---|---|
| GPN-Star V100 | Primary | 100-vertebrate | 5.767 | 0.929 | 0.990 | 0.983 | 1564 | 21 | 33 | 476 |
| GPN-Star M447 | Sensitivity | 447-mammal | 4.889 | 0.935 | 0.989 | 0.979 | 1550 | 35 | 16 | 493 |
| GPN-Star P243 | Sensitivity | 243-primate | 4.897 | 0.900 | 0.973 | 0.956 | 1558 | 27 | 49 | 460 |
| AlphaGenome Atlas AVI | Primary | Composite predictor | 23.8766 | 0.938 | 0.991 | 0.987 | 1575 | 10 | 37 | 472 |

GPN-Star-V100: 100 vertebrate genomes; GPN-Star-M447: 447 mammalian genomes; GPN-Star-P243: 243 primate genomes. GPN-Star thresholds operate on −llr_calibrated scores; AVI uses the PHRED-transformed Atlas quantile. Higher scores correspond to greater predicted variant impact/constraint, but are not calibrated probabilities of disease causation.

Thresholds were selected on the original 4,866-variant training partition to maximize MCC and applied unchanged to the 2,094-variant fixed test set (509 positive; 1,585 negative). Confusion matrices and other secondary metrics in the accompanying curated TSV were obtained directly from the archived native-score test-metric exports rather than estimated from rounded supplementary statistics.

#### Panel B. Direct paired comparison of AlphaGenome AVI and GPN-Star V100

Metric differences are AlphaGenome AVI minus GPN-Star V100 on the fixed 2,094-variant test partition. Positive values favor AVI. Intervals and tests use paired resampling at the genomic-locus level.

| Metric | Difference (AVI – GPN) | 95% paired bootstrap CI | Raw permutation p | Holm-adjusted p |
|---|---|---|---|---|
| MCC | +0.009014 | [−0.012381, +0.030361] | 0.462354 | 1.000000 |
| AUROC | +0.000381 | [−0.010257, +0.009068] | 0.978002 | 1.000000 |
| AUPRC | +0.004355 | [−0.005964, +0.013889] | 0.885111 | 1.000000 |

Paired analyses used chromosome-position loci as the resampling unit, with 10,000 bootstrap replicates and 10,000 random-swap permutations (seed 42). Confidence intervals were calculated using the 95% percentile method. Holm correction was applied across MCC, AUROC and AUPRC within this native-score comparison. All three confidence intervals included zero, and none of the Holm-adjusted tests identified a statistically resolved difference. Statistical equivalence was not evaluated. Numerical results were obtained from the archived paired native-score comparison records and associated analysis manifest.

#### Supplementary Table S5. Five variants misclassified by all 12 original 101-bp configurations

| ClinVar ID | Gene | GRCh38 position | REF>ALT | 201 bp correct /6 | 501 bp correct /6 | AVI correct | GPN-V100 correct | All 24 wrong | All 26 wrong |
|---|---|---|---|---|---|---|---|---|---|
| 2163 | ANO5 | chr11:22255485 | C>G | 0 | 0 | Yes | Yes | Yes | No |
| 1334893 | FKRP | chr19:46755953 | G>A | 0 | 1 | Yes | Yes | No | No |
| 3677001 | LMNA | chr1:156136074 | C>A | 1 | 0 | Yes | Yes | No | No |
| 4538462 | ANO5 | chr11:22239682 | T>G | 0 | 0 | No | No | Yes | Yes |
| 4746214 | FKRP | chr19:46756835 | C>G | 0 | 1 | Yes | Yes | No | No |

All five variants carried positive-class ClinVar labels and were misclassified by the twelve original 101-bp sequence configurations. The 201-bp and 501-bp columns report the number of correct classifications among six frozen models at each context length. Native-score correctness was determined using the respective training-derived thresholds. Two variants remained incorrect across all 24 sequence configurations, and one remained incorrect across all 26 configurations. These counts describe retrospective classification outcomes using known benchmark labels.

#### Supplementary Table S6. Recurrently difficult variants and retrospective outcomes (n = 787)

Recurrent-hard status means misclassified by at least four of the 12 original 101-bp configurations. The first panel summarizes the 787 variants by gene; the complete 787-variant listing follows. The accompanying full-width TSV preserves all 92 original source fields.

| Gene | Total | Positive | Negative | At least 1 201 bp correct | At least 1 501 bp correct | At least 1 native correct | Any later correct |
|---|---|---|---|---|---|---|---|
| ANO5 | 68 | 28 | 40 | 63 | 65 | 66 | 67 |
| CAPN3 | 125 | 68 | 57 | 111 | 117 | 125 | 125 |
| DNAJB6 | 19 | 1 | 18 | 19 | 19 | 19 | 19 |
| DYSF | 268 | 119 | 149 | 250 | 256 | 262 | 268 |
| FKRP | 48 | 17 | 31 | 39 | 46 | 48 | 48 |
| LMNA | 129 | 87 | 42 | 116 | 116 | 128 | 129 |
| SGCA | 53 | 33 | 20 | 40 | 43 | 47 | 49 |
| SGCB | 38 | 20 | 18 | 32 | 31 | 38 | 38 |
| SGCG | 31 | 8 | 23 | 31 | 30 | 31 | 31 |
| TCAP | 8 | 4 | 4 | 8 | 8 | 8 | 8 |
| TOTAL | 787 | 385 | 402 | 709 | 731 | 772 | 782 |

The additional strategy groups overlap in the variants they classify correctly. Five of the 787 variants were misclassified by all fourteen additional configurations: SGCA VariationIDs 188733, 197094, 498275 and 3384982, and ANO5 VariationID 4538462. All five carried positive-class benchmark labels. Each SGCA variant was correctly classified by three of the original twelve 101-bp configurations. The ANO5 variant was the only case misclassified across all 26 configurations.

#### Supplementary Table S6 (continued). Complete variant-level records

Columns: 101-bp errors among 12 original frozen/adapted configurations; number correctly classified at 201 and 501 bp among six frozen models per window; native correct among two native scores; total erroneous predictions among all 26 configurations; and correct under any later strategy. Label is the aggregate ClinVar classification, not LGMD-specific causality.

| VariationID | Gene | Label | 101 bp wrong /12 | 201 bp correct /6 | 501 bp correct /6 | Native correct /2 | Incorrect predictions /26 | Any later correct |
|---|---|---|---|---|---|---|---|---|
| 2163 | ANO5 | P | 12 | 0 | 0 | 2 | 24 | Yes |
| 2165 | ANO5 | P | 8 | 1 | 1 | 2 | 18 | Yes |
| 96679 | ANO5 | P | 7 | 1 | 1 | 2 | 17 | Yes |
| 140544 | ANO5 | B | 5 | 6 | 6 | 2 | 5 | Yes |
| 140546 | ANO5 | B | 5 | 6 | 6 | 2 | 5 | Yes |
| 140555 | ANO5 | P | 7 | 1 | 1 | 2 | 17 | Yes |
| 140556 | ANO5 | B | 5 | 6 | 6 | 2 | 5 | Yes |
| 194688 | ANO5 | P | 6 | 1 | 1 | 2 | 16 | Yes |
| 194805 | ANO5 | P | 5 | 0 | 1 | 2 | 16 | Yes |
| 263307 | ANO5 | B | 5 | 6 | 6 | 2 | 5 | Yes |
| 304130 | ANO5 | B | 4 | 5 | 5 | 2 | 6 | Yes |
| 431848 | ANO5 | P | 8 | 1 | 0 | 2 | 19 | Yes |
| 502370 | ANO5 | P | 9 | 0 | 1 | 2 | 20 | Yes |
| 502588 | ANO5 | P | 5 | 1 | 1 | 2 | 15 | Yes |
| 523571 | ANO5 | P | 7 | 1 | 1 | 2 | 17 | Yes |
| 536729 | ANO5 | P | 7 | 3 | 2 | 2 | 14 | Yes |

| VariationID | Gene | Label | 101 bp wrong /12 | 201 bp correct /6 | 501 bp correct /6 | Native correct /2 | Incorrect predictions /26 | Any later correct |
|---|---|---|---|---|---|---|---|---|
| 631656 | ANO5 | P | 9 | 0 | 1 | 2 | 20 | Yes |
| 650336 | ANO5 | P | 6 | 4 | 2 | 2 | 12 | Yes |
| 673918 | ANO5 | B | 6 | 6 | 6 | 2 | 6 | Yes |
| 681444 | ANO5 | B | 4 | 6 | 6 | 2 | 4 | Yes |
| 694036 | ANO5 | P | 6 | 2 | 1 | 2 | 15 | Yes |
| 694040 | ANO5 | P | 9 | 2 | 1 | 1 | 19 | Yes |
| 705208 | ANO5 | B | 5 | 6 | 6 | 2 | 5 | Yes |
| 813973 | ANO5 | P | 7 | 1 | 1 | 2 | 17 | Yes |
| 877787 | ANO5 | B | 4 | 6 | 6 | 2 | 4 | Yes |
| 1002707 | ANO5 | P | 9 | 2 | 1 | 2 | 18 | Yes |
| 1029543 | ANO5 | P | 5 | 2 | 1 | 2 | 14 | Yes |
| 1082199 | ANO5 | B | 8 | 6 | 5 | 2 | 9 | Yes |
| 1093464 | ANO5 | B | 4 | 6 | 5 | 2 | 5 | Yes |
| 1108719 | ANO5 | B | 4 | 6 | 6 | 2 | 4 | Yes |
| 1142933 | ANO5 | B | 5 | 5 | 6 | 2 | 6 | Yes |
| 1199476 | ANO5 | B | 4 | 6 | 6 | 2 | 4 | Yes |
| 1202441 | ANO5 | B | 8 | 4 | 6 | 2 | 10 | Yes |
| 1202786 | ANO5 | B | 4 | 6 | 6 | 2 | 4 | Yes |
| 1217635 | ANO5 | B | 7 | 6 | 6 | 2 | 7 | Yes |
| 1234963 | ANO5 | B | 6 | 6 | 6 | 2 | 6 | Yes |
| 1393170 | ANO5 | B | 4 | 6 | 6 | 2 | 4 | Yes |
| 1452790 | ANO5 | P | 5 | 1 | 1 | 2 | 15 | Yes |
| 1460312 | ANO5 | P | 6 | 1 | 1 | 2 | 16 | Yes |
| 1494665 | ANO5 | B | 4 | 6 | 6 | 2 | 4 | Yes |
| 1585949 | ANO5 | B | 5 | 5 | 6 | 2 | 6 | Yes |
| 1593728 | ANO5 | B | 5 | 5 | 6 | 2 | 6 | Yes |
| 1611701 | ANO5 | B | 6 | 5 | 6 | 2 | 7 | Yes |
| 1708996 | ANO5 | P | 5 | 2 | 4 | 2 | 11 | Yes |
| 1936524 | ANO5 | B | 6 | 5 | 6 | 2 | 7 | Yes |
| 1976594 | ANO5 | B | 4 | 5 | 6 | 2 | 5 | Yes |
| 2043922 | ANO5 | P | 7 | 1 | 1 | 2 | 17 | Yes |
| 2074063 | ANO5 | B | 4 | 6 | 6 | 2 | 4 | Yes |
| 2082146 | ANO5 | B | 6 | 6 | 6 | 2 | 6 | Yes |
| 2115016 | ANO5 | P | 6 | 2 | 2 | 2 | 14 | Yes |
| 2137032 | ANO5 | P | 10 | 1 | 1 | 0 | 22 | Yes |
| 2148056 | ANO5 | B | 4 | 6 | 6 | 2 | 4 | Yes |
| 2434185 | ANO5 | P | 6 | 2 | 2 | 1 | 15 | Yes |
| 2933794 | ANO5 | B | 6 | 6 | 6 | 2 | 6 | Yes |
| 2935168 | ANO5 | P | 9 | 2 | 1 | 2 | 18 | Yes |
| 2936455 | ANO5 | B | 5 | 6 | 6 | 2 | 5 | Yes |
| 2937035 | ANO5 | B | 5 | 6 | 6 | 2 | 5 | Yes |
| 2939316 | ANO5 | B | 6 | 5 | 6 | 2 | 7 | Yes |
| 2949925 | ANO5 | B | 6 | 5 | 5 | 1 | 9 | Yes |
| 3337803 | ANO5 | P | 6 | 1 | 1 | 2 | 16 | Yes |
| 3756322 | ANO5 | B | 4 | 5 | 6 | 2 | 5 | Yes |
| 3758308 | ANO5 | B | 7 | 4 | 6 | 2 | 9 | Yes |
| 3761419 | ANO5 | B | 5 | 6 | 6 | 2 | 5 | Yes |
| 3763433 | ANO5 | B | 4 | 5 | 6 | 2 | 5 | Yes |

| VariationID | Gene | Label | 101 bp wrong /12 | 201 bp correct /6 | 501 bp correct /6 | Native correct /2 | Incorrect predictions /26 | Any later correct |
|---|---|---|---|---|---|---|---|---|
| 4538462 | ANO5 | P | 12 | 0 | 0 | 0 | 26 | No |
| 4786753 | ANO5 | B | 4 | 6 | 6 | 2 | 4 | Yes |
| 4792793 | ANO5 | B | 5 | 6 | 6 | 2 | 5 | Yes |
| 4793232 | ANO5 | B | 5 | 4 | 4 | 2 | 9 | Yes |
| 17619 | CAPN3 | P | 7 | 1 | 1 | 2 | 17 | Yes |
| 128570 | CAPN3 | P | 9 | 2 | 2 | 2 | 17 | Yes |
| 128575 | CAPN3 | B | 6 | 6 | 6 | 2 | 6 | Yes |
| 166793 | CAPN3 | B | 6 | 5 | 6 | 2 | 7 | Yes |
| 195011 | CAPN3 | B | 6 | 6 | 6 | 2 | 6 | Yes |
| 198690 | CAPN3 | P | 7 | 1 | 1 | 2 | 17 | Yes |
| 217146 | CAPN3 | P | 11 | 0 | 0 | 1 | 24 | Yes |
| 217148 | CAPN3 | P | 6 | 0 | 2 | 2 | 16 | Yes |
| 217152 | CAPN3 | P | 6 | 1 | 2 | 2 | 15 | Yes |
| 217157 | CAPN3 | P | 5 | 4 | 1 | 2 | 12 | Yes |
| 254872 | CAPN3 | B | 5 | 5 | 6 | 2 | 6 | Yes |
| 280038 | CAPN3 | P | 6 | 2 | 0 | 2 | 16 | Yes |
| 280039 | CAPN3 | P | 7 | 1 | 1 | 2 | 17 | Yes |
| 281254 | CAPN3 | P | 7 | 1 | 1 | 2 | 17 | Yes |
| 282646 | CAPN3 | P | 5 | 2 | 1 | 1 | 15 | Yes |
| 282783 | CAPN3 | P | 8 | 2 | 1 | 2 | 17 | Yes |
| 283099 | CAPN3 | P | 7 | 1 | 2 | 2 | 16 | Yes |
| 283991 | CAPN3 | P | 5 | 1 | 2 | 2 | 14 | Yes |
| 284946 | CAPN3 | P | 5 | 3 | 2 | 2 | 12 | Yes |
| 285340 | CAPN3 | P | 10 | 0 | 1 | 2 | 21 | Yes |
| 285460 | CAPN3 | P | 8 | 1 | 1 | 2 | 18 | Yes |
| 286592 | CAPN3 | P | 7 | 3 | 3 | 1 | 14 | Yes |
| 426275 | CAPN3 | P | 9 | 0 | 1 | 1 | 21 | Yes |
| 468639 | CAPN3 | P | 7 | 1 | 1 | 2 | 17 | Yes |
| 496977 | CAPN3 | P | 10 | 0 | 0 | 2 | 22 | Yes |
| 497565 | CAPN3 | P | 8 | 1 | 1 | 2 | 18 | Yes |
| 497833 | CAPN3 | P | 5 | 0 | 1 | 2 | 16 | Yes |
| 498205 | CAPN3 | P | 6 | 2 | 1 | 2 | 15 | Yes |
| 499202 | CAPN3 | P | 5 | 6 | 5 | 1 | 7 | Yes |
| 552636 | CAPN3 | P | 7 | 0 | 2 | 1 | 18 | Yes |
| 552843 | CAPN3 | P | 6 | 4 | 3 | 2 | 11 | Yes |
| 553345 | CAPN3 | P | 5 | 3 | 4 | 2 | 10 | Yes |
| 553662 | CAPN3 | P | 6 | 1 | 1 | 2 | 16 | Yes |
| 554405 | CAPN3 | P | 9 | 1 | 1 | 2 | 19 | Yes |
| 555599 | CAPN3 | P | 8 | 2 | 2 | 1 | 17 | Yes |
| 632230 | CAPN3 | P | 6 | 2 | 2 | 2 | 14 | Yes |
| 661943 | CAPN3 | P | 7 | 1 | 1 | 2 | 17 | Yes |
| 680087 | CAPN3 | B | 5 | 6 | 6 | 2 | 5 | Yes |
| 704728 | CAPN3 | B | 5 | 6 | 6 | 2 | 5 | Yes |
| 772921 | CAPN3 | B | 5 | 6 | 6 | 2 | 5 | Yes |
| 796305 | CAPN3 | B | 7 | 4 | 6 | 2 | 9 | Yes |
| 843051 | CAPN3 | P | 11 | 1 | 1 | 2 | 21 | Yes |
| 959503 | CAPN3 | P | 8 | 1 | 1 | 2 | 18 | Yes |
| 983852 | CAPN3 | P | 4 | 3 | 4 | 2 | 9 | Yes |

| VariationID | Gene | Label | 101 bp wrong /12 | 201 bp correct /6 | 501 bp correct /6 | Native correct /2 | Incorrect predictions /26 | Any later correct |
|---|---|---|---|---|---|---|---|---|
| 983854 | CAPN3 | P | 6 | 1 | 2 | 2 | 15 | Yes |
| 984013 | CAPN3 | P | 7 | 2 | 1 | 2 | 16 | Yes |
| 984014 | CAPN3 | P | 4 | 2 | 1 | 1 | 14 | Yes |
| 990239 | CAPN3 | B | 6 | 6 | 6 | 2 | 6 | Yes |
| 1009814 | CAPN3 | P | 6 | 1 | 1 | 2 | 16 | Yes |
| 1079698 | CAPN3 | B | 4 | 6 | 6 | 2 | 4 | Yes |
| 1087154 | CAPN3 | B | 5 | 6 | 6 | 2 | 5 | Yes |
| 1088268 | CAPN3 | B | 4 | 6 | 5 | 2 | 5 | Yes |
| 1096376 | CAPN3 | B | 4 | 5 | 6 | 2 | 5 | Yes |
| 1120684 | CAPN3 | B | 4 | 6 | 6 | 2 | 4 | Yes |
| 1136652 | CAPN3 | B | 5 | 6 | 6 | 2 | 5 | Yes |
| 1140997 | CAPN3 | B | 5 | 4 | 6 | 2 | 7 | Yes |
| 1143363 | CAPN3 | B | 5 | 4 | 6 | 2 | 7 | Yes |
| 1180649 | CAPN3 | P | 8 | 0 | 1 | 2 | 19 | Yes |
| 1196273 | CAPN3 | B | 4 | 5 | 6 | 2 | 5 | Yes |
| 1214104 | CAPN3 | B | 6 | 6 | 6 | 2 | 6 | Yes |
| 1246204 | CAPN3 | B | 5 | 5 | 6 | 2 | 6 | Yes |
| 1321461 | CAPN3 | P | 10 | 1 | 1 | 2 | 20 | Yes |
| 1333344 | CAPN3 | P | 6 | 3 | 1 | 2 | 14 | Yes |
| 1345765 | CAPN3 | P | 6 | 2 | 2 | 2 | 14 | Yes |
| 1453744 | CAPN3 | P | 7 | 0 | 1 | 2 | 18 | Yes |
| 1458711 | CAPN3 | P | 6 | 1 | 2 | 2 | 15 | Yes |
| 1493433 | CAPN3 | P | 7 | 1 | 1 | 2 | 17 | Yes |
| 1596784 | CAPN3 | B | 5 | 6 | 6 | 2 | 5 | Yes |
| 1633605 | CAPN3 | B | 4 | 6 | 6 | 2 | 4 | Yes |
| 1656691 | CAPN3 | B | 4 | 5 | 6 | 2 | 5 | Yes |
| 1923288 | CAPN3 | B | 4 | 6 | 6 | 2 | 4 | Yes |
| 2002126 | CAPN3 | B | 5 | 6 | 6 | 2 | 5 | Yes |
| 2018320 | CAPN3 | B | 8 | 5 | 6 | 2 | 9 | Yes |
| 2018852 | CAPN3 | B | 4 | 6 | 6 | 2 | 4 | Yes |
| 2028963 | CAPN3 | B | 6 | 6 | 6 | 2 | 6 | Yes |
| 2081612 | CAPN3 | B | 4 | 5 | 6 | 2 | 5 | Yes |
| 2109761 | CAPN3 | P | 8 | 0 | 0 | 2 | 20 | Yes |
| 2137655 | CAPN3 | P | 9 | 1 | 1 | 2 | 19 | Yes |
| 2152226 | CAPN3 | P | 6 | 3 | 2 | 2 | 13 | Yes |
| 2164312 | CAPN3 | B | 8 | 6 | 4 | 2 | 10 | Yes |
| 2197845 | CAPN3 | B | 6 | 5 | 4 | 2 | 9 | Yes |
| 2444303 | CAPN3 | P | 5 | 1 | 0 | 2 | 16 | Yes |
| 2680376 | CAPN3 | P | 10 | 0 | 0 | 1 | 23 | Yes |
| 2680388 | CAPN3 | P | 7 | 2 | 3 | 1 | 15 | Yes |
| 2719496 | CAPN3 | B | 4 | 5 | 6 | 2 | 5 | Yes |
| 2753825 | CAPN3 | B | 7 | 5 | 5 | 2 | 9 | Yes |
| 2797195 | CAPN3 | B | 6 | 4 | 6 | 2 | 8 | Yes |
| 2813736 | CAPN3 | B | 6 | 6 | 5 | 2 | 7 | Yes |
| 2829830 | CAPN3 | B | 7 | 5 | 6 | 2 | 8 | Yes |
| 2846050 | CAPN3 | P | 4 | 3 | 3 | 2 | 10 | Yes |
| 2855012 | CAPN3 | B | 6 | 5 | 6 | 2 | 7 | Yes |
| 2864420 | CAPN3 | B | 5 | 5 | 5 | 2 | 7 | Yes |

| VariationID | Gene | Label | 101 bp wrong /12 | 201 bp correct /6 | 501 bp correct /6 | Native correct /2 | Incorrect predictions /26 | Any later correct |
|---|---|---|---|---|---|---|---|---|
| 2875295 | CAPN3 | B | 7 | 4 | 5 | 2 | 10 | Yes |
| 2879962 | CAPN3 | B | 4 | 6 | 5 | 2 | 5 | Yes |
| 2883766 | CAPN3 | B | 4 | 6 | 6 | 2 | 4 | Yes |
| 2886934 | CAPN3 | B | 5 | 6 | 5 | 2 | 6 | Yes |
| 2897054 | CAPN3 | B | 4 | 6 | 6 | 2 | 4 | Yes |
| 2899337 | CAPN3 | B | 6 | 6 | 5 | 2 | 7 | Yes |
| 2911289 | CAPN3 | B | 5 | 6 | 6 | 2 | 5 | Yes |
| 2912556 | CAPN3 | B | 9 | 5 | 5 | 2 | 11 | Yes |
| 2915682 | CAPN3 | B | 6 | 5 | 6 | 2 | 7 | Yes |
| 2919990 | CAPN3 | B | 4 | 6 | 6 | 2 | 4 | Yes |
| 2971363 | CAPN3 | B | 6 | 4 | 6 | 2 | 8 | Yes |
| 2983378 | CAPN3 | B | 5 | 6 | 6 | 2 | 5 | Yes |
| 3015145 | CAPN3 | B | 5 | 6 | 6 | 2 | 5 | Yes |
| 3063866 | CAPN3 | P | 11 | 1 | 1 | 2 | 21 | Yes |
| 3240997 | CAPN3 | P | 4 | 1 | 1 | 2 | 14 | Yes |
| 3577109 | CAPN3 | P | 7 | 3 | 1 | 2 | 15 | Yes |
| 3577111 | CAPN3 | P | 6 | 4 | 3 | 2 | 11 | Yes |
| 3577112 | CAPN3 | P | 6 | 3 | 2 | 2 | 13 | Yes |
| 3577113 | CAPN3 | P | 6 | 0 | 0 | 2 | 18 | Yes |
| 3678358 | CAPN3 | P | 6 | 3 | 2 | 2 | 13 | Yes |
| 3721675 | CAPN3 | P | 8 | 2 | 2 | 2 | 16 | Yes |
| 3728719 | CAPN3 | B | 5 | 6 | 6 | 2 | 5 | Yes |
| 4077714 | CAPN3 | P | 8 | 0 | 2 | 2 | 18 | Yes |
| 4081229 | CAPN3 | P | 9 | 2 | 3 | 1 | 17 | Yes |
| 4690879 | CAPN3 | P | 5 | 2 | 1 | 1 | 15 | Yes |
| 4699870 | CAPN3 | B | 5 | 6 | 6 | 2 | 5 | Yes |
| 4706030 | CAPN3 | B | 6 | 6 | 6 | 2 | 6 | Yes |
| 4708140 | CAPN3 | B | 4 | 6 | 6 | 2 | 4 | Yes |
| 4720340 | CAPN3 | B | 5 | 6 | 6 | 2 | 5 | Yes |
| 4727509 | CAPN3 | B | 5 | 5 | 6 | 2 | 6 | Yes |
| 4816059 | CAPN3 | P | 9 | 0 | 0 | 2 | 21 | Yes |
| 4816171 | CAPN3 | P | 7 | 1 | 4 | 2 | 14 | Yes |
| 4838710 | CAPN3 | P | 5 | 1 | 1 | 2 | 15 | Yes |
| 30904 | DNAJB6 | P | 5 | 1 | 2 | 2 | 14 | Yes |
| 262304 | DNAJB6 | B | 6 | 5 | 5 | 2 | 8 | Yes |
| 262305 | DNAJB6 | B | 5 | 6 | 6 | 2 | 5 | Yes |
| 359455 | DNAJB6 | B | 5 | 6 | 5 | 2 | 6 | Yes |
| 359475 | DNAJB6 | B | 4 | 6 | 5 | 2 | 5 | Yes |
| 359478 | DNAJB6 | B | 5 | 6 | 6 | 2 | 5 | Yes |
| 385378 | DNAJB6 | B | 4 | 6 | 6 | 2 | 4 | Yes |
| 1019617 | DNAJB6 | B | 6 | 5 | 6 | 2 | 7 | Yes |
| 1105418 | DNAJB6 | B | 5 | 6 | 6 | 2 | 5 | Yes |
| 1298062 | DNAJB6 | B | 4 | 6 | 6 | 2 | 4 | Yes |
| 1554497 | DNAJB6 | B | 5 | 6 | 6 | 2 | 5 | Yes |
| 1585909 | DNAJB6 | B | 6 | 6 | 6 | 2 | 6 | Yes |
| 1594563 | DNAJB6 | B | 4 | 6 | 5 | 2 | 5 | Yes |
| 1604629 | DNAJB6 | B | 4 | 6 | 6 | 2 | 4 | Yes |
| 2195414 | DNAJB6 | B | 7 | 6 | 6 | 2 | 7 | Yes |

| VariationID | Gene | Label | 101 bp wrong /12 | 201 bp correct /6 | 501 bp correct /6 | Native correct /2 | Incorrect predictions /26 | Any later correct |
|---|---|---|---|---|---|---|---|---|
| 2970581 | DNAJB6 | B | 4 | 6 | 6 | 2 | 4 | Yes |
| 4694091 | DNAJB6 | B | 7 | 5 | 6 | 2 | 8 | Yes |
| 4714506 | DNAJB6 | B | 6 | 6 | 6 | 2 | 6 | Yes |
| 4781211 | DNAJB6 | B | 4 | 5 | 6 | 2 | 5 | Yes |
| 6681 | DYSF | P | 10 | 0 | 1 | 2 | 21 | Yes |
| 6682 | DYSF | P | 7 | 0 | 1 | 2 | 18 | Yes |
| 18443 | DYSF | P | 5 | 1 | 2 | 1 | 15 | Yes |
| 94269 | DYSF | P | 5 | 1 | 1 | 2 | 15 | Yes |
| 94270 | DYSF | B | 8 | 6 | 5 | 0 | 11 | Yes |
| 94273 | DYSF | P | 9 | 2 | 2 | 2 | 17 | Yes |
| 94274 | DYSF | P | 4 | 1 | 1 | 2 | 14 | Yes |
| 94278 | DYSF | P | 7 | 2 | 2 | 2 | 15 | Yes |
| 94290 | DYSF | B | 7 | 6 | 6 | 2 | 7 | Yes |
| 94321 | DYSF | P | 9 | 1 | 1 | 2 | 19 | Yes |
| 94340 | DYSF | P | 7 | 0 | 0 | 2 | 19 | Yes |
| 94341 | DYSF | B | 7 | 6 | 6 | 2 | 7 | Yes |
| 94346 | DYSF | B | 4 | 6 | 6 | 2 | 4 | Yes |
| 94365 | DYSF | P | 7 | 2 | 1 | 2 | 16 | Yes |
| 167021 | DYSF | P | 7 | 0 | 2 | 2 | 17 | Yes |
| 191111 | DYSF | P | 8 | 0 | 1 | 1 | 20 | Yes |
| 195490 | DYSF | P | 7 | 2 | 2 | 2 | 15 | Yes |
| 196694 | DYSF | P | 7 | 2 | 1 | 2 | 16 | Yes |
| 197217 | DYSF | P | 5 | 2 | 3 | 2 | 12 | Yes |
| 217224 | DYSF | P | 10 | 2 | 2 | 2 | 18 | Yes |
| 217227 | DYSF | P | 10 | 1 | 0 | 1 | 22 | Yes |
| 242418 | DYSF | P | 6 | 2 | 2 | 2 | 14 | Yes |
| 242527 | DYSF | P | 8 | 0 | 0 | 2 | 20 | Yes |
| 259080 | DYSF | B | 6 | 5 | 5 | 2 | 8 | Yes |
| 265487 | DYSF | P | 7 | 3 | 1 | 2 | 15 | Yes |
| 280068 | DYSF | P | 6 | 1 | 1 | 2 | 16 | Yes |
| 281067 | DYSF | P | 10 | 0 | 0 | 2 | 22 | Yes |
| 281197 | DYSF | P | 11 | 1 | 0 | 0 | 24 | Yes |
| 282164 | DYSF | B | 4 | 6 | 6 | 2 | 4 | Yes |
| 282408 | DYSF | P | 6 | 1 | 0 | 2 | 17 | Yes |
| 282624 | DYSF | P | 8 | 0 | 0 | 2 | 20 | Yes |
| 283206 | DYSF | P | 8 | 0 | 0 | 2 | 20 | Yes |
| 283471 | DYSF | P | 7 | 4 | 2 | 2 | 13 | Yes |
| 283497 | DYSF | P | 6 | 3 | 2 | 2 | 13 | Yes |
| 284882 | DYSF | P | 6 | 1 | 3 | 2 | 14 | Yes |
| 285584 | DYSF | P | 5 | 5 | 3 | 1 | 10 | Yes |
| 285590 | DYSF | P | 8 | 1 | 2 | 2 | 17 | Yes |
| 285665 | DYSF | P | 6 | 1 | 1 | 2 | 16 | Yes |
| 286743 | DYSF | P | 6 | 1 | 2 | 2 | 15 | Yes |
| 288008 | DYSF | B | 6 | 5 | 4 | 1 | 10 | Yes |
| 288183 | DYSF | P | 8 | 1 | 1 | 2 | 18 | Yes |
| 288860 | DYSF | P | 7 | 1 | 1 | 2 | 17 | Yes |
| 290202 | DYSF | P | 9 | 1 | 2 | 2 | 18 | Yes |
| 290515 | DYSF | P | 5 | 2 | 1 | 2 | 14 | Yes |

| VariationID | Gene | Label | 101 bp wrong /12 | 201 bp correct /6 | 501 bp correct /6 | Native correct /2 | Incorrect predictions /26 | Any later correct |
|---|---|---|---|---|---|---|---|---|
| 381677 | DYSF | P | 8 | 1 | 1 | 2 | 18 | Yes |
| 444503 | DYSF | B | 5 | 6 | 6 | 2 | 5 | Yes |
| 471307 | DYSF | P | 7 | 2 | 1 | 2 | 16 | Yes |
| 471318 | DYSF | P | 8 | 1 | 0 | 2 | 19 | Yes |
| 498954 | DYSF | P | 6 | 1 | 2 | 2 | 15 | Yes |
| 500678 | DYSF | P | 7 | 1 | 0 | 2 | 18 | Yes |
| 500814 | DYSF | P | 9 | 0 | 1 | 2 | 20 | Yes |
| 510044 | DYSF | B | 4 | 6 | 6 | 2 | 4 | Yes |
| 513657 | DYSF | B | 8 | 4 | 6 | 2 | 10 | Yes |
| 518129 | DYSF | B | 5 | 5 | 5 | 2 | 7 | Yes |
| 538655 | DYSF | B | 5 | 5 | 6 | 2 | 6 | Yes |
| 551891 | DYSF | P | 8 | 1 | 1 | 2 | 18 | Yes |
| 552023 | DYSF | P | 6 | 2 | 2 | 2 | 14 | Yes |
| 555456 | DYSF | P | 7 | 1 | 1 | 2 | 17 | Yes |
| 555770 | DYSF | P | 7 | 1 | 1 | 2 | 17 | Yes |
| 555783 | DYSF | P | 5 | 2 | 1 | 2 | 14 | Yes |
| 556639 | DYSF | P | 5 | 3 | 5 | 2 | 9 | Yes |
| 565629 | DYSF | P | 4 | 3 | 1 | 2 | 12 | Yes |
| 574900 | DYSF | P | 5 | 4 | 3 | 2 | 10 | Yes |
| 596406 | DYSF | P | 5 | 3 | 4 | 2 | 10 | Yes |
| 619023 | DYSF | P | 10 | 0 | 1 | 2 | 21 | Yes |
| 635045 | DYSF | P | 4 | 4 | 3 | 2 | 9 | Yes |
| 655896 | DYSF | P | 6 | 0 | 2 | 2 | 16 | Yes |
| 658470 | DYSF | P | 5 | 3 | 2 | 2 | 12 | Yes |
| 668121 | DYSF | B | 5 | 5 | 5 | 2 | 7 | Yes |
| 668125 | DYSF | B | 4 | 6 | 6 | 2 | 4 | Yes |
| 670396 | DYSF | B | 5 | 6 | 6 | 2 | 5 | Yes |
| 670490 | DYSF | B | 4 | 6 | 6 | 2 | 4 | Yes |
| 671764 | DYSF | B | 6 | 5 | 6 | 2 | 7 | Yes |
| 677301 | DYSF | B | 6 | 5 | 6 | 2 | 7 | Yes |
| 678253 | DYSF | B | 5 | 6 | 6 | 2 | 5 | Yes |
| 679856 | DYSF | B | 6 | 6 | 3 | 2 | 9 | Yes |
| 681729 | DYSF | B | 4 | 6 | 6 | 2 | 4 | Yes |
| 681743 | DYSF | B | 4 | 6 | 6 | 2 | 4 | Yes |
| 694000 | DYSF | P | 6 | 4 | 1 | 2 | 13 | Yes |
| 741023 | DYSF | B | 4 | 5 | 5 | 2 | 6 | Yes |
| 748572 | DYSF | B | 4 | 5 | 5 | 2 | 6 | Yes |
| 754491 | DYSF | B | 4 | 6 | 6 | 2 | 4 | Yes |
| 792230 | DYSF | B | 5 | 6 | 6 | 2 | 5 | Yes |
| 797824 | DYSF | B | 7 | 6 | 6 | 2 | 7 | Yes |
| 804770 | DYSF | P | 6 | 1 | 1 | 2 | 16 | Yes |
| 837557 | DYSF | P | 6 | 1 | 0 | 2 | 17 | Yes |
| 837580 | DYSF | P | 6 | 2 | 2 | 2 | 14 | Yes |
| 838510 | DYSF | P | 6 | 1 | 2 | 2 | 15 | Yes |
| 850419 | DYSF | P | 5 | 3 | 2 | 2 | 12 | Yes |
| 857602 | DYSF | P | 6 | 1 | 2 | 1 | 16 | Yes |
| 858838 | DYSF | P | 6 | 1 | 1 | 2 | 16 | Yes |
| 869488 | DYSF | P | 7 | 1 | 1 | 2 | 17 | Yes |

| VariationID | Gene | Label | 101 bp wrong /12 | 201 bp correct /6 | 501 bp correct /6 | Native correct /2 | Incorrect predictions /26 | Any later correct |
|---|---|---|---|---|---|---|---|---|
| 938081 | DYSF | P | 5 | 2 | 2 | 2 | 13 | Yes |
| 960932 | DYSF | P | 8 | 1 | 1 | 2 | 18 | Yes |
| 983670 | DYSF | P | 8 | 4 | 4 | 2 | 12 | Yes |
| 983675 | DYSF | P | 6 | 1 | 3 | 2 | 14 | Yes |
| 983957 | DYSF | P | 5 | 3 | 2 | 2 | 12 | Yes |
| 983963 | DYSF | P | 7 | 2 | 2 | 2 | 15 | Yes |
| 984128 | DYSF | P | 6 | 2 | 1 | 2 | 15 | Yes |
| 984229 | DYSF | P | 6 | 2 | 2 | 2 | 14 | Yes |
| 984234 | DYSF | P | 7 | 2 | 2 | 2 | 15 | Yes |
| 984236 | DYSF | P | 9 | 2 | 1 | 2 | 18 | Yes |
| 984245 | DYSF | P | 6 | 3 | 3 | 2 | 12 | Yes |
| 984248 | DYSF | P | 7 | 0 | 1 | 2 | 18 | Yes |
| 1072189 | DYSF | P | 5 | 3 | 2 | 2 | 12 | Yes |
| 1074366 | DYSF | P | 6 | 2 | 3 | 2 | 13 | Yes |
| 1078347 | DYSF | B | 7 | 4 | 6 | 2 | 9 | Yes |
| 1078931 | DYSF | B | 6 | 6 | 6 | 2 | 6 | Yes |
| 1087457 | DYSF | B | 5 | 6 | 6 | 2 | 5 | Yes |
| 1087469 | DYSF | B | 5 | 4 | 5 | 2 | 8 | Yes |
| 1091559 | DYSF | B | 4 | 6 | 6 | 2 | 4 | Yes |
| 1092336 | DYSF | B | 5 | 5 | 6 | 2 | 6 | Yes |
| 1101849 | DYSF | B | 6 | 6 | 6 | 2 | 6 | Yes |
| 1110070 | DYSF | B | 5 | 6 | 5 | 2 | 6 | Yes |
| 1115420 | DYSF | B | 4 | 6 | 6 | 2 | 4 | Yes |
| 1124582 | DYSF | B | 4 | 5 | 5 | 2 | 6 | Yes |
| 1125807 | DYSF | B | 6 | 4 | 4 | 2 | 10 | Yes |
| 1128787 | DYSF | B | 5 | 6 | 6 | 2 | 5 | Yes |
| 1129236 | DYSF | B | 5 | 4 | 4 | 2 | 9 | Yes |
| 1129323 | DYSF | B | 9 | 3 | 2 | 2 | 16 | Yes |
| 1129731 | DYSF | B | 5 | 4 | 6 | 2 | 7 | Yes |
| 1131985 | DYSF | B | 8 | 4 | 5 | 2 | 11 | Yes |
| 1136822 | DYSF | B | 5 | 4 | 5 | 2 | 8 | Yes |
| 1137850 | DYSF | B | 5 | 6 | 6 | 2 | 5 | Yes |
| 1155479 | DYSF | B | 4 | 6 | 6 | 2 | 4 | Yes |
| 1159236 | DYSF | B | 5 | 6 | 6 | 2 | 5 | Yes |
| 1162151 | DYSF | B | 5 | 5 | 5 | 2 | 7 | Yes |
| 1179998 | DYSF | B | 5 | 6 | 6 | 2 | 5 | Yes |
| 1180715 | DYSF | P | 5 | 2 | 3 | 2 | 12 | Yes |
| 1188437 | DYSF | B | 4 | 5 | 6 | 2 | 5 | Yes |
| 1190225 | DYSF | B | 4 | 6 | 6 | 2 | 4 | Yes |
| 1195459 | DYSF | B | 5 | 6 | 6 | 2 | 5 | Yes |
| 1200752 | DYSF | B | 6 | 6 | 6 | 2 | 6 | Yes |
| 1204262 | DYSF | B | 6 | 5 | 6 | 2 | 7 | Yes |
| 1210571 | DYSF | B | 4 | 6 | 6 | 2 | 4 | Yes |
| 1214122 | DYSF | B | 5 | 6 | 6 | 2 | 5 | Yes |
| 1216261 | DYSF | B | 6 | 5 | 6 | 2 | 7 | Yes |
| 1220008 | DYSF | B | 4 | 6 | 6 | 2 | 4 | Yes |
| 1292308 | DYSF | B | 4 | 6 | 6 | 2 | 4 | Yes |
| 1292521 | DYSF | B | 6 | 5 | 5 | 2 | 8 | Yes |

| VariationID | Gene | Label | 101 bp wrong /12 | 201 bp correct /6 | 501 bp correct /6 | Native correct /2 | Incorrect predictions /26 | Any later correct |
|---|---|---|---|---|---|---|---|---|
| 1296222 | DYSF | B | 4 | 6 | 6 | 2 | 4 | Yes |
| 1298155 | DYSF | B | 4 | 5 | 6 | 2 | 5 | Yes |
| 1321153 | DYSF | P | 5 | 2 | 0 | 2 | 15 | Yes |
| 1322794 | DYSF | P | 4 | 1 | 3 | 2 | 12 | Yes |
| 1322801 | DYSF | P | 8 | 2 | 2 | 2 | 16 | Yes |
| 1322803 | DYSF | P | 5 | 4 | 3 | 2 | 10 | Yes |
| 1322810 | DYSF | P | 5 | 2 | 2 | 2 | 13 | Yes |
| 1326969 | DYSF | P | 4 | 3 | 3 | 2 | 10 | Yes |
| 1370138 | DYSF | B | 5 | 5 | 6 | 0 | 8 | Yes |
| 1444196 | DYSF | P | 9 | 1 | 1 | 2 | 19 | Yes |
| 1483857 | DYSF | P | 4 | 1 | 3 | 2 | 12 | Yes |
| 1528379 | DYSF | B | 4 | 5 | 6 | 2 | 5 | Yes |
| 1532130 | DYSF | B | 7 | 5 | 5 | 2 | 9 | Yes |
| 1537695 | DYSF | B | 7 | 5 | 5 | 2 | 9 | Yes |
| 1541840 | DYSF | B | 5 | 6 | 6 | 2 | 5 | Yes |
| 1542964 | DYSF | B | 4 | 5 | 6 | 2 | 5 | Yes |
| 1557386 | DYSF | B | 4 | 6 | 6 | 2 | 4 | Yes |
| 1560573 | DYSF | B | 5 | 5 | 6 | 2 | 6 | Yes |
| 1560740 | DYSF | B | 5 | 6 | 6 | 2 | 5 | Yes |
| 1563467 | DYSF | B | 4 | 6 | 6 | 2 | 4 | Yes |
| 1570567 | DYSF | B | 6 | 6 | 6 | 2 | 6 | Yes |
| 1577197 | DYSF | B | 6 | 5 | 5 | 2 | 8 | Yes |
| 1589325 | DYSF | B | 4 | 6 | 5 | 2 | 5 | Yes |
| 1590053 | DYSF | B | 7 | 5 | 5 | 2 | 9 | Yes |
| 1594114 | DYSF | B | 5 | 4 | 5 | 2 | 8 | Yes |
| 1607951 | DYSF | B | 5 | 6 | 6 | 2 | 5 | Yes |
| 1627507 | DYSF | B | 4 | 6 | 6 | 2 | 4 | Yes |
| 1643251 | DYSF | B | 5 | 5 | 6 | 2 | 6 | Yes |
| 1661184 | DYSF | B | 4 | 6 | 6 | 2 | 4 | Yes |
| 1662952 | DYSF | B | 4 | 6 | 6 | 2 | 4 | Yes |
| 1673123 | DYSF | B | 4 | 6 | 6 | 2 | 4 | Yes |
| 1707937 | DYSF | B | 5 | 5 | 6 | 2 | 6 | Yes |
| 1708553 | DYSF | P | 4 | 3 | 4 | 2 | 9 | Yes |
| 1711502 | DYSF | B | 4 | 6 | 6 | 2 | 4 | Yes |
| 1726282 | DYSF | P | 8 | 3 | 1 | 2 | 16 | Yes |
| 1915977 | DYSF | B | 4 | 6 | 6 | 2 | 4 | Yes |
| 1968746 | DYSF | P | 10 | 1 | 1 | 2 | 20 | Yes |
| 1974745 | DYSF | P | 6 | 2 | 1 | 2 | 15 | Yes |
| 1977927 | DYSF | B | 4 | 6 | 6 | 2 | 4 | Yes |
| 1982098 | DYSF | B | 4 | 5 | 5 | 0 | 8 | Yes |
| 1991591 | DYSF | P | 5 | 1 | 1 | 2 | 15 | Yes |
| 2038699 | DYSF | B | 5 | 6 | 5 | 2 | 6 | Yes |
| 2053855 | DYSF | B | 5 | 6 | 6 | 2 | 5 | Yes |
| 2055621 | DYSF | P | 6 | 2 | 1 | 2 | 15 | Yes |
| 2077391 | DYSF | B | 5 | 5 | 5 | 2 | 7 | Yes |
| 2097056 | DYSF | B | 5 | 6 | 6 | 2 | 5 | Yes |
| 2120408 | DYSF | B | 6 | 6 | 6 | 2 | 6 | Yes |
| 2120943 | DYSF | B | 4 | 6 | 6 | 2 | 4 | Yes |

| VariationID | Gene | Label | 101 bp wrong /12 | 201 bp correct /6 | 501 bp correct /6 | Native correct /2 | Incorrect predictions /26 | Any later correct |
|---|---|---|---|---|---|---|---|---|
| 2122551 | DYSF | B | 5 | 6 | 6 | 2 | 5 | Yes |
| 2145882 | DYSF | B | 5 | 4 | 6 | 2 | 7 | Yes |
| 2153164 | DYSF | P | 6 | 1 | 1 | 2 | 16 | Yes |
| 2154619 | DYSF | B | 4 | 5 | 6 | 1 | 6 | Yes |
| 2166693 | DYSF | B | 4 | 6 | 5 | 0 | 7 | Yes |
| 2170386 | DYSF | B | 5 | 6 | 6 | 2 | 5 | Yes |
| 2203084 | DYSF | P | 10 | 0 | 1 | 2 | 21 | Yes |
| 2203093 | DYSF | P | 5 | 2 | 1 | 2 | 14 | Yes |
| 2420891 | DYSF | B | 5 | 6 | 6 | 2 | 5 | Yes |
| 2570601 | DYSF | P | 5 | 1 | 1 | 2 | 15 | Yes |
| 2674970 | DYSF | P | 5 | 1 | 1 | 2 | 15 | Yes |
| 2674972 | DYSF | P | 8 | 0 | 1 | 2 | 19 | Yes |
| 2674991 | DYSF | P | 6 | 2 | 3 | 2 | 13 | Yes |
| 2674999 | DYSF | P | 5 | 2 | 2 | 2 | 13 | Yes |
| 2694320 | DYSF | B | 5 | 4 | 4 | 2 | 9 | Yes |
| 2698517 | DYSF | B | 4 | 6 | 6 | 2 | 4 | Yes |
| 2702519 | DYSF | B | 5 | 5 | 5 | 2 | 7 | Yes |
| 2703488 | DYSF | B | 4 | 6 | 6 | 2 | 4 | Yes |
| 2703538 | DYSF | P | 6 | 5 | 3 | 2 | 10 | Yes |
| 2711714 | DYSF | B | 4 | 6 | 6 | 2 | 4 | Yes |
| 2721157 | DYSF | B | 5 | 6 | 6 | 2 | 5 | Yes |
| 2724340 | DYSF | B | 5 | 4 | 5 | 1 | 9 | Yes |
| 2729657 | DYSF | B | 4 | 6 | 6 | 2 | 4 | Yes |
| 2756019 | DYSF | B | 7 | 6 | 6 | 2 | 7 | Yes |
| 2766730 | DYSF | P | 7 | 3 | 3 | 2 | 13 | Yes |
| 2769877 | DYSF | B | 4 | 6 | 6 | 2 | 4 | Yes |
| 2784113 | DYSF | B | 4 | 6 | 6 | 2 | 4 | Yes |
| 2787156 | DYSF | B | 6 | 5 | 6 | 2 | 7 | Yes |
| 2791794 | DYSF | B | 6 | 5 | 6 | 2 | 7 | Yes |
| 2827230 | DYSF | P | 7 | 1 | 2 | 2 | 16 | Yes |
| 2837659 | DYSF | B | 7 | 5 | 5 | 2 | 9 | Yes |
| 2838250 | DYSF | B | 6 | 5 | 6 | 2 | 7 | Yes |
| 2844670 | DYSF | B | 5 | 6 | 6 | 2 | 5 | Yes |
| 2845927 | DYSF | B | 4 | 6 | 6 | 2 | 4 | Yes |
| 2846102 | DYSF | B | 4 | 6 | 6 | 2 | 4 | Yes |
| 2855931 | DYSF | B | 4 | 5 | 6 | 1 | 6 | Yes |
| 2856173 | DYSF | B | 4 | 6 | 6 | 2 | 4 | Yes |
| 2866185 | DYSF | B | 5 | 5 | 6 | 2 | 6 | Yes |
| 2866240 | DYSF | B | 6 | 5 | 5 | 2 | 8 | Yes |
| 2885593 | DYSF | P | 5 | 3 | 3 | 2 | 11 | Yes |
| 2887146 | DYSF | B | 6 | 6 | 6 | 2 | 6 | Yes |
| 2893339 | DYSF | B | 5 | 6 | 6 | 2 | 5 | Yes |
| 2896555 | DYSF | B | 5 | 6 | 6 | 2 | 5 | Yes |
| 2900845 | DYSF | B | 7 | 6 | 6 | 2 | 7 | Yes |
| 2904027 | DYSF | B | 4 | 5 | 6 | 2 | 5 | Yes |
| 2909093 | DYSF | B | 5 | 5 | 6 | 2 | 6 | Yes |
| 2912301 | DYSF | B | 4 | 6 | 6 | 2 | 4 | Yes |
| 2912979 | DYSF | P | 4 | 3 | 5 | 2 | 8 | Yes |

| VariationID | Gene | Label | 101 bp wrong /12 | 201 bp correct /6 | 501 bp correct /6 | Native correct /2 | Incorrect predictions /26 | Any later correct |
|---|---|---|---|---|---|---|---|---|
| 2913813 | DYSF | B | 7 | 5 | 6 | 2 | 8 | Yes |
| 2915741 | DYSF | B | 4 | 6 | 6 | 2 | 4 | Yes |
| 2919542 | DYSF | P | 9 | 1 | 1 | 2 | 19 | Yes |
| 2956152 | DYSF | B | 4 | 6 | 6 | 2 | 4 | Yes |
| 2958148 | DYSF | B | 4 | 6 | 6 | 0 | 6 | Yes |
| 2960290 | DYSF | B | 7 | 4 | 5 | 1 | 11 | Yes |
| 2960420 | DYSF | B | 4 | 4 | 6 | 2 | 6 | Yes |
| 2977265 | DYSF | B | 5 | 6 | 5 | 2 | 6 | Yes |
| 2982101 | DYSF | B | 4 | 6 | 6 | 2 | 4 | Yes |
| 3016021 | DYSF | B | 5 | 6 | 6 | 2 | 5 | Yes |
| 3016502 | DYSF | B | 4 | 6 | 6 | 2 | 4 | Yes |
| 3018862 | DYSF | B | 5 | 5 | 5 | 2 | 7 | Yes |
| 3241379 | DYSF | P | 6 | 2 | 4 | 2 | 12 | Yes |
| 3586919 | DYSF | P | 5 | 3 | 1 | 2 | 13 | Yes |
| 3614195 | DYSF | B | 5 | 6 | 6 | 2 | 5 | Yes |
| 3620425 | DYSF | P | 5 | 1 | 2 | 2 | 14 | Yes |
| 3644742 | DYSF | B | 4 | 6 | 5 | 2 | 5 | Yes |
| 3646942 | DYSF | B | 6 | 5 | 6 | 2 | 7 | Yes |
| 3647048 | DYSF | B | 5 | 6 | 6 | 2 | 5 | Yes |
| 3677022 | DYSF | P | 7 | 0 | 1 | 2 | 18 | Yes |
| 3697856 | DYSF | B | 5 | 6 | 6 | 2 | 5 | Yes |
| 3705409 | DYSF | B | 7 | 5 | 6 | 2 | 8 | Yes |
| 3893281 | DYSF | P | 5 | 2 | 2 | 1 | 14 | Yes |
| 4078656 | DYSF | P | 6 | 1 | 3 | 2 | 14 | Yes |
| 4081362 | DYSF | P | 6 | 1 | 1 | 2 | 16 | Yes |
| 4705526 | DYSF | B | 4 | 5 | 5 | 2 | 6 | Yes |
| 4730505 | DYSF | B | 5 | 6 | 6 | 2 | 5 | Yes |
| 4746987 | DYSF | P | 7 | 1 | 1 | 2 | 17 | Yes |
| 4765017 | DYSF | B | 5 | 5 | 6 | 2 | 6 | Yes |
| 4815152 | DYSF | P | 7 | 0 | 1 | 2 | 18 | Yes |
| 4815154 | DYSF | P | 8 | 0 | 2 | 2 | 18 | Yes |
| 4820095 | DYSF | P | 4 | 3 | 2 | 2 | 11 | Yes |
| 4223 | FKRP | P | 11 | 0 | 0 | 2 | 23 | Yes |
| 120180 | FKRP | P | 8 | 1 | 1 | 2 | 18 | Yes |
| 241458 | FKRP | B | 5 | 6 | 6 | 2 | 5 | Yes |
| 553930 | FKRP | P | 5 | 1 | 1 | 2 | 15 | Yes |
| 556052 | FKRP | P | 7 | 3 | 3 | 2 | 13 | Yes |
| 681982 | FKRP | B | 5 | 6 | 6 | 2 | 5 | Yes |
| 766959 | FKRP | B | 4 | 6 | 6 | 2 | 4 | Yes |
| 788194 | FKRP | B | 5 | 6 | 6 | 2 | 5 | Yes |
| 1085022 | FKRP | B | 6 | 6 | 6 | 2 | 6 | Yes |
| 1100752 | FKRP | B | 5 | 6 | 6 | 2 | 5 | Yes |
| 1101867 | FKRP | B | 5 | 6 | 5 | 2 | 6 | Yes |
| 1103670 | FKRP | B | 9 | 4 | 6 | 2 | 11 | Yes |
| 1123016 | FKRP | B | 5 | 5 | 6 | 2 | 6 | Yes |
| 1133530 | FKRP | B | 7 | 5 | 6 | 2 | 8 | Yes |
| 1136570 | FKRP | B | 7 | 4 | 5 | 2 | 10 | Yes |
| 1139801 | FKRP | B | 6 | 5 | 5 | 2 | 8 | Yes |

| VariationID | Gene | Label | 101 bp wrong /12 | 201 bp correct /6 | 501 bp correct /6 | Native correct /2 | Incorrect predictions /26 | Any later correct |
|---|---|---|---|---|---|---|---|---|
| 1157937 | FKRP | B | 4 | 5 | 6 | 2 | 5 | Yes |
| 1180733 | FKRP | P | 6 | 0 | 1 | 2 | 17 | Yes |
| 1194483 | FKRP | B | 5 | 5 | 6 | 2 | 6 | Yes |
| 1270326 | FKRP | B | 5 | 6 | 6 | 2 | 5 | Yes |
| 1322919 | FKRP | P | 5 | 1 | 1 | 2 | 15 | Yes |
| 1334893 | FKRP | P | 12 | 0 | 1 | 2 | 23 | Yes |
| 1516962 | FKRP | P | 7 | 0 | 0 | 1 | 20 | Yes |
| 1524557 | FKRP | P | 10 | 0 | 1 | 2 | 21 | Yes |
| 1587048 | FKRP | B | 7 | 5 | 5 | 2 | 9 | Yes |
| 1636564 | FKRP | B | 5 | 4 | 5 | 2 | 8 | Yes |
| 1638745 | FKRP | B | 6 | 6 | 6 | 2 | 6 | Yes |
| 1639775 | FKRP | B | 5 | 5 | 5 | 2 | 7 | Yes |
| 1963124 | FKRP | P | 7 | 0 | 1 | 2 | 18 | Yes |
| 2040955 | FKRP | B | 4 | 6 | 6 | 2 | 4 | Yes |
| 2134240 | FKRP | P | 7 | 2 | 2 | 2 | 15 | Yes |
| 2148143 | FKRP | B | 6 | 5 | 6 | 2 | 7 | Yes |
| 2188258 | FKRP | B | 4 | 6 | 6 | 2 | 4 | Yes |
| 2577157 | FKRP | P | 8 | 0 | 1 | 2 | 19 | Yes |
| 2705709 | FKRP | B | 6 | 5 | 6 | 2 | 7 | Yes |
| 2736910 | FKRP | P | 10 | 0 | 1 | 2 | 21 | Yes |
| 2815755 | FKRP | P | 4 | 6 | 2 | 2 | 8 | Yes |
| 2879763 | FKRP | B | 4 | 6 | 5 | 2 | 5 | Yes |
| 2881456 | FKRP | B | 4 | 5 | 6 | 2 | 5 | Yes |
| 2909634 | FKRP | B | 4 | 5 | 6 | 2 | 5 | Yes |
| 3241624 | FKRP | P | 7 | 2 | 1 | 2 | 16 | Yes |
| 3241626 | FKRP | P | 6 | 2 | 2 | 2 | 14 | Yes |
| 3515559 | FKRP | B | 8 | 5 | 6 | 2 | 9 | Yes |
| 3515563 | FKRP | B | 6 | 6 | 6 | 2 | 6 | Yes |
| 3515564 | FKRP | B | 5 | 6 | 6 | 2 | 5 | Yes |
| 3515573 | FKRP | B | 4 | 6 | 6 | 2 | 4 | Yes |
| 3673199 | FKRP | B | 4 | 6 | 6 | 2 | 4 | Yes |
| 4746214 | FKRP | P | 12 | 0 | 1 | 2 | 23 | Yes |
| 14479 | LMNA | P | 7 | 3 | 2 | 2 | 14 | Yes |
| 14483 | LMNA | P | 6 | 0 | 0 | 2 | 18 | Yes |
| 14488 | LMNA | P | 9 | 3 | 1 | 2 | 17 | Yes |
| 14495 | LMNA | P | 5 | 3 | 2 | 2 | 12 | Yes |
| 14508 | LMNA | P | 8 | 3 | 1 | 2 | 16 | Yes |
| 14509 | LMNA | P | 8 | 0 | 0 | 2 | 20 | Yes |
| 14522 | LMNA | P | 8 | 0 | 0 | 1 | 21 | Yes |
| 14524 | LMNA | P | 5 | 4 | 3 | 2 | 10 | Yes |
| 29775 | LMNA | P | 6 | 2 | 1 | 2 | 15 | Yes |
| 48042 | LMNA | P | 9 | 0 | 1 | 2 | 20 | Yes |
| 48046 | LMNA | P | 7 | 1 | 2 | 2 | 16 | Yes |
| 48057 | LMNA | B | 4 | 6 | 6 | 2 | 4 | Yes |
| 48060 | LMNA | P | 9 | 1 | 2 | 2 | 18 | Yes |
| 48070 | LMNA | P | 6 | 1 | 0 | 2 | 17 | Yes |
| 48077 | LMNA | P | 6 | 5 | 4 | 2 | 9 | Yes |
| 48096 | LMNA | P | 5 | 4 | 4 | 2 | 9 | Yes |

| VariationID | Gene | Label | 101 bp wrong /12 | 201 bp correct /6 | 501 bp correct /6 | Native correct /2 | Incorrect predictions /26 | Any later correct |
|---|---|---|---|---|---|---|---|---|
| 66769 | LMNA | P | 7 | 1 | 3 | 2 | 15 | Yes |
| 66791 | LMNA | P | 6 | 2 | 2 | 2 | 14 | Yes |
| 66795 | LMNA | P | 8 | 1 | 1 | 2 | 18 | Yes |
| 66849 | LMNA | P | 8 | 1 | 1 | 2 | 18 | Yes |
| 66855 | LMNA | P | 11 | 0 | 0 | 1 | 24 | Yes |
| 66858 | LMNA | P | 7 | 1 | 1 | 2 | 17 | Yes |
| 66859 | LMNA | P | 6 | 4 | 2 | 2 | 12 | Yes |
| 66868 | LMNA | P | 5 | 2 | 2 | 2 | 13 | Yes |
| 66885 | LMNA | P | 9 | 1 | 0 | 2 | 20 | Yes |
| 66888 | LMNA | P | 10 | 0 | 1 | 2 | 21 | Yes |
| 66902 | LMNA | P | 4 | 2 | 1 | 1 | 14 | Yes |
| 66908 | LMNA | P | 5 | 2 | 2 | 2 | 13 | Yes |
| 66922 | LMNA | P | 6 | 2 | 1 | 2 | 15 | Yes |
| 66925 | LMNA | P | 8 | 1 | 1 | 2 | 18 | Yes |
| 66957 | LMNA | P | 8 | 2 | 1 | 1 | 18 | Yes |
| 183362 | LMNA | P | 10 | 1 | 1 | 2 | 20 | Yes |
| 200925 | LMNA | B | 5 | 6 | 6 | 2 | 5 | Yes |
| 200941 | LMNA | P | 7 | 0 | 0 | 1 | 20 | Yes |
| 217451 | LMNA | P | 5 | 1 | 1 | 2 | 15 | Yes |
| 245752 | LMNA | P | 7 | 1 | 1 | 2 | 17 | Yes |
| 246076 | LMNA | P | 7 | 2 | 1 | 2 | 16 | Yes |
| 285468 | LMNA | P | 6 | 2 | 2 | 2 | 14 | Yes |
| 292829 | LMNA | B | 6 | 4 | 6 | 2 | 8 | Yes |
| 408994 | LMNA | P | 6 | 2 | 2 | 2 | 14 | Yes |
| 415345 | LMNA | B | 5 | 6 | 6 | 2 | 5 | Yes |
| 415348 | LMNA | B | 4 | 6 | 6 | 2 | 4 | Yes |
| 429219 | LMNA | P | 4 | 2 | 1 | 2 | 13 | Yes |
| 435771 | LMNA | P | 7 | 0 | 2 | 2 | 17 | Yes |
| 444011 | LMNA | P | 10 | 0 | 1 | 2 | 21 | Yes |
| 476824 | LMNA | P | 8 | 2 | 1 | 2 | 17 | Yes |
| 476837 | LMNA | P | 7 | 1 | 2 | 2 | 16 | Yes |
| 516446 | LMNA | B | 5 | 6 | 6 | 2 | 5 | Yes |
| 519180 | LMNA | P | 5 | 2 | 3 | 1 | 13 | Yes |
| 520416 | LMNA | P | 7 | 1 | 0 | 2 | 18 | Yes |
| 543191 | LMNA | P | 7 | 2 | 1 | 2 | 16 | Yes |
| 543244 | LMNA | P | 7 | 3 | 3 | 2 | 13 | Yes |
| 628758 | LMNA | B | 8 | 4 | 6 | 2 | 10 | Yes |
| 634638 | LMNA | P | 7 | 2 | 3 | 2 | 14 | Yes |
| 636306 | LMNA | P | 4 | 4 | 3 | 2 | 9 | Yes |
| 649203 | LMNA | P | 4 | 2 | 1 | 2 | 13 | Yes |
| 652066 | LMNA | P | 9 | 1 | 2 | 2 | 18 | Yes |
| 663822 | LMNA | P | 4 | 3 | 1 | 2 | 12 | Yes |
| 668922 | LMNA | B | 7 | 6 | 6 | 2 | 7 | Yes |
| 672779 | LMNA | B | 4 | 6 | 6 | 2 | 4 | Yes |
| 683622 | LMNA | B | 5 | 5 | 5 | 2 | 7 | Yes |
| 745112 | LMNA | B | 4 | 6 | 6 | 2 | 4 | Yes |
| 846468 | LMNA | P | 8 | 1 | 1 | 2 | 18 | Yes |
| 916767 | LMNA | B | 7 | 3 | 4 | 0 | 14 | Yes |

| VariationID | Gene | Label | 101 bp wrong /12 | 201 bp correct /6 | 501 bp correct /6 | Native correct /2 | Incorrect predictions /26 | Any later correct |
|---|---|---|---|---|---|---|---|---|
| 920428 | LMNA | B | 6 | 5 | 6 | 2 | 7 | Yes |
| 921773 | LMNA | B | 4 | 6 | 6 | 2 | 4 | Yes |
| 935833 | LMNA | P | 6 | 1 | 2 | 2 | 15 | Yes |
| 943114 | LMNA | P | 9 | 0 | 1 | 2 | 20 | Yes |
| 949101 | LMNA | P | 8 | 2 | 1 | 2 | 17 | Yes |
| 994013 | LMNA | B | 6 | 6 | 6 | 2 | 6 | Yes |
| 1063701 | LMNA | P | 8 | 1 | 1 | 1 | 19 | Yes |
| 1067826 | LMNA | P | 8 | 0 | 1 | 2 | 19 | Yes |
| 1103137 | LMNA | B | 7 | 6 | 6 | 2 | 7 | Yes |
| 1284262 | LMNA | B | 6 | 6 | 6 | 2 | 6 | Yes |
| 1299837 | LMNA | P | 7 | 1 | 2 | 2 | 16 | Yes |
| 1308669 | LMNA | P | 5 | 3 | 1 | 2 | 13 | Yes |
| 1341358 | LMNA | P | 6 | 2 | 2 | 1 | 15 | Yes |
| 1434909 | LMNA | P | 5 | 4 | 4 | 2 | 9 | Yes |
| 1457783 | LMNA | P | 7 | 4 | 3 | 2 | 12 | Yes |
| 1467968 | LMNA | P | 4 | 1 | 2 | 2 | 13 | Yes |
| 1575868 | LMNA | B | 4 | 5 | 5 | 2 | 6 | Yes |
| 1577186 | LMNA | B | 5 | 6 | 6 | 2 | 5 | Yes |
| 1613218 | LMNA | B | 5 | 4 | 6 | 2 | 7 | Yes |
| 1658443 | LMNA | B | 5 | 6 | 5 | 2 | 6 | Yes |
| 1685368 | LMNA | P | 9 | 1 | 0 | 2 | 20 | Yes |
| 1694530 | LMNA | B | 6 | 6 | 6 | 2 | 6 | Yes |
| 1749770 | LMNA | B | 9 | 3 | 4 | 2 | 14 | Yes |
| 1769429 | LMNA | B | 5 | 5 | 6 | 2 | 6 | Yes |
| 1960601 | LMNA | B | 6 | 6 | 4 | 2 | 8 | Yes |
| 1966731 | LMNA | B | 4 | 6 | 5 | 2 | 5 | Yes |
| 2029375 | LMNA | B | 4 | 6 | 6 | 2 | 4 | Yes |
| 2032046 | LMNA | B | 6 | 5 | 4 | 2 | 9 | Yes |
| 2065107 | LMNA | P | 6 | 0 | 0 | 2 | 18 | Yes |
| 2089379 | LMNA | P | 5 | 3 | 3 | 2 | 11 | Yes |
| 2151903 | LMNA | P | 6 | 2 | 1 | 2 | 15 | Yes |
| 2202857 | LMNA | P | 6 | 2 | 2 | 2 | 14 | Yes |
| 2452985 | LMNA | P | 6 | 1 | 3 | 2 | 14 | Yes |
| 2503449 | LMNA | P | 8 | 1 | 1 | 2 | 18 | Yes |
| 2572315 | LMNA | P | 10 | 3 | 3 | 2 | 16 | Yes |
| 2709822 | LMNA | P | 7 | 1 | 1 | 2 | 17 | Yes |
| 2733992 | LMNA | P | 6 | 1 | 2 | 2 | 15 | Yes |
| 2798165 | LMNA | B | 4 | 5 | 6 | 2 | 5 | Yes |
| 2882128 | LMNA | P | 4 | 4 | 3 | 2 | 9 | Yes |
| 2965632 | LMNA | B | 6 | 4 | 6 | 2 | 8 | Yes |
| 2966143 | LMNA | B | 5 | 6 | 5 | 2 | 6 | Yes |
| 3009340 | LMNA | B | 4 | 6 | 5 | 2 | 5 | Yes |
| 3070531 | LMNA | B | 4 | 6 | 6 | 2 | 4 | Yes |
| 3229275 | LMNA | P | 8 | 2 | 1 | 2 | 17 | Yes |
| 3236851 | LMNA | P | 4 | 4 | 3 | 2 | 9 | Yes |
| 3349147 | LMNA | B | 6 | 6 | 6 | 2 | 6 | Yes |
| 3538726 | LMNA | B | 5 | 5 | 4 | 2 | 8 | Yes |
| 3538733 | LMNA | B | 4 | 6 | 6 | 2 | 4 | Yes |

| VariationID | Gene | Label | 101 bp wrong /12 | 201 bp correct /6 | 501 bp correct /6 | Native correct /2 | Incorrect predictions /26 | Any later correct |
|---|---|---|---|---|---|---|---|---|
| 3629832 | LMNA | P | 6 | 1 | 2 | 2 | 15 | Yes |
| 3633621 | LMNA | P | 8 | 1 | 1 | 1 | 19 | Yes |
| 3633629 | LMNA | P | 9 | 1 | 2 | 2 | 18 | Yes |
| 3647177 | LMNA | B | 4 | 5 | 6 | 2 | 5 | Yes |
| 3667532 | LMNA | B | 4 | 6 | 6 | 2 | 4 | Yes |
| 3677001 | LMNA | P | 12 | 1 | 0 | 2 | 23 | Yes |
| 3705844 | LMNA | B | 4 | 6 | 6 | 2 | 4 | Yes |
| 3775011 | LMNA | P | 5 | 2 | 2 | 2 | 13 | Yes |
| 3893920 | LMNA | B | 7 | 6 | 6 | 2 | 7 | Yes |
| 3895533 | LMNA | P | 7 | 1 | 0 | 1 | 19 | Yes |
| 3907369 | LMNA | P | 5 | 2 | 3 | 2 | 12 | Yes |
| 4098591 | LMNA | B | 6 | 3 | 4 | 2 | 11 | Yes |
| 4716669 | LMNA | B | 5 | 6 | 5 | 2 | 6 | Yes |
| 4726441 | LMNA | P | 7 | 2 | 2 | 2 | 15 | Yes |
| 4732685 | LMNA | P | 7 | 2 | 2 | 2 | 15 | Yes |
| 4799953 | LMNA | P | 7 | 0 | 1 | 2 | 18 | Yes |
| 4819325 | LMNA | P | 9 | 2 | 0 | 2 | 19 | Yes |
| 188733 | SGCA | P | 9 | 0 | 0 | 0 | 23 | No |
| 197094 | SGCA | P | 9 | 0 | 0 | 0 | 23 | No |
| 217250 | SGCA | P | 5 | 1 | 1 | 1 | 16 | Yes |
| 284708 | SGCA | P | 8 | 1 | 0 | 0 | 21 | Yes |
| 286025 | SGCA | P | 8 | 0 | 1 | 2 | 19 | Yes |
| 286049 | SGCA | P | 10 | 0 | 0 | 2 | 22 | Yes |
| 432007 | SGCA | P | 10 | 0 | 1 | 2 | 21 | Yes |
| 449294 | SGCA | P | 7 | 2 | 2 | 2 | 15 | Yes |
| 471337 | SGCA | P | 9 | 2 | 0 | 1 | 20 | Yes |
| 496854 | SGCA | B | 5 | 6 | 6 | 2 | 5 | Yes |
| 497672 | SGCA | P | 9 | 1 | 1 | 2 | 19 | Yes |
| 498275 | SGCA | P | 9 | 0 | 0 | 0 | 23 | No |
| 538664 | SGCA | B | 7 | 6 | 6 | 2 | 7 | Yes |
| 581745 | SGCA | P | 9 | 1 | 1 | 1 | 20 | Yes |
| 617535 | SGCA | P | 6 | 1 | 1 | 0 | 18 | Yes |
| 659203 | SGCA | P | 10 | 0 | 0 | 2 | 22 | Yes |
| 669838 | SGCA | B | 7 | 6 | 5 | 2 | 8 | Yes |
| 755603 | SGCA | B | 5 | 6 | 6 | 2 | 5 | Yes |
| 857216 | SGCA | P | 7 | 2 | 2 | 2 | 15 | Yes |
| 988362 | SGCA | P | 5 | 3 | 2 | 2 | 12 | Yes |
| 1066710 | SGCA | P | 7 | 1 | 2 | 2 | 16 | Yes |
| 1066846 | SGCA | P | 5 | 2 | 3 | 2 | 12 | Yes |
| 1108908 | SGCA | B | 4 | 6 | 6 | 2 | 4 | Yes |
| 1347617 | SGCA | B | 4 | 5 | 6 | 2 | 5 | Yes |
| 1460416 | SGCA | B | 7 | 6 | 5 | 2 | 8 | Yes |
| 1492140 | SGCA | P | 10 | 0 | 0 | 2 | 22 | Yes |
| 1498501 | SGCA | P | 10 | 1 | 0 | 2 | 21 | Yes |
| 1596584 | SGCA | B | 5 | 6 | 5 | 2 | 6 | Yes |
| 1596960 | SGCA | B | 6 | 6 | 6 | 2 | 6 | Yes |
| 1597922 | SGCA | B | 5 | 5 | 6 | 2 | 6 | Yes |
| 1638217 | SGCA | B | 7 | 6 | 6 | 2 | 7 | Yes |

| VariationID | Gene | Label | 101 bp wrong /12 | 201 bp correct /6 | 501 bp correct /6 | Native correct /2 | Incorrect predictions /26 | Any later correct |
|---|---|---|---|---|---|---|---|---|
| 1705242 | SGCA | P | 8 | 0 | 1 | 1 | 20 | Yes |
| 1967847 | SGCA | P | 9 | 0 | 1 | 2 | 20 | Yes |
| 1983230 | SGCA | B | 4 | 6 | 6 | 2 | 4 | Yes |
| 2141344 | SGCA | P | 10 | 1 | 1 | 1 | 21 | Yes |
| 2585416 | SGCA | P | 7 | 1 | 1 | 2 | 17 | Yes |
| 2690696 | SGCA | P | 6 | 4 | 2 | 2 | 12 | Yes |
| 2697305 | SGCA | B | 4 | 6 | 5 | 2 | 5 | Yes |
| 2707194 | SGCA | P | 8 | 0 | 1 | 2 | 19 | Yes |
| 2736615 | SGCA | P | 6 | 1 | 1 | 2 | 16 | Yes |
| 2740495 | SGCA | B | 4 | 6 | 6 | 2 | 4 | Yes |
| 2741637 | SGCA | B | 4 | 6 | 6 | 2 | 4 | Yes |
| 2779011 | SGCA | B | 6 | 5 | 6 | 2 | 7 | Yes |
| 2832263 | SGCA | B | 6 | 6 | 6 | 2 | 6 | Yes |
| 2920016 | SGCA | B | 4 | 6 | 6 | 2 | 4 | Yes |
| 2980368 | SGCA | B | 5 | 6 | 6 | 2 | 5 | Yes |
| 2997307 | SGCA | B | 4 | 5 | 5 | 2 | 6 | Yes |
| 3240442 | SGCA | P | 5 | 2 | 4 | 2 | 11 | Yes |
| 3240445 | SGCA | P | 5 | 3 | 3 | 2 | 11 | Yes |
| 3384982 | SGCA | P | 9 | 0 | 0 | 0 | 23 | No |
| 4077502 | SGCA | P | 6 | 1 | 3 | 1 | 15 | Yes |
| 4293792 | SGCA | P | 9 | 0 | 1 | 2 | 20 | Yes |
| 4847205 | SGCA | P | 6 | 3 | 4 | 2 | 11 | Yes |
| 8717 | SGCB | P | 9 | 1 | 1 | 2 | 19 | Yes |
| 42035 | SGCB | P | 7 | 2 | 1 | 2 | 16 | Yes |
| 284502 | SGCB | P | 10 | 2 | 0 | 1 | 21 | Yes |
| 534948 | SGCB | P | 9 | 1 | 1 | 2 | 19 | Yes |
| 551412 | SGCB | B | 6 | 6 | 6 | 2 | 6 | Yes |
| 551805 | SGCB | P | 7 | 0 | 0 | 2 | 19 | Yes |
| 553619 | SGCB | P | 6 | 2 | 1 | 2 | 15 | Yes |
| 554369 | SGCB | P | 6 | 1 | 2 | 2 | 15 | Yes |
| 555821 | SGCB | P | 9 | 1 | 1 | 2 | 19 | Yes |
| 558332 | SGCB | P | 5 | 2 | 1 | 2 | 14 | Yes |
| 665890 | SGCB | P | 9 | 1 | 1 | 2 | 19 | Yes |
| 802072 | SGCB | P | 9 | 0 | 1 | 2 | 20 | Yes |
| 862654 | SGCB | P | 8 | 0 | 0 | 1 | 21 | Yes |
| 936115 | SGCB | P | 7 | 1 | 3 | 2 | 15 | Yes |
| 1076257 | SGCB | P | 6 | 1 | 1 | 2 | 16 | Yes |
| 1112695 | SGCB | B | 5 | 6 | 6 | 2 | 5 | Yes |
| 1187958 | SGCB | B | 4 | 6 | 6 | 2 | 4 | Yes |
| 1479714 | SGCB | P | 10 | 0 | 0 | 2 | 22 | Yes |
| 1580523 | SGCB | B | 5 | 5 | 5 | 2 | 7 | Yes |
| 1630975 | SGCB | B | 6 | 6 | 6 | 2 | 6 | Yes |
| 1640962 | SGCB | B | 5 | 6 | 6 | 2 | 5 | Yes |
| 1659361 | SGCB | B | 5 | 6 | 6 | 2 | 5 | Yes |
| 1686178 | SGCB | P | 5 | 2 | 2 | 2 | 13 | Yes |
| 1686638 | SGCB | P | 11 | 0 | 0 | 1 | 24 | Yes |
| 2030054 | SGCB | B | 8 | 4 | 6 | 2 | 10 | Yes |
| 2036333 | SGCB | B | 5 | 6 | 6 | 2 | 5 | Yes |

| VariationID | Gene | Label | 101 bp wrong /12 | 201 bp correct /6 | 501 bp correct /6 | Native correct /2 | Incorrect predictions /26 | Any later correct |
|---|---|---|---|---|---|---|---|---|
| 2065460 | SGCB | B | 4 | 6 | 6 | 2 | 4 | Yes |
| 2789475 | SGCB | B | 5 | 6 | 6 | 2 | 5 | Yes |
| 2828840 | SGCB | B | 7 | 6 | 6 | 2 | 7 | Yes |
| 2839791 | SGCB | B | 5 | 6 | 5 | 2 | 6 | Yes |
| 2839873 | SGCB | B | 5 | 5 | 6 | 2 | 6 | Yes |
| 2855036 | SGCB | B | 4 | 5 | 5 | 2 | 6 | Yes |
| 2967226 | SGCB | B | 6 | 6 | 6 | 2 | 6 | Yes |
| 3590583 | SGCB | P | 8 | 1 | 2 | 2 | 17 | Yes |
| 4277883 | SGCB | P | 8 | 1 | 0 | 1 | 20 | Yes |
| 4293121 | SGCB | P | 10 | 0 | 0 | 1 | 23 | Yes |
| 4718937 | SGCB | B | 8 | 4 | 5 | 2 | 11 | Yes |
| 4734039 | SGCB | B | 5 | 6 | 6 | 2 | 5 | Yes |
| 2009 | SGCG | P | 8 | 1 | 0 | 2 | 19 | Yes |
| 130295 | SGCG | B | 6 | 6 | 6 | 2 | 6 | Yes |
| 167681 | SGCG | B | 4 | 6 | 6 | 2 | 4 | Yes |
| 281085 | SGCG | P | 10 | 1 | 1 | 2 | 20 | Yes |
| 311478 | SGCG | B | 4 | 6 | 5 | 2 | 5 | Yes |
| 670551 | SGCG | B | 6 | 5 | 6 | 2 | 7 | Yes |
| 684356 | SGCG | B | 7 | 6 | 6 | 2 | 7 | Yes |
| 705034 | SGCG | B | 6 | 6 | 6 | 2 | 6 | Yes |
| 805445 | SGCG | B | 8 | 4 | 6 | 2 | 10 | Yes |
| 813974 | SGCG | P | 5 | 2 | 2 | 2 | 13 | Yes |
| 1099752 | SGCG | B | 5 | 6 | 6 | 2 | 5 | Yes |
| 1105694 | SGCG | B | 5 | 5 | 6 | 2 | 6 | Yes |
| 1177521 | SGCG | B | 4 | 6 | 6 | 2 | 4 | Yes |
| 1209395 | SGCG | B | 6 | 5 | 6 | 2 | 7 | Yes |
| 1213219 | SGCG | B | 5 | 5 | 6 | 2 | 6 | Yes |
| 1459762 | SGCG | P | 5 | 1 | 2 | 2 | 14 | Yes |
| 1598164 | SGCG | B | 4 | 6 | 6 | 2 | 4 | Yes |
| 1650782 | SGCG | B | 4 | 5 | 6 | 2 | 5 | Yes |
| 1691313 | SGCG | P | 8 | 1 | 1 | 2 | 18 | Yes |
| 1725964 | SGCG | P | 7 | 1 | 2 | 2 | 16 | Yes |
| 2119723 | SGCG | B | 5 | 6 | 6 | 2 | 5 | Yes |
| 2793307 | SGCG | B | 4 | 6 | 6 | 2 | 4 | Yes |
| 2890939 | SGCG | B | 4 | 6 | 6 | 2 | 4 | Yes |
| 2919971 | SGCG | B | 6 | 6 | 6 | 2 | 6 | Yes |
| 2963224 | SGCG | B | 7 | 6 | 6 | 2 | 7 | Yes |
| 2985916 | SGCG | B | 4 | 6 | 5 | 2 | 5 | Yes |
| 3005045 | SGCG | B | 5 | 5 | 6 | 2 | 6 | Yes |
| 3575784 | SGCG | P | 6 | 3 | 2 | 2 | 13 | Yes |
| 3644861 | SGCG | B | 5 | 5 | 6 | 2 | 6 | Yes |
| 4077506 | SGCG | P | 6 | 1 | 2 | 2 | 15 | Yes |
| 4810277 | SGCG | B | 5 | 5 | 6 | 2 | 6 | Yes |
| 44709 | TCAP | B | 7 | 4 | 5 | 2 | 10 | Yes |
| 202113 | TCAP | P | 5 | 4 | 4 | 2 | 9 | Yes |
| 286261 | TCAP | P | 5 | 1 | 1 | 2 | 15 | Yes |
| 1055526 | TCAP | P | 6 | 3 | 4 | 2 | 11 | Yes |
| 1066432 | TCAP | P | 10 | 1 | 2 | 2 | 19 | Yes |

| VariationID | Gene | Label | 101 bp wrong /12 | 201 bp correct /6 | 501 bp correct /6 | Native correct /2 | Incorrect predictions /26 | Any later correct |
|---|---|---|---|---|---|---|---|---|
| 1732581 | TCAP | B | 5 | 6 | 6 | 2 | 5 | Yes |
| 3453999 | TCAP | B | 5 | 5 | 5 | 2 | 7 | Yes |
| 3454000 | TCAP | B | 7 | 4 | 6 | 2 | 9 | Yes |

#### Supplementary Table S7. Strategy-level classification error burden

| Strategy | Subset | Variants | Configs | Mean wrong per variant | Median wrong | Zero wrong | All six wrong | Variant-config accuracy |
|---|---|---|---|---|---|---|---|---|
| Frozen 101 bp | All variants | 2094 | 6 | 1.342 | 0.0 | 1086 | 96 | 77.6% |
| Frozen 101 bp | Negative class | 1585 | 6 | 0.391 | 0.0 | 1086 | 0 | 93.5% |
| Frozen 101 bp | Positive class | 509 | 6 | 4.305 | 5.0 | 0 | 96 | 28.3% |
| Adapted 101 bp | All variants | 2094 | 6 | 2.294 | 2.0 | 249 | 46 | 61.8% |
| Adapted 101 bp | Negative class | 1585 | 6 | 2.471 | 2.0 | 149 | 41 | 58.8% |
| Adapted 101 bp | Positive class | 509 | 6 | 1.743 | 1.0 | 100 | 5 | 71.0% |
| Frozen 201 bp | All variants | 2094 | 6 | 1.241 | 0.0 | 1198 | 80 | 79.3% |
| Frozen 201 bp | Negative class | 1585 | 6 | 0.291 | 0.0 | 1195 | 0 | 95.2% |
| Frozen 201 bp | Positive class | 509 | 6 | 4.198 | 4.0 | 3 | 80 | 30.0% |
| Frozen 501 bp | All variants | 2094 | 6 | 1.145 | 0.0 | 1364 | 59 | 80.9% |
| Frozen 501 bp | Negative class | 1585 | 6 | 0.159 | 0.0 | 1362 | 0 | 97.4% |
| Frozen 501 bp | Positive class | 509 | 6 | 4.214 | 5.0 | 2 | 59 | 29.8% |

Each strategy contains six sequence-model configurations. “All six wrong” counts variants misclassified by all six configurations within the specified strategy. Variant-configuration accuracy was calculated as 1 − mean errors/6 and represents the proportion of correct model-by-variant predictions across the six configurations. It does not measure the performance of a separately trained ensemble.

#### Supplementary Table S8. Native-score classification by sequence-model difficulty

| Sequence difficulty (wrong of 24) | Variants | Mean seq. errors | AVI correct | GPN-V100 correct | Either native correct | Either percent | All 26 wrong |
|---|---|---|---|---|---|---|---|
| 0-5 wrong | 1352 | 2.62 | 1347 | 1341 | 1348 | 99.7% | 0 |
| 6-11 wrong | 355 | 7.72 | 343 | 344 | 351 | 98.9% | 0 |
| 12-17 wrong | 254 | 14.59 | 242 | 242 | 252 | 99.2% | 0 |
| 18-23 wrong | 131 | 19.57 | 114 | 112 | 124 | 94.7% | 0 |
| 24 wrong | 2 | 24.00 | 1 | 1 | 1 | 50.0% | 1 |

Difficulty strata were defined by the number of incorrect predictions across all 24 sequence configurations. Native-score correctness was assessed retrospectively against the known benchmark labels, using the existing classification thresholds. “Either native correct” counts variants correctly classified by GPN-Star V100, AlphaGenome AVI or both. The final stratum contains only two variants, making its observed 50% coverage unsuitable for generalization. Across all strata, 2,047 variants were correctly classified by AVI, 2,040 by GPN-Star V100 and 2,076 by at least one native score. One variant remained misclassified across all 26 configurations.

#### Supplementary Table S9. Residual variant misclassified by all 26 primary configurations

| ClinVar ID | Gene | GRCh38 locus | REF>ALT | Label | 101bp wrong /12 | 201/501 wrong /12 | Native wrong /2 | All wrong /26 |
|---|---|---|---|---|---|---|---|---|
| 4538462 | ANO5 | chr11:22239682 | T>G | Positive | 12 | 12 | 2 | 26 |

For this variant, AlphaGenome Atlas AVI yielded a PHRED score of 23.670678, while GPN-Star V100 yielded a sign-reversed mutation-rate-calibrated log-likelihood ratio of 5.001000. Both scores fell below their respective training-derived classification thresholds, producing negative predictions despite the positive ClinVar benchmark label. Additional clinical or functional evidence would be required to assess the basis of this disagreement.

**Source and interpretation note.** Tables S5, S6 and S9 were verified against archived variant-level integrated records. The identity, predicted classification and AVI PHRED score of the residual universally misclassified variant were also checked against the original AVI test predictions. Tables S7 and S8 were reconciled with the complete 2,094-variant test partition and checked for internal arithmetic consistency. Their individual strata were not independently reconstructed from all 26 original configuration-level prediction streams during supplementary assembly. The reported coverage describes retrospective classification outcomes using known benchmark labels; prospective ensemble performance was not evaluated.

#### Supplementary Table S10. Gene-level results under two distinct evaluation protocols

Panel A reports 60 gene-held-out evaluations of frozen 101-bp sequence readouts, with each downstream classifier trained using labelled variants from the remaining nine genes. Panel B reports 20 native-score evaluations using unchanged continuous scores and gene-excluded threshold selection. The two protocols differ in their downstream training procedures and should be interpreted separately. DNAJB6, SGCG and TCAP contain only 2, 11 and 6 positive-class test variants, respectively. Gene-level AUROC and AUPRC values cannot be averaged to reconstruct the pooled estimates reported in main Table 4.

#### Panel A. Frozen embedding readouts (60 gene-model evaluations)

| Model | Excluded gene | Test N | Positive N | MCC | AUROC | AUPRC |
|---|---|---|---|---|---|---|
| DNABERT-2 | ANO5 | 164 | 33 | 0.119 | 0.482 | 0.205 |
| DNABERT-2 | CAPN3 | 295 | 88 | 0.068 | 0.524 | 0.338 |
| DNABERT-2 | DNAJB6 | 65 | 2 | 0.187 | 0.611 | 0.094 |
| DNABERT-2 | DYSF | 768 | 166 | 0.084 | 0.580 | 0.264 |
| DNABERT-2 | FKRP | 172 | 26 | 0.007 | 0.548 | 0.168 |
| DNABERT-2 | LMNA | 284 | 117 | 0.168 | 0.589 | 0.504 |
| DNABERT-2 | SGCA | 144 | 40 | -0.104 | 0.459 | 0.307 |
| DNABERT-2 | SGCB | 83 | 20 | -0.088 | 0.381 | 0.201 |
| DNABERT-2 | SGCG | 84 | 11 | 0.224 | 0.639 | 0.334 |
| DNABERT-2 | TCAP | 35 | 6 | 0.195 | 0.575 | 0.493 |
| NT-v2 | ANO5 | 164 | 33 | 0.172 | 0.576 | 0.342 |
| NT-v2 | CAPN3 | 295 | 88 | 0.196 | 0.669 | 0.487 |
| NT-v2 | DNAJB6 | 65 | 2 | -0.039 | 0.849 | 0.134 |
| NT-v2 | DYSF | 768 | 166 | 0.229 | 0.675 | 0.409 |
| NT-v2 | FKRP | 172 | 26 | 0.158 | 0.698 | 0.336 |
| NT-v2 | LMNA | 284 | 117 | 0.232 | 0.572 | 0.540 |
| NT-v2 | SGCA | 144 | 40 | 0.075 | 0.674 | 0.426 |
| NT-v2 | SGCB | 83 | 20 | 0.032 | 0.489 | 0.284 |
| NT-v2 | SGCG | 84 | 11 | 0.094 | 0.621 | 0.215 |
| NT-v2 | TCAP | 35 | 6 | 0.551 | 0.874 | 0.661 |
| HyenaDNA | ANO5 | 164 | 33 | -0.001 | 0.539 | 0.241 |
| HyenaDNA | CAPN3 | 295 | 88 | 0.101 | 0.642 | 0.414 |
| HyenaDNA | DNAJB6 | 65 | 2 | 0.484 | 0.921 | 0.583 |
| HyenaDNA | DYSF | 768 | 166 | 0.185 | 0.639 | 0.349 |
| HyenaDNA | FKRP | 172 | 26 | 0.263 | 0.640 | 0.300 |
| HyenaDNA | LMNA | 284 | 117 | 0.133 | 0.638 | 0.548 |
| HyenaDNA | SGCA | 144 | 40 | 0.224 | 0.666 | 0.532 |
| HyenaDNA | SGCB | 83 | 20 | 0.095 | 0.549 | 0.340 |
| HyenaDNA | SGCG | 84 | 11 | 0.029 | 0.616 | 0.208 |
| HyenaDNA | TCAP | 35 | 6 | 0.402 | 0.753 | 0.472 |
| Caduceus-Ph | ANO5 | 164 | 33 | 0.019 | 0.570 | 0.239 |
| Caduceus-Ph | CAPN3 | 295 | 88 | 0.113 | 0.603 | 0.404 |
| Caduceus-Ph | DNAJB6 | 65 | 2 | -0.046 | 0.492 | 0.044 |
| Caduceus-Ph | DYSF | 768 | 166 | 0.227 | 0.639 | 0.388 |
| Caduceus-Ph | FKRP | 172 | 26 | 0.078 | 0.620 | 0.271 |
| Caduceus-Ph | LMNA | 284 | 117 | 0.224 | 0.614 | 0.590 |
| Caduceus-Ph | SGCA | 144 | 40 | 0.178 | 0.567 | 0.414 |
| Caduceus-Ph | SGCB | 83 | 20 | -0.157 | 0.630 | 0.308 |

| Model | Excluded gene | Test N | Positive N | MCC | AUROC | AUPRC |
|---|---|---|---|---|---|---|
| Caduceus-Ph | SGCG | 84 | 11 | 0.094 | 0.638 | 0.222 |
| Caduceus-Ph | TCAP | 35 | 6 | -0.163 | 0.776 | 0.331 |
| modernGENA | ANO5 | 164 | 33 | 0.169 | 0.499 | 0.290 |
| modernGENA | CAPN3 | 295 | 88 | 0.187 | 0.612 | 0.439 |
| modernGENA | DNAJB6 | 65 | 2 | 0.225 | 0.754 | 0.158 |
| modernGENA | DYSF | 768 | 166 | 0.214 | 0.655 | 0.357 |
| modernGENA | FKRP | 172 | 26 | 0.034 | 0.642 | 0.264 |
| modernGENA | LMNA | 284 | 117 | 0.227 | 0.608 | 0.568 |
| modernGENA | SGCA | 144 | 40 | 0.129 | 0.629 | 0.398 |
| modernGENA | SGCB | 83 | 20 | -0.089 | 0.514 | 0.302 |
| modernGENA | SGCG | 84 | 11 | 0.115 | 0.513 | 0.216 |
| modernGENA | TCAP | 35 | 6 | 0.541 | 0.586 | 0.477 |
| Evo2 | ANO5 | 164 | 33 | 0.450 | 0.765 | 0.643 |
| Evo2 | CAPN3 | 295 | 88 | 0.676 | 0.870 | 0.854 |
| Evo2 | DNAJB6 | 65 | 2 | 0.702 | 1.000 | 1.000 |
| Evo2 | DYSF | 768 | 166 | 0.700 | 0.899 | 0.825 |
| Evo2 | FKRP | 172 | 26 | 0.619 | 0.711 | 0.637 |
| Evo2 | LMNA | 284 | 117 | 0.647 | 0.888 | 0.892 |
| Evo2 | SGCA | 144 | 40 | 0.521 | 0.792 | 0.728 |
| Evo2 | SGCB | 83 | 20 | 0.389 | 0.639 | 0.555 |
| Evo2 | SGCG | 84 | 11 | 0.648 | 0.956 | 0.867 |
| Evo2 | TCAP | 35 | 6 | 0.541 | 0.730 | 0.586 |

#### Panel B. Native scoring with nine-gene threshold selection (20 gene-model evaluations)

| Model | Excluded gene | Test N | Positive N | MCC | AUROC | AUPRC |
|---|---|---|---|---|---|---|
| GPN-Star V100 | ANO5 | 164 | 33 | 0.903 | 0.986 | 0.977 |
| GPN-Star V100 | CAPN3 | 295 | 88 | 0.951 | 0.995 | 0.993 |
| GPN-Star V100 | DNAJB6 | 65 | 2 | 1.000 | 1.000 | 1.000 |
| GPN-Star V100 | DYSF | 768 | 166 | 0.921 | 0.993 | 0.984 |
| GPN-Star V100 | FKRP | 172 | 26 | 1.000 | 1.000 | 1.000 |
| GPN-Star V100 | LMNA | 284 | 117 | 0.942 | 0.989 | 0.990 |
| GPN-Star V100 | SGCA | 144 | 40 | 0.842 | 0.994 | 0.985 |
| GPN-Star V100 | SGCB | 83 | 20 | 0.834 | 0.961 | 0.956 |
| GPN-Star V100 | SGCG | 84 | 11 | 1.000 | 1.000 | 1.000 |
| GPN-Star V100 | TCAP | 35 | 6 | 0.898 | 0.989 | 0.958 |
| AlphaGenome AVI | ANO5 | 164 | 33 | 0.962 | 1.000 | 1.000 |
| AlphaGenome AVI | CAPN3 | 295 | 88 | 0.853 | 0.956 | 0.957 |
| AlphaGenome AVI | DNAJB6 | 65 | 2 | 1.000 | 1.000 | 1.000 |
| AlphaGenome AVI | DYSF | 768 | 166 | 0.958 | 0.999 | 0.996 |
| AlphaGenome AVI | FKRP | 172 | 26 | 0.977 | 1.000 | 1.000 |
| AlphaGenome AVI | LMNA | 284 | 117 | 0.964 | 0.997 | 0.996 |
| AlphaGenome AVI | SGCA | 144 | 40 | 0.827 | 0.980 | 0.968 |
| AlphaGenome AVI | SGCB | 83 | 20 | 1.000 | 1.000 | 1.000 |
| AlphaGenome AVI | SGCG | 84 | 11 | 1.000 | 1.000 | 1.000 |
| AlphaGenome AVI | TCAP | 35 | 6 | 1.000 | 1.000 | 1.000 |

#### Supplementary Table S11. Exploratory paired locus-bootstrap differences

Paired differences were calculated as Evo2 minus the indicated comparator, with positive values favoring Evo2. The 95% percentile confidence intervals were obtained from 1,000 paired genomic-locus bootstrap replicates within evaluation genes. These exploratory intervals are conditional on the observed genes and fitted predictors. No multiple-testing adjustment was applied.

#### Panel A. Frozen Evo2 versus the five other frozen 101-bp readouts

| Comparator | Metric | Evo2 – comparator | 95% bootstrap CI | Excludes zero |
|---|---|---|---|---|
| DNABERT-2 | MCC | +0.563 | [+0.499, +0.628] | Yes |
| DNABERT-2 | AUROC | +0.312 | [+0.272, +0.351] | Yes |
| DNABERT-2 | AUPRC | +0.509 | [+0.465, +0.552] | Yes |
| NT-v2 | MCC | +0.450 | [+0.388, +0.516] | Yes |
| NT-v2 | AUROC | +0.218 | [+0.180, +0.257] | Yes |
| NT-v2 | AUPRC | +0.394 | [+0.344, +0.443] | Yes |
| HyenaDNA | MCC | +0.487 | [+0.421, +0.553] | Yes |
| HyenaDNA | AUROC | +0.231 | [+0.196, +0.271] | Yes |
| HyenaDNA | AUPRC | +0.435 | [+0.385, +0.483] | Yes |
| Caduceus-Ph | MCC | +0.480 | [+0.417, +0.545] | Yes |
| Caduceus-Ph | AUROC | +0.240 | [+0.202, +0.279] | Yes |
| Caduceus-Ph | AUPRC | +0.409 | [+0.365, +0.456] | Yes |
| modernGENA | MCC | +0.455 | [+0.390, +0.515] | Yes |
| modernGENA | AUROC | +0.239 | [+0.204, +0.276] | Yes |
| modernGENA | AUPRC | +0.420 | [+0.370, +0.466] | Yes |

#### Panel B. Frozen Evo2 versus native impact scores (different evaluation regimes)

| Comparator | Metric | Evo2 – comparator | 95% bootstrap CI | Excludes zero |
|---|---|---|---|---|
| GPN-Star V100 | MCC | −0.286 | [−0.332, −0.244] | Yes |
| GPN-Star V100 | AUROC | −0.138 | [−0.166, −0.112] | Yes |
| GPN-Star V100 | AUPRC | −0.194 | [−0.227, −0.161] | Yes |
| AlphaGenome AVI | MCC | −0.297 | [−0.342, −0.253] | Yes |
| AlphaGenome AVI | AUROC | −0.139 | [−0.167, −0.111] | Yes |
| AlphaGenome AVI | AUPRC | −0.198 | [−0.234, −0.162] | Yes |

Panel B compares frozen Evo2 readouts with native scores evaluated under a different gene-excluded protocol. These comparisons cannot isolate the effects of architecture or biological input information. A separate direct comparison between AVI and GPN-Star V100 on the primary fixed-test scores is reported in Supplementary Table S4, Panel B.

#### Supplementary Table S12. Native impact-score thresholds selected using nine-gene training data

For each evaluation gene, native-score decision thresholds were selected by maximizing MCC on labelled training variants from the remaining nine genes, with ties resolved in favor of the higher threshold. The continuous scores remained unchanged. GPN-Star V100 used the sign-reversed mutation-rate-calibrated log-likelihood ratio, while AlphaGenome AVI used its PHRED-scaled composite score. Their numerical thresholds operate on different scales.

#### GPN-Star V100

| Held-out gene | Threshold | Train N | Train pos. | Train MCC | Test N | Test pos. | Test MCC | AUROC | AUPRC |
|---|---|---|---|---|---|---|---|---|---|
| ANO5 | 5.767 | 4481 | 1089 | 0.931 | 164 | 33 | 0.903 | 0.986 | 0.977 |
| CAPN3 | 5.767 | 4146 | 960 | 0.931 | 295 | 88 | 0.951 | 0.995 | 0.993 |
| DNAJB6 | 5.338 | 4714 | 1172 | 0.937 | 65 | 2 | 1.000 | 1.000 | 1.000 |
| DYSF | 4.893 | 3105 | 812 | 0.937 | 768 | 166 | 0.921 | 0.993 | 0.984 |
| FKRP | 5.338 | 4483 | 1090 | 0.931 | 172 | 26 | 1.000 | 1.000 | 1.000 |
| LMNA | 5.338 | 4203 | 952 | 0.931 | 284 | 117 | 0.942 | 0.989 | 0.990 |
| SGCA | 5.77 | 4556 | 1107 | 0.942 | 144 | 40 | 0.842 | 0.994 | 0.985 |
| SGCB | 5.767 | 4669 | 1150 | 0.936 | 83 | 20 | 0.834 | 0.961 | 0.956 |
| SGCG | 5.767 | 4661 | 1141 | 0.933 | 84 | 11 | 1.000 | 1.000 | 1.000 |
| TCAP | 5.767 | 4776 | 1174 | 0.934 | 35 | 6 | 0.898 | 0.989 | 0.958 |

#### AlphaGenome AVI

| Held-out gene | Threshold | Train N | Train pos. | Train MCC | Test N | Test pos. | Test MCC | AUROC | AUPRC |
|---|---|---|---|---|---|---|---|---|---|
| ANO5 | 23.87662 | 4481 | 1089 | 0.941 | 164 | 33 | 0.962 | 1.000 | 1.000 |
| CAPN3 | 23.87662 | 4146 | 960 | 0.952 | 295 | 88 | 0.853 | 0.956 | 0.957 |
| DNAJB6 | 23.87662 | 4714 | 1172 | 0.945 | 65 | 2 | 1.000 | 1.000 | 1.000 |
| DYSF | 23.964919 | 3105 | 812 | 0.942 | 768 | 166 | 0.958 | 0.999 | 0.996 |
| FKRP | 23.87662 | 4483 | 1090 | 0.946 | 172 | 26 | 0.977 | 1.000 | 1.000 |
| LMNA | 23.87662 | 4203 | 952 | 0.939 | 284 | 117 | 0.964 | 0.997 | 0.996 |
| SGCA | 23.87662 | 4556 | 1107 | 0.955 | 144 | 40 | 0.827 | 0.980 | 0.968 |
| SGCB | 23.87662 | 4669 | 1150 | 0.944 | 83 | 20 | 1.000 | 1.000 | 1.000 |
| SGCG | 23.87662 | 4661 | 1141 | 0.944 | 84 | 11 | 1.000 | 1.000 | 1.000 |
| TCAP | 23.964919 | 4776 | 1174 | 0.945 | 35 | 6 | 1.000 | 1.000 | 1.000 |

Additional archived records preserve the unrounded numerical values, number of threshold candidates, number of training variants predicted positive at the selected threshold, and the threshold-selection rule.

#### Supplementary Table S13. Frozen-readout hyperparameter selection in the leave-one-gene-out analysis

For each sequence model and held-out gene, StandardScaler plus L2 logistic regression was fitted using variants from the other nine genes in the canonical training partition. The candidate regularization grid was C ∈ {0.001, 0.01, 0.1, 1, 10, 100}; selection used five-fold locus-grouped cross-validation and MCC at threshold 0.5. Smaller C was preferred in a tie. Frozen foundation-model parameters were unchanged.

| Model | Held-out gene | Training N | Selected C | CV mean MCC | Final fit iterations |
|---|---|---|---|---|---|
| DNABERT-2 | ANO5 | 4481 | 0.1 | 0.133 | 105 |
| DNABERT-2 | CAPN3 | 4146 | 100 | 0.141 | 162 |
| DNABERT-2 | DNAJB6 | 4714 | 0.1 | 0.124 | 89 |
| DNABERT-2 | DYSF | 3105 | 0.1 | 0.101 | 91 |
| DNABERT-2 | FKRP | 4483 | 1 | 0.133 | 120 |
| DNABERT-2 | LMNA | 4203 | 100 | 0.115 | 136 |
| DNABERT-2 | SGCA | 4556 | 1 | 0.122 | 151 |
| DNABERT-2 | SGCB | 4669 | 100 | 0.135 | 150 |
| DNABERT-2 | SGCG | 4661 | 0.1 | 0.120 | 91 |
| DNABERT-2 | TCAP | 4776 | 100 | 0.126 | 135 |
| NT-v2 | ANO5 | 4481 | 0.01 | 0.270 | 51 |
| NT-v2 | CAPN3 | 4146 | 0.01 | 0.249 | 51 |
| NT-v2 | DNAJB6 | 4714 | 0.01 | 0.257 | 54 |
| NT-v2 | DYSF | 3105 | 0.01 | 0.188 | 44 |
| NT-v2 | FKRP | 4483 | 0.01 | 0.215 | 45 |
| NT-v2 | LMNA | 4203 | 0.01 | 0.231 | 56 |
| NT-v2 | SGCA | 4556 | 0.01 | 0.257 | 51 |
| NT-v2 | SGCB | 4669 | 0.01 | 0.248 | 48 |
| NT-v2 | SGCG | 4661 | 0.01 | 0.237 | 51 |
| NT-v2 | TCAP | 4776 | 0.01 | 0.244 | 57 |
| HyenaDNA | ANO5 | 4481 | 1 | 0.195 | 234 |
| HyenaDNA | CAPN3 | 4146 | 1 | 0.186 | 237 |
| HyenaDNA | DNAJB6 | 4714 | 100 | 0.187 | 287 |
| HyenaDNA | DYSF | 3105 | 1 | 0.149 | 231 |
| HyenaDNA | FKRP | 4483 | 100 | 0.198 | 267 |
| HyenaDNA | LMNA | 4203 | 1 | 0.193 | 222 |
| HyenaDNA | SGCA | 4556 | 10 | 0.187 | 278 |
| HyenaDNA | SGCB | 4669 | 100 | 0.231 | 285 |
| HyenaDNA | SGCG | 4661 | 1 | 0.215 | 233 |
| HyenaDNA | TCAP | 4776 | 10 | 0.190 | 263 |
| Caduceus-Ph | ANO5 | 4481 | 0.1 | 0.188 | 139 |
| Caduceus-Ph | CAPN3 | 4146 | 0.1 | 0.170 | 125 |
| Caduceus-Ph | DNAJB6 | 4714 | 100 | 0.186 | 433 |
| Caduceus-Ph | DYSF | 3105 | 1 | 0.197 | 229 |
| Caduceus-Ph | FKRP | 4483 | 1 | 0.157 | 267 |
| Caduceus-Ph | LMNA | 4203 | 10 | 0.158 | 363 |
| Caduceus-Ph | SGCA | 4556 | 0.1 | 0.177 | 132 |
| Caduceus-Ph | SGCB | 4669 | 100 | 0.175 | 327 |
| Caduceus-Ph | SGCG | 4661 | 10 | 0.202 | 362 |
| Caduceus-Ph | TCAP | 4776 | 10 | 0.193 | 346 |
| modernGENA | ANO5 | 4481 | 0.01 | 0.259 | 39 |
| modernGENA | CAPN3 | 4146 | 0.01 | 0.257 | 36 |
| modernGENA | DNAJB6 | 4714 | 0.01 | 0.254 | 38 |
| modernGENA | DYSF | 3105 | 0.01 | 0.227 | 35 |

| Model | Held-out gene | Training N | Selected C | CV mean MCC | Final fit iterations |
|---|---|---|---|---|---|
| modernGENA | FKRP | 4483 | 0.01 | 0.260 | 37 |
| modernGENA | LMNA | 4203 | 0.01 | 0.232 | 37 |
| modernGENA | SGCA | 4556 | 0.01 | 0.240 | 38 |
| modernGENA | SGCB | 4669 | 0.01 | 0.260 | 38 |
| modernGENA | SGCG | 4661 | 0.01 | 0.244 | 42 |
| modernGENA | TCAP | 4776 | 0.01 | 0.255 | 40 |
| Evo2 | ANO5 | 4481 | 0.1 | 0.641 | 380 |
| Evo2 | CAPN3 | 4146 | 0.1 | 0.643 | 354 |
| Evo2 | DNAJB6 | 4714 | 0.1 | 0.645 | 371 |
| Evo2 | DYSF | 3105 | 0.1 | 0.641 | 323 |
| Evo2 | FKRP | 4483 | 1 | 0.654 | 1012 |
| Evo2 | LMNA | 4203 | 1 | 0.620 | 961 |
| Evo2 | SGCA | 4556 | 0.1 | 0.662 | 369 |
| Evo2 | SGCB | 4669 | 0.1 | 0.652 | 359 |
| Evo2 | SGCG | 4661 | 1 | 0.646 | 999 |
| Evo2 | TCAP | 4776 | 0.1 | 0.651 | 369 |

Convergence iteration counts are retained as optimization diagnostics and do not measure predictive performance. Checkpoint identifiers and available software provenance are documented in Supplementary Methods S1.2.1. Native scoring systems are excluded because their evaluation used existing continuous scores without fitting logistic-regression readouts.

#### Supplementary Table S14. Exploratory HGVS-derived annotation-category analysis

##### Panel A. Category-level class composition across the canonical benchmark and fixed partitions

| HGVS-derived category | Full N− | Full N+ | Train N− | Train N+ | Test N− | Test N+ |
|---|---|---|---|---|---|---|
| Amino-acid substitution HGVS pattern | 114 | 696 | 81 | 462 | 33 | 234 |
| Intronic HGVS pattern | 2187 | 463 | 1525 | 347 | 662 | 116 |
| Stop-gained HGVS pattern | 0 | 521 | 0 | 365 | 0 | 156 |
| Synonymous HGVS pattern | 2807 | 12 | 1968 | 9 | 839 | 3 |
| UTR HGVS pattern | 151 | 0 | 103 | 0 | 48 | 0 |
| Unclassified HGVS pattern | 9 | 0 | 6 | 0 | 3 | 0 |
| Total | 5268 | 1692 | 3683 | 1183 | 1585 | 509 |

##### Panel B. Classification performance on the fixed 2,094-variant test set

| Evaluation | MCC | AUROC | AUPRC | Balanced accuracy | TN / FP / FN / TP |
|---|---|---|---|---|---|
| Fixed-test baseline | 0.796 | 0.939 | 0.828 | 0.873 | 1552/33/119/390 |
| Gene-excluded baseline | 0.796 | 0.927 | 0.842 | 0.873 | 1552/33/119/390 |

##### Methods and interpretation

Six mutually exclusive categories were assigned using regular-expression patterns in the ClinVar HGVS-like Name field, with precedence given to stop-gained, synonymous, amino-acid substitution, UTR, intronic and unclassified. UTR-prefixed intronic descriptions were assigned to UTR. These categories represent naming patterns without independent molecular-consequence validation. In Panel A, N− and N+ denote the aggregate ClinVar negative and positive classes.

For the fixed-test baseline, category-specific positive-class frequencies estimated from the 4,866 training variants were assigned as continuous scores to the corresponding test variants. Binary predictions used a threshold of 0.5. The gene-excluded baseline repeated this procedure using training variants from the other nine genes for each evaluation gene. Predictions were pooled across the same 2,094 test variants. AUPRC was calculated as average precision.

These exploratory controls demonstrate substantial associations between HGVS naming patterns and benchmark labels. They use different input information from the foundation models and native scoring systems. Their performance does not establish molecular causality, clinical diagnostic utility or generalization to independent cohorts.

#### Supplementary Table S15. Exploratory HGVS-category-stratified discrimination

This retrospective analysis evaluates 12 frozen/adapted 101-bp sequence-model configurations and the two native scoring systems on identical canonical test variants. All original continuous scores, 0.5 sequence-model decision thresholds and training-selected native thresholds were preserved. Model-specific metrics were recalculated only within each HGVS-derived annotation category; no refitting or subgroup threshold optimization was performed. The category definitions and fixed/gene-excluded prevalence-only baseline are given in Supplementary Table S14.

##### Panel A. Test-set category composition

| HGVS text category | N test | Positive | Negative | Positive fraction | Both classes ≥20 |
|---|---|---|---|---|---|
| Amino-acid substitution | 267 | 234 | 33 | 87.6% | Yes |
| Intronic | 778 | 116 | 662 | 14.9% | Yes |
| Synonymous | 842 | 3 | 839 | 0.4% | No |
| Stop-gained | 156 | 156 | 0 | 100.0% | No |
| UTR | 48 | 0 | 48 | 0.0% | No |
| Unclassified | 3 | 0 | 3 | 0.0% | No |

Categories are mutually exclusive text-pattern groupings based on the source ClinVar HGVS Name field, not independently validated variant-consequence assignments. AUROC and AUPRC are undefined when only one class is present. The synonymous category contains only three positive test variants and is not used for comparative interpretation.

##### Panel B. Classification metrics in two mixed-class categories

| Model | Regime | Amino MCC | Amino AUROC | Amino AUPRC | Intronic MCC | Intronic AUROC | Intronic AUPRC |
|---|---|---|---|---|---|---|---|
| DNABERT-2 | frozen | 0.112 | 0.564 | 0.909 | 0.053 | 0.556 | 0.176 |
| DNABERT-2 | adapted | 0.000 | 0.534 | 0.891 | 0.218 | 0.750 | 0.482 |
| NT-v2 | frozen | 0.136 | 0.569 | 0.915 | 0.245 | 0.678 | 0.344 |
| NT-v2 | adapted | 0.195 | 0.675 | 0.941 | 0.314 | 0.868 | 0.735 |
| HyenaDNA | frozen | 0.054 | 0.469 | 0.889 | 0.275 | 0.724 | 0.346 |
| HyenaDNA | adapted | 0.012 | 0.505 | 0.884 | 0.331 | 0.815 | 0.528 |
| Caduceus-Ph | frozen | 0.075 | 0.566 | 0.908 | 0.033 | 0.620 | 0.208 |
| Caduceus-Ph | adapted | 0.039 | 0.593 | 0.922 | 0.156 | 0.648 | 0.269 |
| modernGENA | frozen | 0.091 | 0.601 | 0.916 | 0.326 | 0.735 | 0.375 |
| modernGENA | adapted | 0.011 | 0.533 | 0.886 | 0.104 | 0.613 | 0.274 |
| Evo2 | frozen | 0.171 | 0.700 | 0.940 | 0.613 | 0.856 | 0.735 |
| Evo2 | adapted | 0.208 | 0.681 | 0.933 | 0.302 | 0.813 | 0.653 |
| GPN-Star V100 | native | 0.495 | 0.889 | 0.981 | 0.928 | 0.986 | 0.972 |
| AlphaGenome Atlas AVI | native | 0.544 | 0.923 | 0.988 | 0.980 | 1.000 | 0.998 |

The amino-acid substitution category contained 267 variants (234 positive, 33 negative), while the intronic category contained 778 (116 positive, 662 negative). AUROC and AUPRC were calculated from unchanged continuous scores, with AUPRC reported as average precision. MCC used the original classification thresholds: 0.5 for sequence models, 5.767 for GPN-Star V100 and 23.87662025 for AVI PHRED. No category-specific threshold optimization was performed. AUPRC values for amino-acid substitutions should be interpreted in light of the 87.6% positive-class prevalence. Comparisons between native scores and sequence-model readouts remain descriptive because their underlying information sources differ.

##### Panel C. Within-intronic HGVS position offsets and label composition

| Absolute cDNA offset from exon junction | Positive | Negative | Total |
|---|---|---|---|
| 1–2 bases | 104 | 0 | 104 |
| 3–6 bases | 7 | 44 | 51 |

| Absolute cDNA offset from exon junction | Positive | Negative | Total |
|---|---|---|---|
| 7–20 bases | 3 | 441 | 444 |
| 21–100 bases | 0 | 53 | 53 |
| >100 bases | 2 | 124 | 126 |

Among 116 positive-class variants in the intronic HGVS category, 104 (89.7%) had text-inferred cDNA offsets of ±1 or ±2, compared with none of the 662 negative-class variants. The remaining intronic subset contained 12 positive and 662 negative variants. This strong association between splice-adjacent naming patterns and clinical-significance labels may contribute to the high discrimination observed for native scores within the broad intronic category. The offsets were inferred from HGVS text without independent molecular-consequence validation.

#### Panel D. Exploratory paired genomic-locus bootstrap uncertainty

| HGVS group | Score or paired AUROC difference | Estimate | 95% bootstrap percentile interval |
|---|---|---|---|
| Amino acid | Evo2 frozen (101 bp) | 0.700 | [0.605, 0.789] |
| Amino acid | GPN-Star V100 | 0.889 | [0.823, 0.946] |
| Amino acid | AlphaGenome AVI | 0.923 | [0.874, 0.960] |
| Amino acid | AVI minus GPN-Star | 0.034 | [-0.008, 0.083] |
| Amino acid | GPN-Star minus Evo2 | 0.189 | [0.103, 0.279] |
| Amino acid | AVI minus Evo2 | 0.223 | [0.145, 0.306] |
| Intronic | Evo2 frozen (101 bp) | 0.856 | [0.795, 0.911] |
| Intronic | GPN-Star V100 | 0.986 | [0.969, 0.998] |
| Intronic | AlphaGenome AVI | 1.000 | [0.999, 1.000] |
| Intronic | AVI minus GPN-Star | 0.014 | [0.002, 0.030] |
| Intronic | GPN-Star minus Evo2 | 0.130 | [0.075, 0.188] |
| Intronic | AVI minus Evo2 | 0.144 | [0.089, 0.205] |

Exploratory paired genomic-locus bootstrap intervals were calculated using 2,000 replicates (seed 42), with loci resampled within each category and identical resampled indices used across scores. The reported 95% percentile intervals are descriptive and were not adjusted for multiple comparisons. They do not account for upstream score-development data, sampling additional genes or refitting downstream classifiers. Differences between native scores and sequence readouts cannot be attributed to individual architectural or biological information sources.

#### Reproducibility and interpretation

All 2,094 test VariationIDs were matched against the canonical ClinVar records, with identical labels and genomic coordinates. The twelve frozen and adapted 101-bp sequence-model scores and classifications agreed with the archived fixed-test sequence-model predictions, while GPN-Star scores agreed with the archived native-score predictions. Overall performance and category counts were consistent with the archived benchmark and Supplementary Table S14. Full-precision metrics, paired bootstrap intervals and matched variant-level scores are supplied as machine-readable evidence. This analysis includes only the original 101-bp sequence configurations, not the 201- or 501-bp predictions.
